# Subclonal mutation load predicts survival and response to immunotherapy in cancers with low to moderate tumor mutation burden

**DOI:** 10.1101/2024.07.03.601939

**Authors:** Yujie Jiang, Matthew D Montierth, Kaixian Yu, Shuangxi Ji, Quang Tran, Xiaoqian Liu, Jessica C. Lal, Shuai Guo, Aaron Wu, Yaoyi Dai, Seung Jun Shin, Ruonan Li, Shaolong Cao, Yuxin Tang, Tom Lesluyes, Scott Kopetz, Pavlos Msaouel, Anil K. Sood, Ginny Devonshire, Christopher M. Jones, Jaffer Ajani, Sumit K Subudhi, Ana Aparicio, Padmanee Sharma, John Paul Shen, Marek Kimmel, Jennifer R. Wang, Maxime Tarabichi, Rebecca C. Fitzgerald, Hongtu Zhu, Peter Van Loo, Wenyi Wang

## Abstract

Intra-tumor heterogeneity is characterized by a diverse population of tumor clones and subclones which are important drivers of tumor evolution and therapeutic response. However, accurate subclonal reconstruction at scale remains challenging. We developed a machine learning tool, CliPP, and surveyed 9,972 tumors from 32 cancer types. We found that high subclonal mutation load (sML), the fraction of subclonal single nucleotide variants (SNVs) to all SNVs in the coding region, was prognostic of survival (progression free survival or overall survival) in 18 cancer types. In 14 cancers with low to moderate tumor mutation burden (TMB), high sML was associated with better prognosis. In immunotherapy trials for 42 metastatic prostate cancer (mCRPC), high sML was predictive of favorable response to ipilimumab and associated with increased CD8^+^ T-cell infiltration and decreased macrophage population. A validation using 613 whole-genomes of esophageal adenocarcinoma confirms the favorable effect of high sML and the observed tumor-associated macrophage. Our study identifies sML as a key feature of cancer, suggesting a biphasic relationship between evolutionary dynamics and differential immune environments. Finally, sML may serve as an orthogonal approach to identify likely responders of immune checkpoint blockade in low to moderate TMB tumors.

## Introduction

Mutations in tumor cells accumulate over time, originating from a single cancer-initiating cell and progressing to the diverse mutations present across a mature tumor^1,2^. This process leads to the development of heterogeneous subpopulations of cells, i.e., (sub)clones, with unique genetic profiles from the ancestral cancer cell^3–5^ (**Fig. 1a**). Over the past decade, substantial improvement in understanding of the temporal and spatial hallmarks of mutation events across tumor clones^6–10^, with many supported by pan-cancer investigations^11–15^, has created new opportunities for the development of cancer therapy.

**Figure 1:**
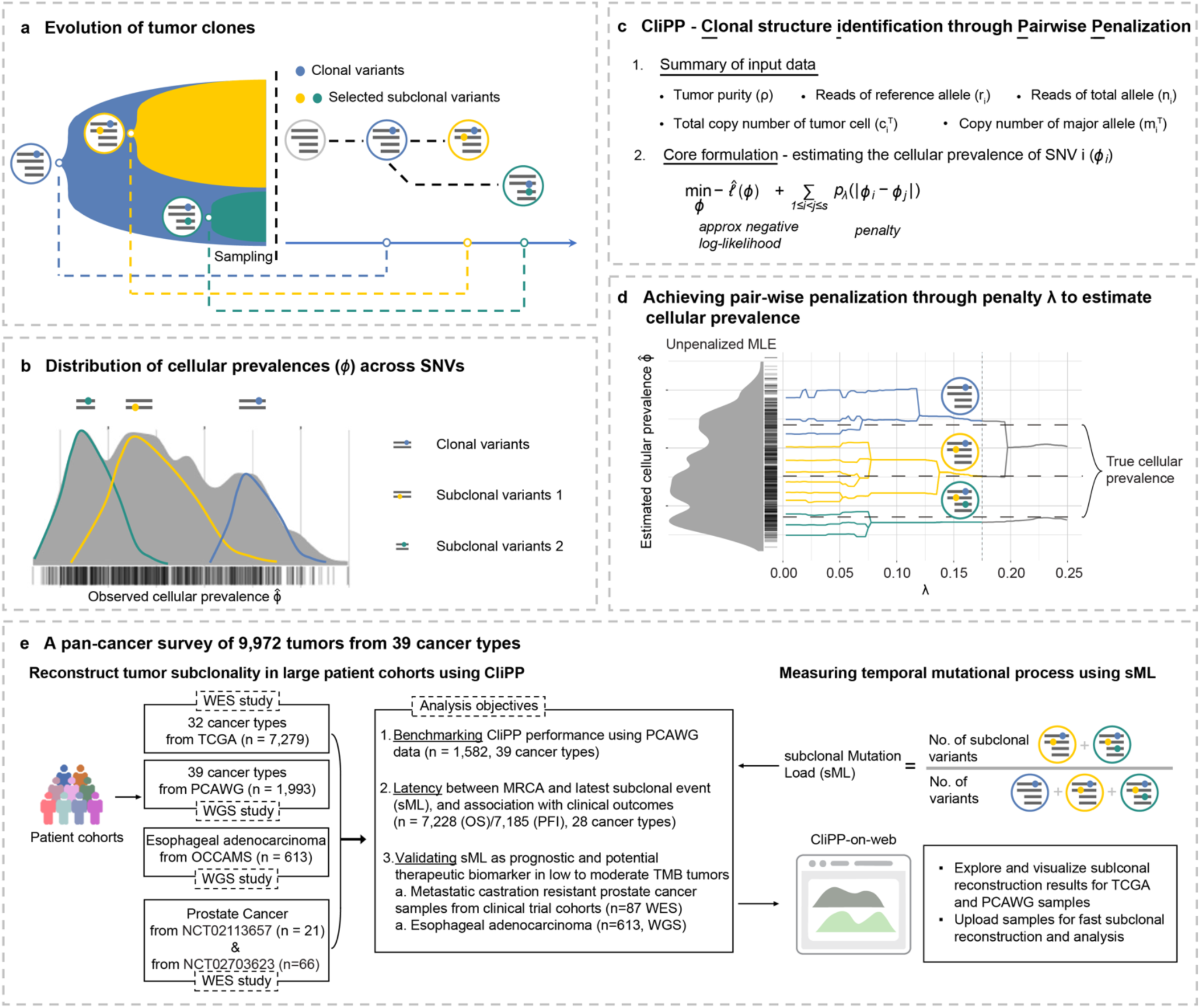
Subclonal reconstruction and CliPP model overview. **(a)** A visual representation of tumor evolution with genotype circles indicating representative somatic SNVs. Colored dots represent SNVs. **(b)** Cellular prevalence (CP) distribution of SNVs (each SNV is a bar along the X axis) for an illustrative example sample, characterized by one clonal cluster (blue) and two subclonal clusters (yellow and green). **(c)** Input data and the core optimization problem for the CliPP model, a novel computational tool for subclonal reconstruction. **(d)** CliPP infers clusters of SNVs and CP values using a penalty function that varies over a penalty parameter, λ, which is shown along the x-axis. The y-axis corresponds to CP estimates (0 to 1) connecting the nearest CP estimates over 100 evenly spaced λ values (0.01 to 0.25). The vertical dotted line indicates the λ corresponding to the identified optimal solution (see **Methods**). Somatic SNV clusters distinguish clonal cancerous cells (blue) from new subclonal cancerous cell populations (yellow and green) that converge for homogeneity pursuit in CPs. **(e)** Objectives for utilizing the CliPP-based tumor subclonal reconstruction results from 9,972 tumor samples across 39 cancer types from multiple cohorts.

Early pan-cancer studies identified a strong association of total tumor mutation load with patient outcomes^14,16^, and later with immunotherapy response^17,18^. In 2020, the US Food and Drug Administration (FDA) approved the first histology-agnostic programmed cell death (PD-1) therapy for patients with solid tumors exhibiting a tumor mutation burden (TMB, calculated as the number of mutations per million bases) of at least 10 (high-TMB)^19^. Among high-TMB cancers, clonal TMB was the strongest predictor of immune checkpoint blockade (ICB) response, while subclonal TMB showed no significant association^20,21^. These studies have paved a promising path to assess simultaneously how selection shapes mutation patterns as well as patient survival^14^. Still, more work is required to adopt a robust biomarker of ICB response that is inclusive of low to moderate TMB (TMB < 10 mut/Mb)^22^.

Recent pan-cancer studies have demonstrated distinct evolutionary paths and immune environments between cancers with high versus low mutational load. Negative selection appears stronger in low mutational load cancers^23^, while high mutational load cancers uniquely exhibit focal loss of heterozygosity of HLA-I as a mechanism of immune evasion^24^. With ∼90% (6,341 out of 7,279 studied) of The Cancer Genome Atlas (TCGA) tumors having a TMB < 10 mut/Mb, evaluating the clinical impact of the subclonal structures across low, moderate, and high TMB cancers is paramount. However, most large-scale efforts to reconstruct pan-cancer subclonal structures lack annotation of clinical outcomes^12^. While smaller studies with paired genetic and clinical annotations have provided valuable insights^25–29^, scaling translational studies of tumor evolutionary dynamics remains challenging. This is due to the high computational burden, inter and intra-tumor heterogeneity, and the need for manual curations of the per-sample mutation data for subclonal reconstruction^12^. These challenges highlight the need for scalable, efficient, and accurate subclonal reconstruction methods to advance cancer evolution research.

In this study, we have developed a novel scalable method for subclonal reconstruction that overcomes these challenges by 1) performing Clonal structure identification through Pairwise Penalization (CliPP) to improve the computational speed per sample by 100-1,000 fold, compared to existing methods^30–32^; 2) benchmarking accuracy using 1,582 WGS, and 447 matched whole-exome sequencing (WES)/whole-genome sequencing (WGS) patient samples; and 3) generating pan-cancer subclonal structures of pre-treatment tumors with clinical follow-up data of at least 5 years. We evaluated the clinical impact of subclonal mutational load (sML) in 7,279 WES patient samples in TCGA, validating findings in 42 WES metastatic castration-resistant prostate cancer (mCRPC) samples treated with ipilimumab^33,34^ and 613 WGS esophageal adenocarcinoma samples with clinical follow-up from the Oesophageal Cancer Clinical And Molecular Stratification (OCCAMS) consortium^35–37^. We find that sML is an essential feature of cancer and may serve as a robust prognostic and therapeutic biomarker, particularly in low to moderate TMB cancers. Finally, we provide a Graphic User Interface tool, CliPP-on-web, for interactive real-time subclonal reconstruction, freely available at: https://bioinformatics.mdanderson.org/apps/CliPP.

## Results

### Overview of subclonal reconstruction by CliPP

For a given tumor sample, the sequencing-based variant allele frequency (VAF) spectrum (**Fig. 1b**) exhibits a mixture of multiple peaks corresponding to varying cellular prevalence (CP) of single-nucleotide variants (SNVs). Subclonal reconstruction refers to a procedure to identify cancer cell clones through clustering of variant read counts, adjusting for allele copies and tumor purity, to identify groups of variants with similar CPs. Available tools employ a Bayesian modeling framework^31,32,38,39^, which comes with a high computational cost, particularly for running WGS data. This challenge was recently alleviated by a variational Bayes^30,40^ approach, yet there remains a large margin to be minimized when processing tens of thousands of tumor samples. We, therefore, propose an orthogonal approach to existing methods called <u>Cl</u>onal structure identification through <u>P</u>airwise <u>P</u>enalization (CliPP), which is rooted in the domain of regularized regression in machine learning. Imposing a pairwise penalty in the form of smoothly clipped absolute deviation (SCAD)^41^ to the parameter (e.g., CP) estimation per data point (e.g., mutation) is, in general terms, a modeling advancement over the fused LASSO regression^42^ and a specialized extension of the recently proposed CARDS algorithm^43^, which searches for homogeneity in regression parameters. Consequently, in this novel application of the penalized likelihood statistical framework to subclonal reconstruction, our resulting sparse and homogeneous pattern of CPs across mutations corresponds to a clustering procedure (**Fig. 1c-d**, **Methods**). The output includes the clustered mutations along with their associated homogenized CP values. CliPP offers exceptional computational efficiency by reformulating the subclonal reconstruction problem as a constrained optimization problem that can be efficiently solved by existing optimization techniques (see **Methods**). This fast tool enabled our pan-cancer study across >5,000 simulated samples and >9,900 sequenced patient samples from >30 cancer types to identify and validate clinically relevant features (see **Fig. 1e** for an overview and **Extended Data** Fig. 1 for a consort diagram of the study).

In terms of clustering accuracy, using three simulation datasets (total n=5,515) covering a wide range of values for key features that are known to influence accuracy^12^ including tumor purity, percent of genome with copy number alterations, sequencing read depth, and the number of mutation clusters, CliPP performed similarly to PhyloWGS^31^, PhylogicNDT^32^, and PyCloneVI^30^ (**Methods**, **Extended Data** Fig. 2a-d**, Supplementary Information Section 1**). Using WGS data from 1,582 patient tumor samples in the Pan Cancer Analysis of Whole Genomes (PCAWG) study, CliPP ranked the second highest concordance to the consensus among all 11 methods^12,30^ (concordance correlation coefficient = 0.97, **Fig. 2a, Supplementary Information Section 1.4**, **Methods**), confirming its high accuracy.

**Figure 2:**
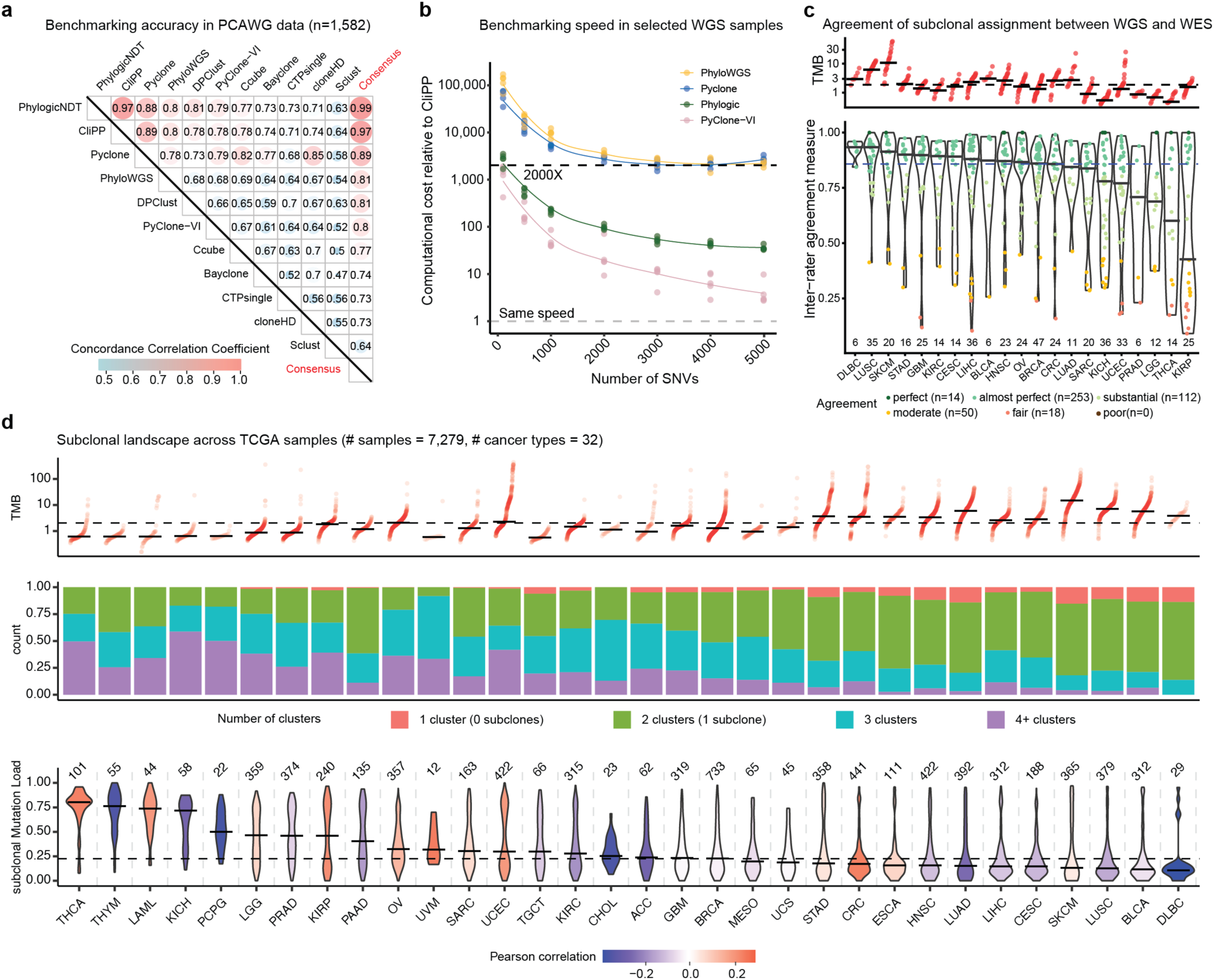
CliPP benchmarking and its pan-cancer application in TCGA. **(a-b).** Benchmarking accuracy and speed using PCAWG WGS data. **(a)** Pairwise Concordance Correlation Coefficient (CCC) heatmap of the estimated fraction of clonal mutations by 11 reconstruction methods and their consensus on the PCAWG dataset (n=1,582). **(b)** Computational time comparison in fold-change relative to CliPP on PCAWG samples with up to 5,000 SNVs. Each dot represents one of the 35 samples where all five methods were applied. **(c)** Violin plots of inter-rater agreement of subclonal mutations called by CliPP from WES versus WGS. Points represent tumor samples, and are colored by agreement category. Cancer types are ordered by median agreement scores (black bar), with the median across all samples indicated by the dashed line. Numbers at the bottom correspond to the total number of tumor samples for the corresponding cancer types. Cancer types with higher TMB (shown in red in the top panel) tend to have higher agreement. See **Extended Data** Figure 1b for the reference of cancer-type acronyms. **(d)** Distributions of the TMB, proportion of mutation clusters, and subclonal mutation load (sML) across 32 cancer types. Every column corresponds to one cancer type. The sigmoid plot depicts within cancer type distributions of TMB with the black horizontal bar indicating the median TMB, and the median TMB across all samples denoted by the dashed horizontal line. The stacked bar plots show the number of samples with a given number of clusters identified. The violin plots show the distribution of subclonal Mutation Load (sML) within each cancer type. The number of tumor samples for each cancer type is indicated above each violin plot. Violin plots color represents Pearson correlation coefficients of sML versus TMB using all samples within a given cancer type.

In terms of computational speed, CliPP finished analyzing 2,778 PCAWG samples within 16 hours and 9,654 TCGA samples in one hour, presenting at least a 2,000-fold improvement in speed as compared to PyClone^39^ and PhyloWGS^31^, and 4-1,000 fold improvement when compared to PyClone-VI^30^ (**Fig. 2b, Extended Data** Fig. 2e**, Methods**). WES data covers only 1-2% of the genome, and as such, most samples (75%) in TCGA present with less than 200 mutations. It is in samples with lower numbers of SNVs that CliPP has the largest speed advantage **(Fig. 2b)**. The overall speedup of CliPP over PyClone-VI for the entire TCGA is 514-fold (**Extended Data** Fig. 2f), and we project a speedup of ∼1,000 fold of CliPP versus PhylogicNDT in TCGA (**Extended Data** Fig. 2e**, Supplementary Information Section 2**).

In summary, the comparable accuracy and substantial speedup achieved by CliPP serve as a foundation for processing the large pan-cancer patient cohorts of WES data in TCGA. To further maximize access to our expedited and expanded pan-cancer survey, we have deployed CliPP-on-web, a Shiny app, where users with no prior computational experience can perform subclonal reconstruction and visualize the subclonal architecture (**Extended Data** Fig. 3).

### The subclonal landscape of 7,279 tumors

Using CliPP, we provide an updated portrait of the extent of genetic intra-tumor heterogeneity (ITH) across and within tumor types with a larger sample size in TCGA, as compared to the previous two pan-cancer studies with ∼1,200 tumors^13^ and ∼2,700 tumors^12^. As the first study at a much larger scale, we address the bottleneck question of whether WES can also provide sufficient signal for cancer evolution **(Extended Data** Fig. 4a). With 447 samples from 21 cancer types analyzed by CliPP on both WES and WGS data (profiled by TCGA and PCAWG, **Extended Data** Fig. 1a), we find CliPP-based subclonal reconstruction results are highly consistent between the two platforms, with a median inter-rater agreement (B-statistic, see **Methods**) of 0.86, indicating high agreement (**Fig. 2c**). Among those with disagreement, we found a trend in changing from WGS-clonal to WES-subclonal (**Methods**, 1,754 out of 1,907 (92%) mutations in **Extended Data** Fig. 4b), which may be explained by the higher read depths in WES (**Extended Data** Figs. 4c-d). Further differences in technical factors such as the input VAF distribution, purity estimate, and library preparation (14% were from different analytes) were also associated with lower agreement (**Methods**, **Supplementary Information Section 3**). These factors are independent causes from the mathematical process of subclonal reconstruction. Thus, we conclude it is appropriate, after proper pre-processing to alleviate the impact of the above factors, mainly the quality of copy number calling (see details in **Extended Data** Fig. 1a), to evaluate subclonality in TCGA WES data, hence taking advantage of an associated wealth of clinical annotations.

Across 7,279 tumors from 32 cancer types in TCGA that met our inclusion criteria (**Methods, Extended Data** Fig. 1a), we observed high consistency with the PCAWG study of ∼2,700 WGS tumors in the subclonal landscape across tumors, as well as across driver genes. As high as 94% (n=6,849) of TCGA samples present at least one subclonal cluster (**Fig. 2d**). Using a robustly annotated set of 8,586 driver mutations in 574 genes across 2,469 samples from 16 cancer types^44^, we find that 1,757 (20.5%) driver mutations were subclonal (**Extended Data** Fig. 5a, provided as a resource on CliPP-on-web). Overall, driver mutations were reported in 12% of subclonal mutation clusters in TCGA, in line with the 11% observed in PCAWG^12^ **(Extended Data** Fig. 5b, **Supplementary Information Section 4**). Utilizing the available clinical outcomes in TCGA, we find in six (out of 16 total) driver gene/cancer-type combinations, the presence of subclonal driver mutations is associated with differing prognosis: *PTEN, PIK3CA,* and *CTNNB1* in uterine corpus endometrial carcinoma (UCEC), *FAT1* in head and neck squamous cell carcinoma (HNSC), *IDH1* in low-grade glioma (LGG), and *BRAF* in thyroid papillary carcinoma (THCA) (**Supplementary Table 1**, **Extended Data** Fig. 5c-j). This novel finding highlights that an understanding of the clonality of driver mutations can provide clinicians with important information on which to guide treatment decisions.

Following TMB as a widely accepted clinical marker^19^ and the recent endeavors in further delineating the timing of mutations^11^, we quantify ITH via subclonal mutation load (sML), the proportion of subclonal mutations, merging mutations from multiple subclonal clusters (see **Methods**). We hypothesize that sML is a surrogate measure for the latent time between the most recent common ancestor and the latest subclonal expansion, and for the acceleration of mutation accumulation^11^. Consistent with the reported PCAWG results^12^, in TCGA across cancer types, median sML is moderately negatively correlated with median TMB (Pearson correlation = -0.50, *P*-value = 0.003, **Fig. 2d**) but not correlated with the technical factor median number of reads per chromosome copy (nrpcc^45^) (Pearson correlation = 0.28, *P*-value = 0.12). Within individual cancer types, sML is mostly weakly correlated with nrpcc, and not correlated with TMB (**Supplementary Information Section 5.1**, **Fig 2d**). Thyroid and thymus cancers, which have low median TMB, exhibit the highest sML values. Melanoma, lung squamous cancer, bladder cancer, and B-cell lymphoma, which have high TMB, exhibit the lowest sML values. The dynamic landscape of sML and TMB levels across cancer types suggests that sML may be biologically informative for cancers with low to moderate TMB.

### Subclonal mutation load (sML) is predictive of cancer prognosis

Across TCGA, we find that lower sML is significantly associated with clinical or pathological features/subtypes suggesting a poorer prognosis such as triple-negative breast cancer^48,49^, high Gleason score in prostate cancer^50^, smoking status, and low PD-L1 expression in lung cancer^51^, HPV negative in head and neck cancer^52^ and GBM classical subtype^53^ (**Fig. 3a, Methods**). In contrast, we observe no significant associations between sML and age, sex, TMB, clinical stage, or technical factors such as the size of the tumor biopsy or nrpcc (**Supplementary Information Section 5.1**). Together, our observations suggest sML may be a clinically relevant feature independent of TMB, stage, or other possible confounders.

**Figure 3:**
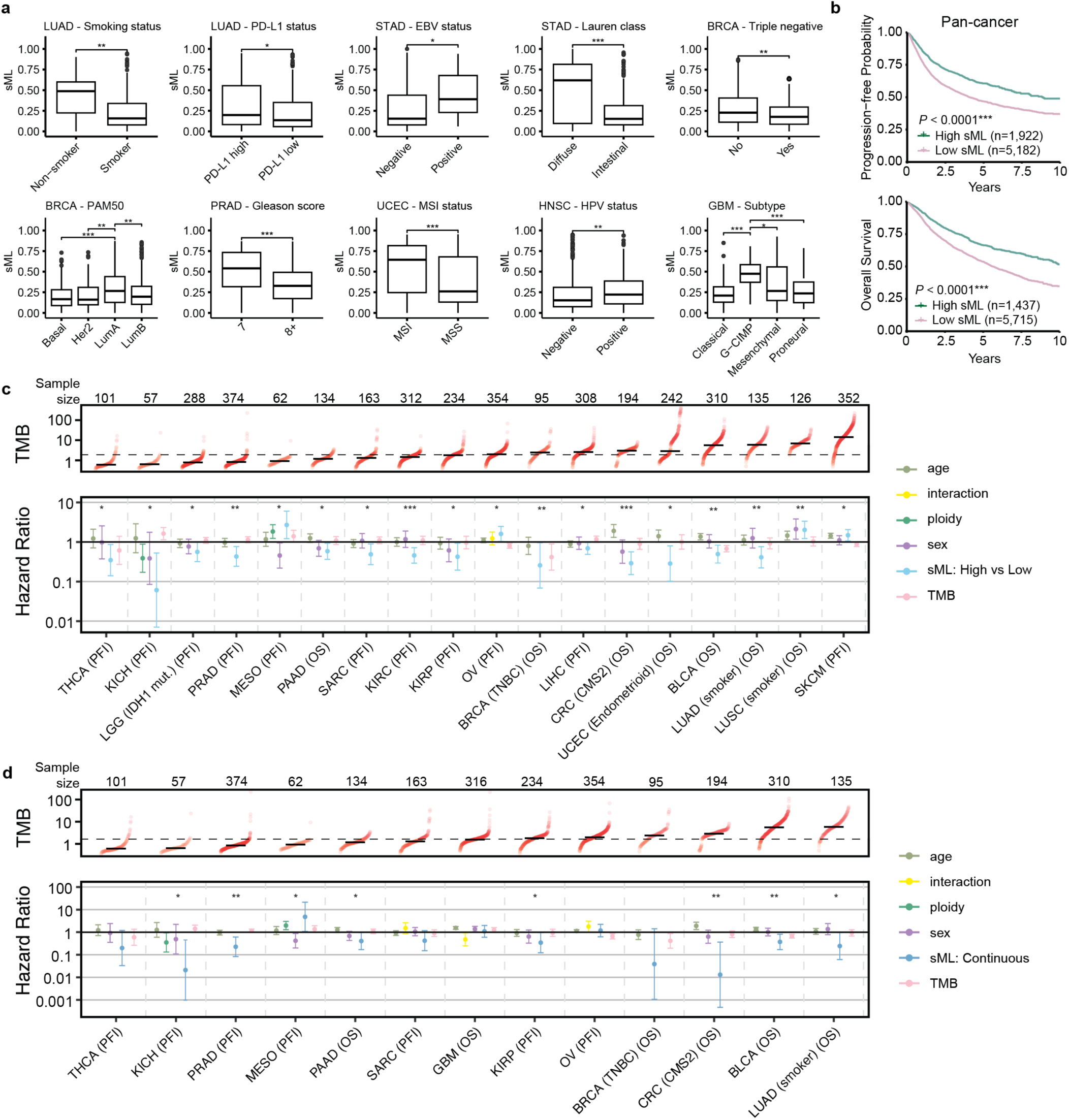
Biological and clinical associations of sML in TCGA. (**a**) Boxplots comparing sML between clinically utilized subtypes and cancer phenotypes. *P-*values for pairwise comparisons were calculated using the Wilcoxon rank-sum test. (**b**) Kaplan-Meier (KM) curves of progression-free interval (PFI, n=7,104) and overall survival (OS, n=7,152) for TCGA. Dark green lines indicate patients with high sML, and light purple lines indicate patients with low sML. Log-rank test *P*-values between high- and low-sML groups are shown. **(c)** Forest plot of hazard ratios (HRs) and 95% confidence intervals (CIs) from multivariate Cox proportional hazard models for Overall Survival (OS) or Progression-Free Interval (PFI) in TCGA for binarized (low/high) sML. Models are adjusted for age, TMB, and sex if applicable. Stepwise variable selection was performed with additional possible variables including ploidy, purity, coverage, and an interaction term between sML and TMB. The selected model is presented for each cancer type. \**P* < 0.05, \*\**P* < 0.01, \*\*\**P* < 0.001. Cancer types are ordered by median TMB, for which a sigmoid plot is shown on top. The dotted line in the TMB panel indicates the median value across all samples. The number of tumor samples for each cancer type is shown above each sigmoid plot. **(d)** Forest plot of HRs and 95% CIs of multivariate Cox proportional hazard models for OS or PFI in TCGA for continuous sML. Models are adjusted for age, TMB, and sex, if applicable. Stepwise variable selection was performed with additional possible variables, including ploidy, purity, coverage, and an interaction term between sML and TMB. The selected model is presented for each cancer type. Cancer types shown have either the main effect or the interaction effect of sML to be at least marginally statistically significant (*P* ≤ 0.1).

Using clinical outcomes (Overall Survival, OS and Progression Free Interval, PFI) in TCGA across 26 cancer types and their subtypes (see **Methods, Supplementary Table 2**), we find that high sML is associated with better OS/PFI at a pan-cancer level (**Fig. 3b**) within 14 cancer (sub)types, with an additional four cancer (sub)types demonstrating the opposite trend (**Methods**, **Fig. 3c, Extended Data** Fig. 6a-b**, Supplementary Tables 3**). These four cancer (sub)types were mesothelioma, ovarian cancer, lung squamous-smokers, and melanoma. Other than mesothelioma, three cancer types presented the highest levels of genome instability (**Extended Data** Fig. 6c) and/or the highest median TMB among all cancers (**Fig 3c**: top panel), which may explain the opposite trend. The predictive effects of sML remain unchanged after adjusting for potential technical confounders, including purity, ploidy, read depth, and nrpcc (**Supplementary Table 3**, **Supplementary Information Section 5.2**). The significant predictive effect of sML persisted as we moved from dichotomizing sML to using it as the continuous variable in a Cox model. Using variable selection, we find significant main effects of continuous sML in 10 cancer (sub)types and a significant interaction term of sML and TMB in three cancer types (**Methods**, **Fig. 3d, Supplementary Table 4**). Finally, replacing sML-based patient classification with TMB-based classification does not provide the same statistical significance (**Supplementary Information Section 5.2**), supporting the unique contribution of sML.

We also evaluated sML patient stratification with the addition of Shannon Index (SI)^26^, a diversity measure that increases when there are more mutation clusters and when mutations are more equally divided in each cluster. Given the collinear relationship between sML and SI (**Extended Data** Fig. 7a), SI contributes additional information in tumors with sML greater than 0.5, which represents the minority of TCGA (median sML = 0.23 in **Fig. 2d, Extended Data** Fig. 7b). Among those samples, a high SI indicates that subclonal mutations belong to multiple mutation clusters with distinct CPs, whereas a low SI indicates all mutations belong to the same or a small number of clusters. As these behaviors represent different evolutionary trajectories, we indeed find that the addition of SI partitions high sML tumors into further distinct survival groups of small numbers of patients in 11 cancer types (**Extended Data** Figs. 7c-d, **Supplementary Information Section 6**). In summary, with low and moderate TMB cancers, SI can sometimes complement sML, but overall, sML remains an easily interpretable and representative feature that encodes cancer evolution.

Recent findings reported that clonal TMB or clonal neoantigens can be more predictive of response to ICB but are limited in studying cohorts enriched with high TMB tumors^20,54^. In contrast, our pan-cancer findings in sML presenting two opposite directions of prognosis outcomes provide a new foundation for a further hypothesis that sML is predictive of ICB in the opposite direction in low to moderate TMB cancers compared to high-TMB cancers, i.e., in many cancer types, tumors with high sML will respond better to ICB.

### Subclonal mutation load is associated with immunotherapy response

mCRPC is a tumor type with low TMB despite which some patient subgroups benefit from immune checkpoint blockade. Biomarkers that can identify them remain to be defined^34^. We evaluated the effect of sML in two cohorts of mCRPC patients exposed to ipilimumab (IPI): monotherapy NCT02113657^55^ and DynAMo trial NCT02703623^33^ (**Fig. 4a**, see cohort demographics in **Supplementary Table 5**). In the IPI monotherapy trial, patients with high sML experienced improved radiographic/clinical PFS (rcPFS) in this trial (n=8 vs. 13, log rank *P*-value=0.005, **Fig. 4b**) and overall survival, with up to 8 years of follow-up (log rank *P*-value=0.03, **Extended Data** Fig. 8a) compared to those with low sML. The distribution of sML was not correlated with TMB, metastatic sites, or nrpcc, and further subset-specific analysis for prostate tissues only replicated the significant clinical outcome association and the direction of sML (**Supplementary Information Section 7.1**, **Methods**). In the DynAMo trial (**Supplementary Table 5**), patients with high sML also experienced longer failure-free survival (n=6 vs. 15, log rank *P*-value= 0.004, **Fig. 4c**, **Supplementary Table 6**, **Methods**), after receiving ARPi plus ipilimumab compared to those with low sML. For both datasets, using Cox regression with variable selection, we found the significant effect of sML remained after adjusting for TMB, age, PD-L1 density, and other technical confounders (**Fig. 4d, Supplementary Table 6, Supplementary Information Section 7.1, Methods**). In the DynAMo trial where additional mCRPC patients (n = 45) received ARPi alone or combined ARPi and chemotherapy (ARPi+CC), we observed that high sML is not significantly associated with outcomes, although still presenting a similar but weaker trend in the ARPi+CC arm (**Extended Data** Fig. 8b). This contrast supports an immune-specific activation that underlies tumors with high sML.

**Figure 4:**
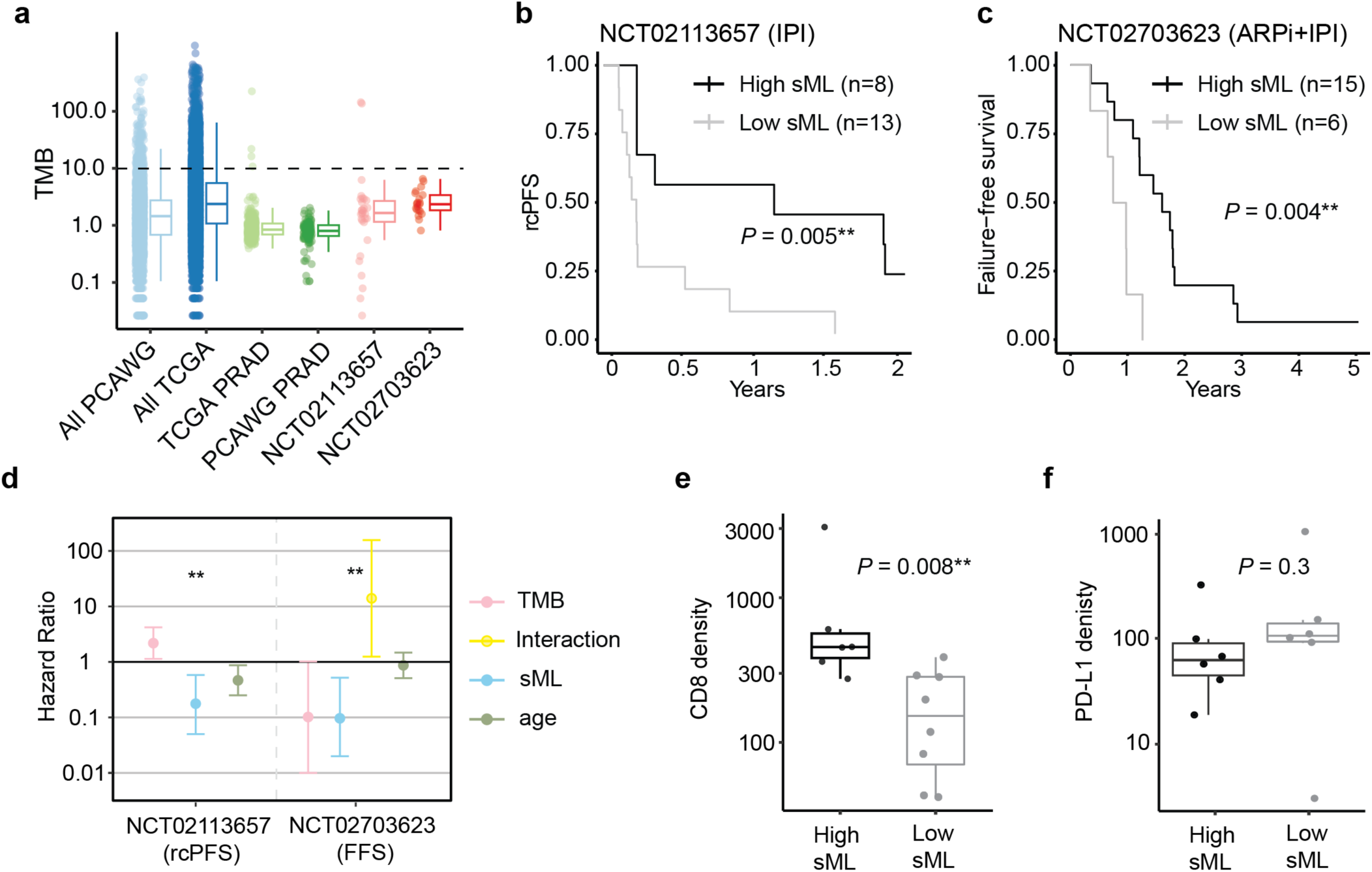
Association of sML with response to ICB in clinical trial cohorts. **(a)** Boxplots of TMB distributions for all PCAWG and TCGA samples, prostate cancer samples from PCAWG and TCGA, and for castration-resistant metastatic prostate cancer (mCRPC) samples from clinical trials NCT02113657 and NCT02703623. Dashed line represents the TMB=10 **(b)** Kaplan-Meier (KM) plot showing radiographic/clinical progression-free survival (rcPFS) in patients from the IPI monotherapy stratified by sML, with log-rank test *P*-value shown **(c)** KM plot showing failure-free survival (FFS) among patients from the combination ARPi+IPI therapy, stratified by sML, with log-rank test *P*-value shown. **(d)** Forest plot depicting hazard ratios for Cox PH models for patients from (b) and (c). Displayed hazard ratios are from the final selected model, after performing bi-direction stepwise variable selection, with age, sex, sML and TMB as the base model, and sML x TMB interaction, PD-L1 density, purity, ploidy, and coverage as candidate predictors. (**e**) Boxplot quantifying distributions of CD8 T-cell density (cells/mm^2^) from immunohistochemistry staining in high versus low sML patient groups (n=8 vs. 13) from NCT02113657. *P*-value of the two-sided Wilcoxon rank sum test is shown. **(f)** Boxplot quantifying distributions of PD-L1 density from immunohistochemical assay in high versus low sML patient groups (n=8 vs. 13) from IPI monotherapy. *P*-value of the two-sided Wilcoxon rank sum test is shown. Significance levels are denoted as follows: \**P* < 0.05, \*\**P* < 0.01 and ***P < 0.001.

We used additional profiling data of these patients to determine whether sML corresponded to changes in the tumor immune microenvironment. We allocated paired immunohistochemistry slides from a subset of patients in the IPI monotherapy trial and observed high versus low sML tumor samples present significantly different levels of CD8^+^ T cell densities (Wilcoxon test *P*-value = 0.008, **Fig. 4e**. This distinction is independent of PD-L1 protein expression levels (Wilcoxon test *P*-value = 0.3, **Fig. 4f**). We further obtained CIBERSORTx^56^ deconvolved immune cell type proportions for the IPI trial. The deconvolution-based CD8^+^ T cell proportions showed a similar trend, although not statistically significant, while the myeloid cell population showed a significantly higher proportion of monocytes and a significantly lower proportion of macrophages in the high sML tumors (Wilcoxon test *P*-values = 0.004, 0.03, **Extended Data** Fig. 8c). Together the data suggest that a high subclonal mutation load corresponds with a more favorable tumor immune microenvironment, while a low subclonal mutation load may correspond with tumor-associated macrophage (TAM) mediated immunosuppression^57,58^, providing a plausible biological explanation for the prediction of responsiveness to ICB by high sML.

### Validating sML in low to moderate TMB cancers using a large patient cohort

Following our pan-cancer hypothesis, we move to another moderate TMB cancer type, esophageal adenocarcinoma (EAC), using WGS data generated by the OCCAMS Consortium, where detailed clinical follow-up was collected. Esophageal cancer is the eighth most diagnosed cancer and sixth most common cause of cancer-related death globally^59^, with a 10-30% 5-year survival rate^60^. Despite recent reports by OCCAMS^36,37,61^, knowledge of the evolutionary dynamics during disease progression remains incomplete, and better prognostic indicators are needed, especially those indicative of response to immunotherapy. This is of particular importance given that ICB achieves an overall response rate of lower than 30% in EAC and is predominantly restricted to the metastatic setting^62,63^.

We applied CliPP to perform subclonal reconstruction and then calculated sML in 613 WGS EAC samples from the OCCAMS consortium^35–37^ (**Methods**). The majority of these samples have TMB < 10 mutations/Mb (**Fig. 5a**). It took CliPP 2 hours to finish running the first 400 samples with < 35,000 SNVs, then about 10 hours for the remaining samples (**Supplementary Information Section 2.2**). Both TCGA and OCCAMS esophageal adenocarcinoma datasets shared similar fractions of high versus low sML samples, with the latter achieving a higher level of statistical significance in distinguishing patients’ OS outcomes using a larger sample size, i.e., patients with high sML had longer survival (TCGA log-rank test *P*-value = 0.08, **Fig. 5b**, OCCAMS log-rank test *P*-value = 0.0001, **Fig. 5c, Extended Data** Fig. 9, **Methods**). This separation in survival persisted across tumor stages (**Fig. 5d-e**) and between patients with or without a history of smoking (**Fig. 5f**). Hence, Cox regression analyses show high sML remains significantly associated with improved survival after adjusting for TMB, age, gender, and stage (**Fig. 5g**, **Supplementary Table 6**). CIBERSORTx deconvolved immune cell type proportions again reveal a significantly higher proportion of macrophages in the high sML group (Wilcoxon test *P*-value = 0.03, **Extended Data** Fig. 9g), which is consistent with our observation in the mCRPC IPI trial. Taken together, this provides evidence that the evolutionary dynamics captured by sML are a unique feature independent of age, gender, stage or TMB, and that it may provide insight into the TME state and clinical relevance that is orthogonal to currently used clinical biomarkers.

**Figure 5:**
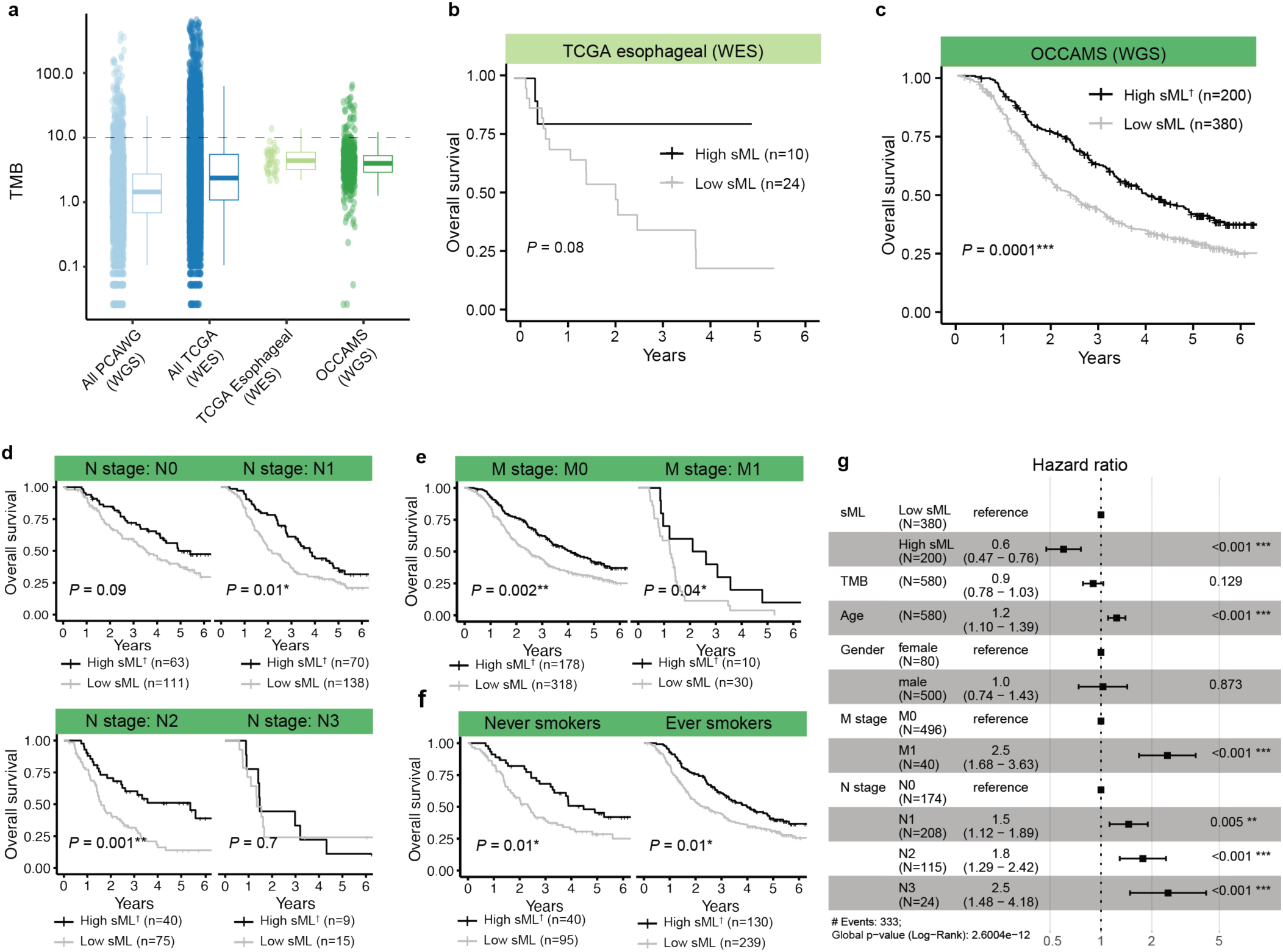
Validation of sML using WGS data from patients with esophageal adenocarcinoma. **(a)** Boxplots of TMB distributions for all PCAWG and TCGA samples, esophageal cancer samples from TCGA, and OCCAMS esophageal samples. Individual data points are shown next to the boxplots. **(b-c)** KM curves depict OS in esophageal adenocarcinoma patients from TCGA (**b**) and the OCCAMS cohort (**c**), divided by high and low sML. Within patients with high sML, we removed 33 samples with high SI (**Methods, Extended Data** Figs. 9a-b) to achieve a more homogeneous population of clonal expansion hence the remaining patient group is denoted as High sML†. **(d-f)** KM curves depicting OS difference between high and low sML esophageal adenocarcinoma patients from OCCAMS, with patients divided by N stage **(d)** M stage **(e)** and smoking history **(f)** Log-rank test *P*-values between high- and low-sML groups are shown**. (g)** HR and 95% CI for the significant terms in a Cox PH model for the OCCAMS esophageal adenocarcinoma samples, after performing stepwise variable selection. Smoking status and body mass index (BMI) were not retained in the variable selection process. \**P*<0.05, \*\**P*<0.01, \*\*\**P*<0.001.

## Discussion

In this study, we identify subclonal mutation load (sML) as a biologically and clinically relevant evolutionary marker in low-to-moderate TMB cancers. Using CliPP to analyze sequencing data from 9,972 tumors across ICGC, TCGA, two clinical trial cohorts and OCCAMS, we find sML to be prognostic of survival (PFI or OS) across 18 cancers. The prognostic effects are in opposite directions between a few high-TMB cancers (n=4) and the remaining majority of cancer types (n=14, **Fig. 6**), with the latter direction of high sML predicting favorable outcomes being novel compared to earlier studies^20^. High sML is also prognostic of ICB response in two metastatic prostate cancer (low-TMB cancer), where neoantigens alone cannot explain the favorable responses in low TMB tumors^34^. To our knowledge, this study provides the first observation associating cancer subclonal architecture with tumor-associated macrophage (TAM) activities, which compliments the recent study using single-cell RNA sequencing to characterize TAM as a key player in the immunotherapy resistance of prostate cancer^64^. These findings highlight that distinct evolutionary processes and tumor-immune evasion mechanisms persist in high versus low to moderate TMB cancers^21,34^. Future efforts to delineate these diverse mechanisms will have profound biological and clinical implications for cancer. Our CliPP-on-web Shiny app: https://bioinformatics.mdanderson.org/apps/CliPP is accessible to non-computational scientists, which will facilitate the discovery and dissemination of this study as well as future studies.

**Figure. 6:**
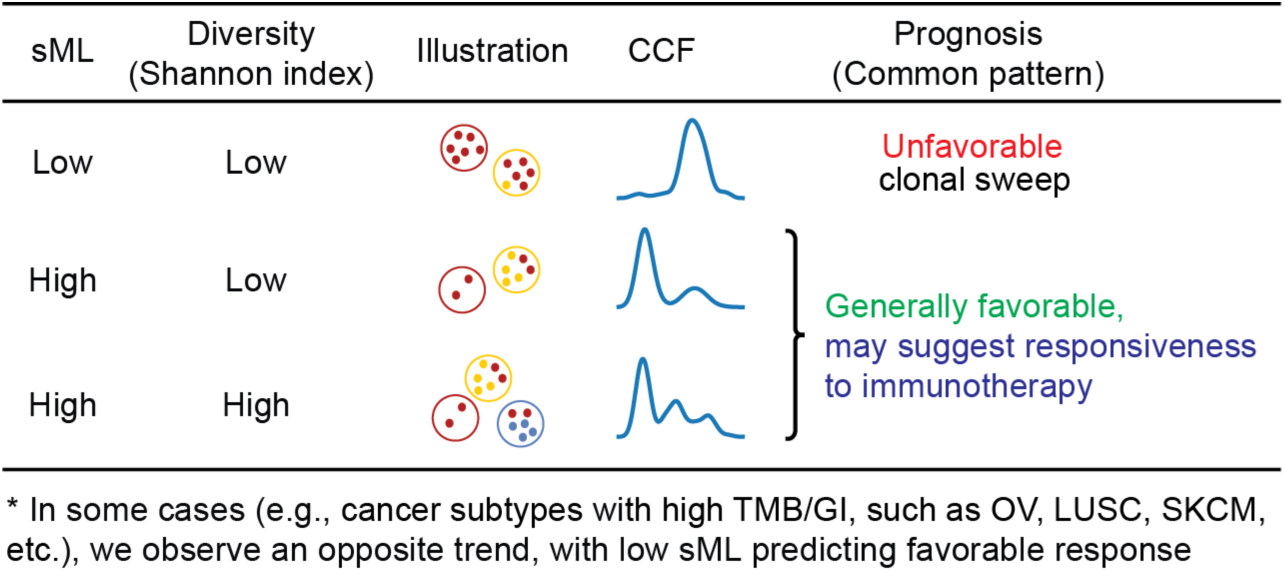
Summary and the concluding hypothesis. Illustration of the clinically relevant findings of sML. The red dots in the individual cancer cells represent clonal mutations, whereas the other dots represent subclonal mutations that are unique to their correspondingly colored cancer cell populations (yellow or blue).

Since the FDA approved TMB as a biomarker for ICB in 2020, clinical trials of ICB have disproportionately focused on high TMB cancers, with only recently pilot studies in low to moderate TMB cancers (clinicaltrials.gov), suggesting an urgent need to understand and better treat these cancers. Mismatch repair deficiency (MMRd), the only other clinically useful biomarker for ICB, shares similar biological underpinnings and is associated with high TMB cancers. Furthermore, MMRd occurs in only ∼3% of advanced prostate cancer patients^65^ and presents a 50% durable response rate^66^. Therefore, our finding of high sML predicting favorable ICB response in low TMB cancers is critical, as it cautions clinicians from recommending ICB for patients with low sML based on the current high TMB-biased literature^20^. Moreover, it highlights a new group of cancer patients who might benefit from ICB.

With limited resolution from single-sample sequencing, the load of subclonal mutations may fit under a variety of evolutionary models, such as branching evolution, punctuated evolution and neutral evolution^45^. Hence, we mainly interpret sML in terms of evolutionary phenomena such as local clonal sweep and the speed of mutation accumulation, which may happen multiple times during the process of any of these models. Thus without assuming any specific evolutionary model, we propose a plausible explanation: a low sML could indicate either local clonal sweeps or a lower mutation accumulation rate, while a high sML could indicate either prolonged latency before clonal sweep or a faster mutation accumulation rate. Therefore at a cancer-type population level, guided by the clinical outcomes, when high sML corresponds to a favorable prognosis, tumors likely have not reached a clonal sweep; when high sML corresponds to an adverse outcome, tumors may have gone through the clonal sweep. While most ICB strategies target late-stage tumors^67^, our observation that high sML is favorable in a majority of cancer types suggests that in low to moderate TMB cancers, tumors may have been sampled at an earlier evolutionary time point, hence earlier intervention may be more beneficial.

Immune evasion represents a key phenotypic trait driving tumor evolution^68,69^. Our observation suggests that low TMB tumors can exhibit immunogenicity, which varies with the subclonal architecture. While neoantigen diversity and quantity are widely accepted predictors of immunotherapy response, they do not fully explain ICB efficacy in low TMB responders. Factors such as oxidative stress and cell death resistance may also influence clonal selection and immunosurveillance^70–72^. The observed association of sML with T cell and myeloid populations suggests that for many histological types, tumor samples with a high sML may present a relatively friendlier immune environment, e.g., less obstruction from TAMs^58,64^. Further functional validation will be required to elucidate the relationship between tumor evolution, the immune system, and ICB efficacy, as well as address whether the sML feature can be generalized to predict the response of other cancer treatment modalities that require a functional immune environment.

Like other subclonal reconstruction callers, CliPP may struggle to determine precise subclonal mutation clusters from single-sample sequencing due to limited data complexity. Calculating sML, however, only requires the division of clonal versus subclonal mutations, which is less demanding than defining multiple subclones, e.g., the data requirement for calling one clonal cluster versus the rest is lower than calling four distinct clusters. In contrast, SI, relies more on cluster calling accuracy and may lack reliability. Overall, calculating sML from single-sample sequencing is informative for large-scale clinical studies, particularly for cancers lacking modernized datasets such as multi-region or single-cell sequencing^28,73^, as it focuses on all subclonal mutation cluster count rather than evolutionary roles. Future studies could investigate neutral tails to better interpret variations in sML for a specific histology type. However, it remains unclear whether existing methods^15^ can accurately detect neutral tails from WES data, which covers significantly fewer SNVs than WGS data.

In conclusion, a high-level summary measure of cancer evolution, such as the clonal/subclonal status of single-nucleotide variants, when computed across a large number of patient samples, sheds new light on the relationship between genetic intra-tumor heterogeneity and therapeutic resistance. Highly efficient and accurate methods like CliPP represent a new generation of techniques that bridge the gap between studying tumor biology and developing precision treatment strategies.

## Methods

Additional details and results are described in **Supplementary Information**. Here, we summarize the key aspects of the analysis.

### The CliPP Model

In the wake of the genomic revolution, several approaches have been developed to call and quantify cancer cell clones through clustering of variant read counts^74^. Many of the existing methods address subclonal reconstruction with mixture models, as implemented via Dirichlet processes, which are computationally costly, especially when the underlying subclonal structure is complex. CliPP is a novel statistical framework that addresses this task through penalized regression, drastically reducing the computational burden while maintaining high accuracy. It ensures homogeneity by introducing a pairwise penalty on parameter estimation without the need to pre-order coefficients, an advancement over preceding methods such as fused LASSO and CARDS algorithm. The full list of mathematical notations is provided in **Supplementary Information Section 8**. First, we assume that each sample is composed of two populations of cells with respect to the 𝑖-th SNV: one population containing normal cells or cancer cells without the SNV, and the other population of cancer cells harboring this SNV. We adopt the infinite sites assumption^75^ to assume that there is a single origin for each SNV. Within cancer cells, SNV 𝑖 may occur in one or a few cell populations (**Fig 1a**), therefore we introduce 𝛽_!_ to denote the cancer cell fraction (CCF) of SNV 𝑖, i.e., the fraction of cancer cells that carry SNV 𝑖. A clonal SNV exhibits a CCF of 1.0, indicating 100% presence in tumor cells. We further define the cellular prevalence (CP) of SNV 𝑖 as 𝜙_i_, so that 𝜙_i_ = 𝜌𝛽_i_, where 𝜌 denotes tumor purity, i.e., the proportion of cancer cells among all cells. Given 𝑆 SNVs, we assume that there exist 𝐾 groups of SNVs presenting distinct CCFs, where 𝐾 is significantly smaller than 𝑆. We expect SNVs from a common cancer cell population or subclone to have identical CP (or CCF) and correspondingly share the same parameter 𝜙_i_ (or 𝛽_i_). We follow an established model for observed variant read counts^12^, and assume that the observed number of variant reads 𝑟_i_at SNV 𝑖 follows a binomial distribution 𝐵𝑖𝑛𝑜𝑚𝑖𝑎𝑙(𝑛_i_, 𝜃_i_), where the total number of reads 𝑛_i_follow a Poisson distribution 𝑃𝑜𝑠𝑠𝑖𝑜𝑛(𝐷). We further express the expected proportion of the variant allele 𝜃_i_ as:

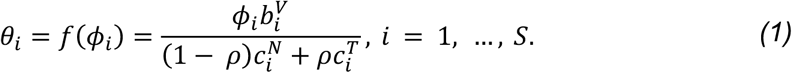

Hence the observed log-likelihood function, after canceling some constant, follows:

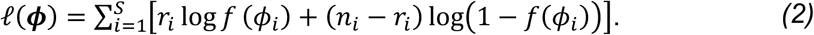

We use 𝑏_i_^V^ to denote the SNV-specific copy number at the 𝑖-th SNV. This measurement is distinct from allele-specific copy number when the SNV does not occur in all tumor cells. We then use 𝑐*_i_^N^* and 𝑐*_i_^T^* to !denote the total copy number for normal and tumor cells, respectively.

#### Input data

CliPP requires information of 𝑐*_i_^N^*, 𝑐*_i_^T^*, 𝑚*_i_^T^*, 𝑟*_i_*, 𝑛*_i_*, and 𝜌 as its input, which in order represent over SNV 𝑖: the total copy number for tumor cells, the total copy number for normal cells, the copy number of the major allele in tumor cells, the number of reads observed with variant alleles covering SNV and the total number of reads. Parameters 𝑐*_i_^T^*, 𝑚*_i_^T^*, and 𝜌 can be obtained using CNA-based deconvolution methods such as ASCAT^76^ and ABSOLUTE^77^, whereas 𝑐*_i_^N^* is typically set as 2. Parameter 𝑏*_i_^V^*,the SNV-specific copy number, is not directly observed or estimable using existing CNA software tools. Following a current convention that proved to be effective (see Eq.4)^74^, we assume 𝑏*_i_^V^* as:

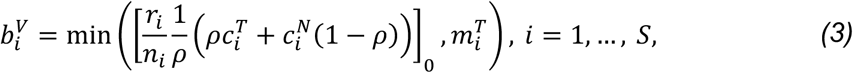

where [⋅]_0_rounds 𝑥 to the nearest positive integer. Here, using 𝑚*_i_^T^* as the upper bound enforces the logical constraint that even if an SNV happened before any copy number event, the number of alleles carrying said SNV cannot exceed the total number of major alleles.

#### Output data

CliPP provides the inferred subclonal structure, including the total number of clusters, the total number of SNVs in each cluster, the estimated CP for each cluster, and the mutation assignment, i.e., cluster ID for each mutation. This output can then serve as the basis for various downstream analysis including the inference of phylogenetic trees^74^.

### Parameter estimation using a regularized likelihood

The CPs, 𝝓 = (𝜙_1_, …, 𝜙_S_)^T^ may be estimated by maximizing the corresponding log-likelihood. However, our primary goal is to identify a homogeneous structure, i.e., clusters, of the CPs across all SNVs. Penalized estimation is a canonical tool to achieve homogeneity detection and parameter estimation simultaneously^78^. Therefore, we introduce a pairwise penalty to seek such homogeneity in 𝝓. To facilitate computation, we employ a normal approximation of the binomial random variable, i.e., 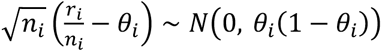, which lead to the following approximated log-likelihood

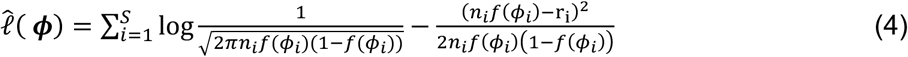

To promote homogeneity in 𝝓, we equip the negative log-likelihood 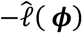 with a pairwise shrinkage penalty, thereby solving the following optimization problem

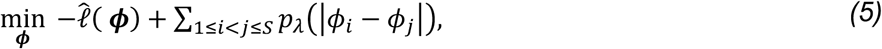

where 𝜆 > 0 is a tuning parameter that controls the degree of the penalization, and 𝑝_λ_(⋅) denotes a sparsity-inducing penalty function to identify the homogeneity structure in 𝝓, such as LASSO^79^, SCAD^41^, and MCP^80^ to name a few. Here, we focus on the SCAD penalty which is defined as

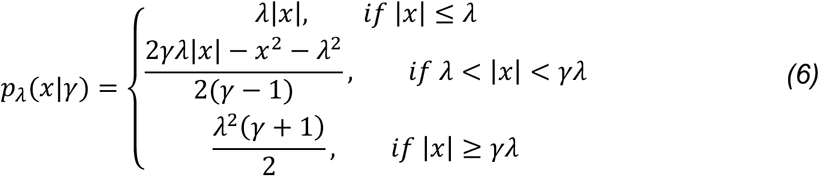

where 𝛾 is a hyper-parameter that controls the concavity of the SCAD penalty. Building upon the work of Fan and Li^41^, we set 𝛾 to 3.7. These concave penalties offer sparsity similar to the LASSO penalty, allowing them to automatically yield sparse estimates. It is proven that the SCAD-penalized estimators possess the oracle property in identifying sparsity structure of the data^41^. One can also choose other penalties, such as LASSO, and the formulation is straightforward.

Optimizing the objective function in Eq. (5) is nontrivial because the target variable 𝝓 is bounded, *i.e*., 𝜙_!_ = 𝜌𝛽_i_ ∈ [0,1], and the SCAD penalty is not convex. We employ a re-parametrization of 𝜙_!_ by defining 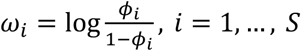, to remove the box constraint, which yields

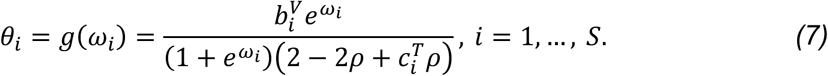

Note that 𝜔_!_ ’s are monotonic with respect to 𝜙_i_ ’s; 𝜔_i_ = 𝜔_8_ implies 𝜙_i_ = 𝜙_8_ and vice versa, thus the homogeneity pursuit of 𝝓 can be achieved by the homogeneity pursuit of 𝝎. We therefore perform a transformation on the loss function from Eq. (5), with an updated penalty function that identifies the homogeneity structure in 𝝎. Consequently, we reformulate the optimization problem as follows:

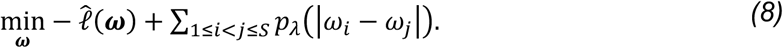

This is a non-convex optimization problem with respect to 𝝎, and its computation is non-trivial due to the complex log-likelihood. To address this challenge, we employ the alternating direction method of multipliers (ADMM)^81^, where we approximate the negative log-likelihood at each iteration by a quadratic function, thereby simplifying the computation. Note that the SCAD penalty possesses the unbiasedness property, ensuring that it does not shrink large estimated parameters throughout iterations. This property is particularly important in ADMM algorithms, as biases during the iterations could significantly impact the search for subgroups. Further mathematical derivations and the computational construction of ADMM can be found in the **Supplementary Information Sections 8.1-8.3**.

Finally, it is crucial for the regularized likelihood-based approach to select a proper 𝜆, in order to balance between over- and under-fitting. Here, the choice of 𝜆 determines the final number of clusters, with higher values yielding fewer clusters. Traditional approaches for selecting 𝜆, including cross-validation, bootstrapping, AIC, and BIC, suffer from respective disadvantages in our study. Cross-validation and bootstrapping require down-sampling and multiple rounds of estimation, causing a heavy computational burden, which conflicts with the major motivation of this work. AIC, BIC, and their extensions select 𝜆 by minimizing the negative log-likelihood equipped with a penalty term to control the model’s complexity. They are generally applicable to tuning parameter problems but lead to improper clustering results without incorporating the biological requirements in our study. To address this issue, we currently implement an *ad hoc* selection approach, which focuses on the interpretability of the outcome by ensuring that clonal mutations have an estimated CCF of around 1. Our automated 𝜆 selection pipeline operates as follows: for each sample, CliPP will run on the data with 11 different 𝜆′𝑠 spanning from 0 to 0.25, specifically: 0.01, 0.03, 0.05, 0.075, 0.1, 0.125, 0.15, 0.175, 0.2, 0.225, 0.25. For each sample, we compute a score 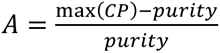. A lower score indicates a clonal CCF closer to 1, which is more consistent with our model assumption and unlikely to present a superclonal cluster. If there are one or more results that satisfy 𝐴 < 0.05, we choose the largest 𝜆 associated with those results. If all scores 𝐴 are greater than 0.01, we choose the 𝜆 associated with the smallest 𝐴. Note that when applying the CliPP software to any new datasets, the choice of 𝜆 can be decided by users.

CliPP has an automatic post-processing pipeline for better biological interpretability. It also allows for down-sampling when processing samples with huge amounts of SNVs. Please find the details in **Supplementary Information Section 8.4**.

### Benchmarking CliPP performance in simulated datasets

We assessed the ability of CliPP to correctly reconstruct the subclonal organization of a tumor on three simulated datasets. We generated an in-house simulation dataset CliPPSim4k to thoroughly benchmark the performance in samples with fewer copy number aberrations (CNAs) and higher read depth to cover the scenario of both whole-genome and whole-exome sequencing (WGS, WES) data (**Extended Data** Fig. 2a). We further included PhylogicSim500, a simulation dataset containing 500 samples where the copy number profiles were sampled from the PCAWG WGS data, with other parameters independently sampled from fixed distributions (**Extended Data** Fig. 2b). We also included the SimClone1000 dataset, a dataset containing 965 samples that were simulated based on a grid design to cover the spectrum of scenarios encountered in the PCAWG WGS data (**Extended Data** Fig. 2c). Details for both datasets can be found in Dentro et al^12^. CliPP was run on all datasets using the default setting, and PhyloWGS v1.0-rc2, PhylogicNDT, and PyClone-VI were run on CliPPSim4k using the recommended settings. We used available PhyloWGS results on the other two simulation datasets^12^. We evaluated the accuracy of each method using three metrics that measure bias in estimated number of clusters (rdNC), fraction of clonal mutations (rdCF), and CP across all variants (RMSE), respectively, and a total error score that averages over the three metrics to represent an overall performance. Please refer to **Supplementary Information Section 1** for further details.

### Subclonal reconstruction in PCAWG

The Pan-Cancer Analysis of Whole Genomes (PCAWG) study^12,82^ dataset comprises whole-genome sequencing (WGS) data obtained from a cohort of tumor samples spanning 39 cancer types. PCAWG related data were downloaded from the ICGC data portal, using Bionimbus Protected Data Cloud (PDC). The PCAWG dataset contains WGS data obtained from 2,778 tumor samples (2,658 distinct donors), including 2,605 primary tumors and 173 metastases or recurrences (minimum average coverage at 30x in the tumor)^82^. As input to CliPP, we use the consensus mutation calls derived from 4 distinct callers^82^ and the consensus CNA calls from 6 distinct callers^12^. In adherence to established best practices^45^, we employ the metric of reads per tumor chromosomal copy (nrpcc) to filter out samples with insufficient read coverage for subclonal reconstruction. We calculate 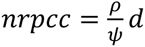, where 𝑑 is the depth of sequencing, 𝜌 is the purity and 𝜓 is the average tumor sample ploidy, i.e., 𝜓 = 𝜌𝜓*_T_* + (1 − 𝜌)𝜓*_N_*, where 𝜓_$_ is tumor cell ploidy and 𝜓_#_ = 2 is normal cell ploidy. We applied a stringent threshold of 𝑛𝑟𝑝𝑐𝑐 ≥ 10 to obtain a high-confidence cohort of 1,993 samples, following the PCAWG study^12^. For 149 samples with >50,000 SNVs we deployed the down-sampling strategy described in **Supplementary Information Section 8.4**.

To benchmark the accuracy of CliPP in real data, we used subclonal reconstruction results from 9 individual methods (Bayclone^83^, Ccube^84^, cloneHD^85^, CTPsingle^86^, DPClust^38^, Phylogic^32^, PhyloWGS^31^, PyClone^39^, Sclust^87^) and the consensus calls, which were available for 1,582 samples. We further ran CliPP and PyClone-VI^30^ on these samples. The comparison was performed based on the proportion of subclonal mutations estimated by each method and summarized by calculating the concordance correlation coefficient (CCC) for each pair of methods, including the consensus.

### Subclonal reconstruction in TCGA

The Cancer Genome Atlas (TCGA) contains whole-exome sequencing (WES) data from a cohort of tumor samples spanning 32 TCGA cancer types. The TCGA dataset consists of 9,654 WES samples with consensus mutation calls from 5 variant callers and 2 indel callers^88^ with matched copy number segments and tumor purity estimates from ASCAT^76^. The CliPP analysis pipeline for TCGA samples includes additional steps for quality control and filtering (**Extended Data** Fig. 1a). Sample quality control was performed using multiple criteria. Samples were flagged for a separate processing if they met any of the following conditions: (1) classified as ’likely normal’ by ASCAT (defined as Loss of Heterozygosity < 0.1, Genome Instability < 0.1, purity = 1, and sex-specific ploidy constraints [XX: 1.99-2.01; XY: 1.945-1.965]), (2) exhibited unrealistically high purity estimates (> 0.99), or (3) showed minimal genome instability (< 0.01). For these flagged samples, we implemented an alternative analysis pipeline: first assume no copy number alterations with purity set to 1, then use the CliPP estimated cellular prevalence of the clonal cluster to replace the purity estimate, and run CliPP a second time using this purity. In other words, CliPP can be used to estimate purity for quiet genomes where copy number changes are minimal. Additionally, samples were excluded if they had either too few SNVs (resulting in insufficient statistical power for clustering) or too many SNVs (indicating potential artifacts). Samples with low read coverage (nrpcc < 10) were excluded. The resulting cohort size in TCGA was 𝑛 = 7,279 from 32 cancer types.

### Computational speed benchmark

We measured the elapsed real time for each method from initiation to completion on the same machine equipped with Intel(R) Xeon(R) Gold 6,132 CPU @ 2.60GHz, using 28 CPU cores for consistency. In order to compute the speed for PhyloWGS, PyClone, and PhylogicNDT, which are time-consuming to study, we employed a grid-based sampling approach on the PCAWG dataset by selecting five samples across seven predefined SNV grids (100, 500, 1,000, 2,000, 3,000, 4,000 and 5,000 SNVs). In instances of sample scarcity at these exact SNV counts, the nearest equivalent (within a +/- 5% window) was selected. For the computational speed benchmark in TCGA WES data, we employed a strategy to match the distribution of relatively low SNV numbers to already obtained PCAWG data without further running these methods on more samples. Please refer to **Supplementary Information Section 2** for further details.

PhyloWGS v1.0-rc2 was executed using the same settings applied in the simulation study. PyClone v0.13.1 was run with both recommended and default configurations (https://github.com/Roth-Lab/pyclone). PyClone-VI v0.1.1 was applied to all PCAWG samples using recommended and default settings, incorporating 40 clusters for fitting (-c 40) and 100 restarts of variational inference (-r 100) (https://github.com/Roth-Lab/pyclone-vi). PhylogicNDT v1.0 (https://github.com/broadinstitute/PhylogicNDT) was executed for 1,000 iterations (-ni 1000), with the command to calculate cff histograms (--maf_input_type calc_ccf).

### Analysis of matched WGS and WES data from the same tumor samples

We identified a set of tumor samples (n=488) that contributed to the generation of both PCAWG WGS and TCGA WES data with neighboring slides through matching study IDs. To ensure a fair comparison of the two sequencing platforms, we removed samples with outlier TMBs, i.e. the 3^rd^ quartile+1.5x interquartile range (IQR) of all samples for each cancer type. The cohort comprised 488 samples with 124,725 SNVs before filtering, and after filtering, it consisted of 447 samples with a total of 49,554 SNVs. We then compared the CliPP output on these samples using Bangdiwala’s B statistic^89^, to quantify the agreement of clone/subclonal mutation assignment from WGS and WES data. This statistic can address zero counts in the contingency table better than the commonly used Kappa statistic, which makes it a better fit for this comparison. Interpretation thresholds for Bangdiwala’s B statistic were obtained from Munoz and Bangdiwala, and follow widely used and well established thresholds from other agreement metrics^90^. Bias in disagreement was assessed in the moderate to high agreement samples (B ≥ 0.49) and the low to moderate agreement samples (0.09 ≤ B < 0.49) using McNemar’s test. To evaluate the similarity between VAF distributions from WGS and WES samples, and therefore the appropriateness of comparing subclonal reconstruction results, we computed the Komogorov-Smirnov (K-S) test statistic and Wasserstein distance for each pair of sequencing data. We established background distributions through permutation analysis, performing 50,000 random samplings with replacement from the 447 overlapping WGS-WES sample pairs. Based on these empirical distributions, we defined thresholds for high dissimilarity (K-S statistic > 0.25 or Wasserstein distance > 0.04). We then compared agreement (Bangdiwala’s B statistic) between samples exceeding these thresholds and the remaining samples using a two-sided Wilcoxon rank-sum test. Finally, we compared sample barcodes to identify samples with different sources, by matching the sample, vial, portion, and analyte fields within the TCGA IDs. Agreement (measured by Bangdiwala’s B statistic) between sample pairs derived from the same versus different analytes were evaluated using a two-sided Wilcoxon rank-sum test. Please refer to **Supplementary Information Section 3** for further details.

### Subclonal mutational load (sML)

We introduce subclonal mutational load (sML) as the percentage of subclonal SNVs out of all SNVs. 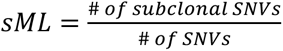. Here, we call the SNVs in the cluster with the highest CP as clonal SNVs and the rest as subclonal SNVs. We calculate tumor mutation burden (TMB) as the total number of SNVs per mega-base (Mb). With WES data, we restrict the TMB and sML calculations to the coding sequence (CDS). We annotate CDS as defined by GENCODE Release 19 (annotated as “gene_type” = “protein_coding” and “feature” = “CDS”). We use 38Mb as the exome size^91^.

### Driver mutation analysis

Driver mutations were identified and annotated by OncodriveMut^44^ for TCGA samples from 16 cancer types. After merging with the samples that passed CliPP filtering criteria, the resulting driver annotation set consisted of 2,469 samples with subclonal estimates for 8,586 SNV driver mutations in 574 unique genes. Two-sided Wilcoxon rank-sum test was used to compare mutation position distribution between clonal and subclonal drivers within genes, with no significant observed bias for any gene. To address sequence similarity issues affecting variant allele frequency (VAF) accuracy, mapping quality scores were computed for the 79 genes with at least 20 driver mutations. This process involved generating perfect 100-base read fastq files for every genome position and aligning them with BWA-mem. Subsequently, mapping quality scores ranging from 0 to 60 were extracted from the resulting BAMs. We then compare the distribution of mapping quality scores between clonal and subclonal mutations for the 9 genes containing annotated mutation sites with a mapping quality score less than 60 using two-sided Wilcoxon rank sum tests. We found evidence supporting significantly lower mapping quality scores in only 2 of these genes (*MLL3* and *NCOR1*, see **Supplementary Information Section 4**). These two genes were subsequently filtered out from our analyses and visualizations. Log-rank tests comparing overall survival and progression-free survival between clonal and subclonal mutations were performed for each gene-cancer pair with 9 or more subclonal driver mutations (16 pairs in total, **Supplementary Table 1**). For genes with frequently recurrent sites, additional survival analysis was performed, looking exclusively at these hotspot mutations. The survival analysis in *FAT1* mutated HNSC samples focused on HPV-negative samples, excluding a single HPV-positive case with a *FAT1* mutation.

### CliPP-on-Web

The CliPP web app was developed using CliPP (v1.3.3) as the back end, while the front end was built with R Shiny (v1.9.1) on R (v4.4.1). The app incorporates interactive visualizations powered by plotly (v4.10.4) and ggplot2. The CliPP web app enables users to run CliPP and visualize the distribution of VAF and estimated CP values across constructed subclones. The navigation bar allows users to upload and project their own samples using “CliPP-on-web”, query a sample from TCGA or PCAWG via “CliPP Data Resources”, and check the clonality of driver mutations in the TCGA cohort under the “Driver Mutation” feature. The control panel provides interactive options for selecting and uploading samples, and users can choose which sample to analyze and view. Additionally, the app provides an option to download the result files from CliPP.

### Association of sML with survival outcomes

To assess the clinical relevance of sML through its association with clinical outcomes, we characterized the relationship between sML and the patient survival data (overall survival (OS) and progression-free interval (PFI) on all 26 cancer types that present more than 50 samples, see a sample size summary in **Supplementary Table 2**). For well-defined clinical cancer subtypes and features, we further evaluated the effect of sML within these patient subgroups, including hormone receptor status in breast cancer, CMS subtype 2 (most frequently occurring) and MSI status in colorectal cancer, adenocarcinoma and squamous cell carcinoma in esophageal cancer, HPV status in head and neck cancer, *IDH1* mutation status in low-grade glioma, smoking status in lung cancer, Gleason score in prostate cancer, and endometroid status in uterine and endometrial cancer. The following cancer-type specific features were accessed using the TCGAquery_subtype function from the TCGAbiolinks Bioconductor package^92^: Gleason score for prostate cancer, PAM50 classification for breast cancer, smoking status for lung adenocarcinoma and lung squamous cell carcinoma, molecular subtypes for glioblastoma, Lauren classification and Epstein-Barr virus (EBV) status for stomach adenocarcinoma, microsatellite instability (MSI) and histological status for uterine corpus endometrial cancer. Additionally, we obtained human papillomavirus (HPV) status for head and neck squamous cell carcinoma from Lawrence et al^93^, and triple-negative hormone status for breast adenocarcinoma from Koboldt et al^94^. We also obtained the consensus molecular subtypes (CMS), a clinically utilized classification system for TCGA colorectal cancer (CRC)^95^.

Clinical data were obtained from the TCGA Pan-Cancer Clinical Data Resource (TCGA-CDR)^96^, which provides quality-controlled and standardized outcome data. Following TCGA-CDR recommendations, we included only survival endpoints rated as ’Yes’ or ’Caution’ for reliability within each cancer type. Treatment data, although available, were excluded from analysis due to incomplete documentation as noted in TCGA-CDR^96^. We first associated sML with clinical outcomes by splitting samples into high and low sML using a recursive partitioning survival tree model (rpart)^97^ for each cancer type, with the maximum tree depth constrained to 1 (sML cutoffs are provided in **Supplementary Information Section 5.2**). To ensure well-balanced grouping, we required at least 10 samples and ≥10% representation per sML group (“high” versus “low”) for each cancer type. To analyze survival outcomes, we applied Kaplan-Meier (KM) analysis, generating KM curves through the survminer R package^98^. These curves illustrate survival probability across time, stratified by high vs. low sML. Patient outcome stratification in cancer type or subgroups with log-rank test *P*-values ≤ 0.1 was deemed statistically significant or marginally significant.

We furthered our analysis on how sML impacts clinical outcomes along with other patient characteristics such as age and sex by utilizing multivariate Cox proportional hazard (PH) models. The PH assumption was deemed appropriate by testing the Schoenfeld residuals for survival data across cancer types. We employed multivariate CoxPH models in two iterations for all 26 cancer types: 1) using sML as a binary variable, high versus low along with continuous TMB, age, and sex where applicable, plus a model selection step using Bayesian information criterion (BIC) to include the interaction of binary sML and TMB, ploidy, purity, and sample read depth as candidate variables (the selected models are provided in **Supplementary Table 3**); and 2) using both sML and TMB as continuous variables, along with age, and sex where applicable, plus a model selection step using Bayesian information criterion (BIC) to include the interaction of continuous sML and TMB, ploidy, purity, and sample read depth as candidate variables (the selected models are provided in **Supplementary Table 4**). We report cancer (sub)types where the grouping is balanced for high/low sML (at least 10 samples and ≥10% of the total sample size per grouping), and the estimated hazard ratio for the main or interaction term of sML is statistically significantly or marginally significantly different from 1 (*P*-value ≤ 0.1). We excluded cancer types with wide confidence intervals, which is a common outcome of small sample size. When both OS and PFI show significant effects, or both cancer type and cancer (sub)type show significant effects of sML, we only present one outcome in the figures. All results of selected models are provided in **Supplementary Tables 3-4**.

Please refer to **Supplementary Information Section 5** for further details on evaluating confounders in the clinical relevance of sML, and **Supplementary Information Section 6**for further details on the evaluation of Shannon Index (SI) in TCGA.

### Analysis of WES data from NCT02113657 and NCT02703623

We obtained WES data from two clinical trials involving patients with metastatic castration-resistant prostate cancer (mCRPC). In the first trial, NCT02113657^34^, WES data were collected from 30 tumor samples prior to treatment with anti-CTLA-4 therapy (ipilimumab). In the second trial, NCT02703623^33^, WES data were collected from tumor samples of patients before treatment with abiraterone acetate, prednisone, and apalutamide (ARPi) for 8 weeks. Patients who showed a favorable response (≥50% PSA decline and <5 circulating tumor cells/7.5 mL) were then assigned to receive either ARPi alone (n=18), ARPi combined with ipilimumab (n = 21), or ARPi combined with carboplatin and cabazitaxel (CC) (n=27). Both datasets were processed using the same pipeline as follows. The paired-end FASTQ files were generated on the Illumina HiSeq 2500 for trial NCT02703623 and HiSeq 2000 for trial NCT02113657. After adapter trimming with Trim_Galore v0.6.10, sequence quality was assessed using FASTQC v0.11.8. Subsequently, the reads were aligned to the GRCh38 build through BWA-mem 0.7.15-r1140^99^. The resulting BAM files were processed according to the Genome Analysis Toolkit (GATK) Best Practices^100,101^, including steps such as marking duplicates, joint realignment of paired tumor-normal BAMs, and recalibration of base quality scores. Consensus mutation calls were derived as those from 2 out of 3 callers: MuSE 2.0^102^ (v2.1.2) calls and PASS-filtered results from MuTect2^103^ (v4.2.4.0) and Strelka2^104^ (v2.9.10). Copy number aberrations (CNAs), tumor purity and ploidy were called using ASCAT^76^ (v3.1.2) configured for WES data processing. Patients’ response to immunotherapy outcome data as well as immunohistochemistry (IHC) staining for CD8 T cell densities were obtained from the previous report of NCT02113657, as was PD-L1 expression, which was assessed using an automated immunohistochemical assay^34^. Following the pre-processing pipeline used for TCGA, we removed samples with nrpcc<10 and TMB>100, and further removed samples with less than 10 coding SNVs (i.e., TMB > 0.25) which would otherwise inflate the subclonal fraction estimates (**Fig. 4b**).

For NCT02113657 samples, since some patients have multiple metastases, we retained only the sample with the lowest sML for each patient to better align with survival outcomes. We then set 0.5 as the sML cutoff and classify patients with sML ≥ 0.5 as high sML. To analyze survival outcomes, we applied Kaplan-Meier (KM) analysis, generating KM curves through the survminer R package^98^. These curves illustrate survival probability across time, stratified by high vs. low sML. Survival *P*-values were obtained via log-rank test. *P*-value for comparison of CD8 T-cell density and for PD-L1 density between high and low sML samples was calculated using a two-sided Wilcoxon rank-sum test. We employed a multivariate Cox proportional hazards model, implementing bi-directional stepwise variable selection with model comparison via Bayesian information criterion (BIC) to identify the optimal set of variables. The base model used sML as a binary variable (high versus low) along with continuous TMB, and age. Additional candidate variables included PD-L1 density, purity, ploidy, coverage, and the interaction between sML and TMB (**Supplementary Table 6**).

For NCT02703623 samples, within each treatment arm, we split samples into high and low sML using a recursive partitioning survival tree model (rpart)^97^, with the maximum tree depth constrained to 1. Survival *P*-values were computed via log-rank test. We employed a multivariate Cox proportional hazards model, implementing bi-directional stepwise variable selection with model comparison via Bayesian information criterion (BIC) to identify the optimal set of variables. The base model used sML as a binary variable (high versus low) along with continuous TMB, and age. Additional candidate variables included PD-L1 density, purity, ploidy, coverage, and the interaction between sML and TMB (**Supplementary Table 6**). Age information was only available for 17 of the 21 patients.

### RNAseq data processing for NCT02113657

Libraries were sequenced on the Illumina HiSeq system with 76-bp paired-end reads. Raw sequencing data quality was assessed using FastQC (version 0.11.9) (https://www.bioinformatics.babraham.ac.uk/projects/fastqc/), and low-quality reads and adapter sequences were trimmed using Trimmomatic^105^ (version 0.4.0). Following quality control, the reads were aligned to the GRCh38 genome using the two-pass mode of STAR^106^ (version 2.5.3a), and read counts were generated using HTSeq^107^.

### Analysis of OCCAMS WGS data

The OCCAMS WGS cohort includes 710 patients, with 496 samples sequenced by Illumina or the CRUK Cambridge Institute, and 214 sequenced by the Sanger Institute as part of the CRUK Grand Challenge Mutographs of Cancer project. We obtained SNV and CNA calls from previous studies^36,61^. Briefly, quality control was performed with FastQC (http://www.bioinformatics.babraham.ac.uk/projects/fastqc) and in-house tools. For mutation calling, sequencing reads were aligned to the reference genome GRCh37 using the latest BWA-mem version. The aligned reads were sorted by genomic coordinates, and duplicates were removed using Picard (http://broadinstitute.github.io/picard). Strelka^108^ (v2.0.15) was used for calling SNVs and small indels. Additionally, MuTect2 was run on 10 randomly selected samples to validate Strelka calls. Somatic variants were called using MuTect2^103^ (v4.1.7.0) in matched normal mode, with a panel of normals and a population germline resource, using default settings throughout^37^. For copy number variation analysis, ASCAT^76^ (v2.1) was used to infer copy number alterations and estimate tumor purity and ploidy. Clinical/pathological information was obtained, including patient gender, age at diagnosis, survival time, histology classification, smoking history, alcohol intake history, height, weight, BMI, TNM staging, and treatment response.

We ran CliPP and calculated sML on 710 samples, removing 93 that had nrpcc <10 or with extremely high TMB (TMB>100) and 4 samples without survival information. Using this WGS dataset, we confirm that there is a strong correlation between sML (of the coding region) and subclonal fraction (on all SNVs) in OCCAMS dataset, suggesting that either estimate can be used in downstream analyses. For consistency with the TCGA results, we use CDS sML for all subsequent analysis of the OCCAMS ESAD samples. We split samples into high and low sML using a recursive partitioning survival tree model (rpart)^97^, with the maximum tree depth constrained to 1. Among high sML samples, we further apply rpart to partion the samples based on their Shannon Index (SI).

We furthered our analysis on how sML impacts clinical outcomes along with other patient characteristics such as age and sex by utilizing multivariate Cox proportional hazard (PH) models. sML status along with TMB, age, gender, M stage, N stage, smoking status, and patient BMI were included as covariates. We used stepwise model selection with Bayesian information criterion (BIC) to identify the most informative set of predictors for patient survival (**Supplementary Table 6**).

### RNAseq data processing for OCCAMS

Raw sequencing reads from 306 tumor samples and 16 normal samples were quality-controlled and trimmed before alignment to the human reference genome (GRCh37) using STAR^106^ aligner. Following alignment, reads mapped to gene features were quantified using the union and intersection-strict counting methods to generate raw read counts. Gene expression values were then normalized to account for both gene length and sequencing depth, calculating both Reads Per Kilobase of transcript per Million mapped reads (RPKM) and Transcripts Per Million (TPM) values.

### Cellular deconvolution

Cellular composition was estimated for samples from the IPI monotherapy trial and OCCAMS using the Impute Cell Fractions module of CIBERSORTx (https://cibersortx.stanford.edu/). Bulk RNA-seq data were deconvolved with the LM22 immune cell-type reference matrix to infer relative immune cell proportions. To correct for technical variation, we applied the recommended built-in batch correction method (batch mode = “B”). Deconvolution was performed in ’absolute’ mode with 1,000 permutations.

## Data availability

Raw read counts of bulk DNA sequencing data, clinical data and somatic mutations from 10,039 tumor samples across 32 TCGA cancer types are available for download from the Genomic Data Commons data portal (https://portal.gdc.cancer.gov/). The tumor purity and copy number profiling of TCGA samples were obtained using ASCAT v3^76^ and the results were released on https://github.com/VanLoo-lab/ascat/tree/master/ReleasedData.

PhylogicSim500 and SimClone1000 dataset can be downloaded from https://data.mendeley.com/datasets/compare/by4gbgr9gd.

The PCAWG dataset is available through the ICGC data portal: https://dcc.icgc.org/pcawg. Specifically, the SNV sequencing reads can be downloaded through https://dcc.icgc.org/repositories, while the CNV and purity data can be accessed through https://dcc.icgc.org/releases/PCAWG/consensus_cnv. Sample information, including mapping from ICGC identifiers to TCGA identifiers was downloaded through the ICGC portal.

The previous subclonal reconstruction results from 11 methods and the consensus calls can be downloaded from Zenodo at https://doi.org/10.5281/zenodo.14510686.

Driver mutation annotations were downloaded from IntOGen on https://www.intogen.org/download?file=intogen_driver_mutations_catalog-2016.5.zip

OCCAMS sequencing BAM files (WGS of primary tumors and matched normal) used in this study are publicly available at the European Genome-Phenome Archive (EGA) under accession codes EGAD00001007785 (Illumina/CRUK) and EGAD00001006083 (Sanger Institute).

Subclonal structure results of all samples and the identified subclonality of driver mutations for this study are available for download at https://odin.mdacc.tmc.edu/∼wwang7/CliPPdata.html (for reviewers only, password: wwylab_CliPP). All data can be visualized on the CliPP-on-web at https://bioinformatics.mdanderson.org/apps/CliPP. Alternatively, CliPP data for CliPPSim4k, TCGA, and PCAWG can be accessed on Zenodo at [https://doi.org/10.5281/zenodo.14008458] (currently restricted, access available upon request).

All other relevant data are available from the corresponding author upon reasonable request.

## Code availability

The CliPP model used for reconstructing tumor subclonal structure is freely available and can be downloaded from https://github.com/wwylab/CliPP. CliPP version 1.3.3 was used to generate the results in this work. Additionally, the code for the Shiny app can be found at https://github.com/wwylab/CliPP-ShinyApp.

## Author contributions

Y.J., M.D.M, K.Y., S.J.S., W.W. conceived the project. Y.J., K.Y., S.J.S., H.Z. and W.W. developed the mathematical method. Y.J., K.Y. and S.J. built the software tool, Y.J. and M.D.M performed the data analysis and interpretation, planned the figure design, and wrote the manuscript collaborating with all the other authors. S.J., S.G., Q.T., Y.D. and S.C. assisted on data analysis and interpretation. R.L., J.C.L., Q.T, and X.L. assisted with data maintenance and manuscript writing. Y.T. contributed to the code optimization. T.L. assisted with the copy number data. S.K., J.P.S., J.A., A.S., J.R.W. and P.M. advised on cancer-type specific clinical outcome analysis. S.S., P.S. and A.A. contributed the clinical trial dataset for validation. S.S. and A.A. advised on data analysis for prostate cancer. M.T. and P.V.L assisted in data interpretation, manuscript writing, and copy number profiling. M.K., X.L. and H.Z. advised on mathematical modeling. R.C.F, G.D. and C.M.J provided the OCCAMS data and assisted with data analysis and interpretation related to these data. W.W. conceived the study, planned and supervised the work, performed the analysis, and wrote the manuscript collaborating with all the other authors. All authors contributed to the interpretation of results and commented on and approved the final manuscript.

## Acknowledgements

Y.J. is supported in part by RP210006. Y.J., M.D.M., S.J. and W.W. are supported by R01CA268380. S.G., Q.T. and W.W. are supported by DoD PC210079. Q.T. is supported in part by R01CA275990. J.C.L is a TRIUMPH Fellow in the CPRIT Training Program (RP210028). R.L. is supported in part by R01CA283402. J.A. is supported by the Caporella Family fund, the Kohn family fund, the Shipman Family fund, and Stupid Strong Foundation. P.M. was supported by the National Cancer Institute R37CA288448 and R01CA285454, the Andrew Sabin Family Foundation Fellowship, Gateway for Cancer Research, a Translational Research Partnership Award (KC200096P1) and an Idea Development Award (RA230062) by the United States Department of Defense, an Advanced Discovery Award by the Kidney Cancer Association, a Translational Research Award by the V Foundation, the MD Anderson Physician-Scientist Award, donations from the Renal Medullary Carcinoma Research Foundation in honor of Ryse Williams, the Finneran Family Endowment, as well as philanthropic donations by the Chris “CJ” Johnson Foundation, and by the family of Mike and Mary Allen. T.L. was supported by the Francis Crick Institute which receives its core funding from Cancer Research UK (CC2008), the UK Medical Research Council (CC2008), and the Wellcome Trust (CC2008). A.K.S. was funded by NCI (CA281701 and CA209904), Ovarian Cancer Research Fund Alliance, American Cancer Society Research Professor Award, and the Frank McGraw Memorial Chair in Cancer Research. S. J. Shin was supported by the National Research Foundation of Korea (NRF) grant funded by the Korea government (MSIT), grant number 2022M3J6A1063595, and 2023R1A2C1006587. Research reported in this publication was supported by the National Institute On Aging of the National Institutes of Health under Award Number RF1AG082938. The content is solely the responsibility of the authors and does not necessarily represent the official views of the National Institutes of Health. J.P.S is supported by the Cancer Prevention & Research Institute of Texas as a CPRIT Scholar in Cancer Research (RR180035) and Conquer Cancer (CDA-7604125121). J.R.W. was supported by the Mark Foundation for Cancer Research ASPIRE award. M.K. is supported in part by R01CA268380. P.V.L. is a Winton Group Leader in recognition of the Winton Charitable Foundation’s support towards the establishment of The Francis Crick Institute. P.V.L. is a CPRIT Scholar in Cancer Research and acknowledges CPRIT grant support (RR210006). R.C.F. is a recipient of a Core Programme Grant from the Medical Research Council [MR/W014122/1]. C.M.J. is supported by a Clinical Lectureship part-funded by Cancer Research UK RadNet Cambridge [C17918/A28870]. The Oesophageal Cancer Clinical & Molecular Stratification (OCCAMS) study and its successor OCCAMS 2 study were supported by Cancer Research UK [A15874, A22720, A22131]. This work uses data provided by patients and collected by the NHS as part of their care and support. We thank the Human Research Tissue Bank, which is supported by the UK National Institute for Health Research (NIHR) Cambridge Biomedical Research Centre, from Addenbrooke’s Hospital. The views expressed are those of the authors and not necessarily those of the NIHR or the Department of Health and Social Care. The results presented here are in part based upon data generated by the TCGA Research Network: https://www.cancer.gov/tcga. We thank John Weinstein for his valuable feedback on this manuscript. Special thanks to Sylvain Laroche and Chris Wakefield from the Quantitative Research Computing team for their assistance in deploying the CliPP-on-web and enabling public access. The author(s) acknowledge the support of the High Performance Computing for research facility at the University of Texas MD Anderson Cancer Center for providing computational resources that have contributed to the research results reported in this paper.

## Declaration of interests

This work is associated with a patent filing by W.W., Y.J. and M.D.M. at MD Anderson Cancer Center. Y.J. is currently a Senior Statistician at AbbVie. S.S. consults for Bristol Myers Squibb. P.M. has received honoraria for service on a Scientific Advisory Board for Mirati Therapeutics, Bristol Myers Squibb, and Exelixis; consulting for Axiom Healthcare Strategies; non-branded educational programs supported by DAVA Oncology, Exelixis and Pfizer; and research funding for clinical trials from Regeneron Pharmaceuticals, Takeda, Bristol Myers Squibb, Mirati Therapeutics, Gateway for Cancer Research, and the University of Texas MD Anderson Cancer Center. A.K.S. Consults for GSK, Astra Zeneca, Merck, ImmunoGen, Onxeo, Iylon, and Advenchen. P.S. is a scientific advisory committee member for Achelois, Afinni-T, Apricity, Asher Bio, BioAlta LLC, Candel Therapeutics, Catalio, Carisma, C-Reveal Therapeutics, Dragonfly Therapeutics, Earli Inc, Enable medicine, Glympse, Henlius/Hengenix, Hummingbird, ImaginAB, InterVenn Biosciences, Lava Therapeutics, Lytix Biopharma, Marker Therapeutics, Oncolytics, PBM Capital, Phenomic AI, Polaris Pharma, Trained Therapeutix Discovery, Two Bear Capital, Xilis, Inc. Additionally, P.S. has private investments in Adaptive Biotechnologies, BioNTech, JSL Health, Sporos, and Time Bioventures. R.C.F is a shareholder in Cyted Health and on the advisory board for Astra Zeneca and CRUK Functional Genomics Centre and consults for Astra Zeneca, 23andMe. The other authors declare no competing interests.

All supplementary tables are provided as separate excel files, and are additionally available at https://drive.google.com/drive/folders/1Lz-wP_wt_h8tHdetjGLZTiHwNEhS31zq

**Supplementary Table 1.** Log-rank tests for cancer-type driver gene pairs with 9 or more subclonal mutations.

**Supplementary Table 2.** Sample size and number of survival events among cancer types from TCGA, OCCAMS and clinical trial cohorts.

**Supplementary Table 3.** Multivariate Cox proportional hazard models for binary sML for OS and PFI across cancer types in TCGA. Variable selection using BIC was performed. The base model includes age, sex, binary sML (high vs low), and TMB. Candidate predictors include sML x TMB interaction, purity, ploidy, and coverage. The final selected model is presented for each cancer type.

**Supplementary Table 4.** Multivariate Cox proportional hazard models for continuous sML for OS and PFI across cancer types in TCGA. Variable selection using BIC was performed. The base model includes age, sex, binary sML (high vs low), and TMB. Candidate predictors include sML x TMB interaction, purity, ploidy, and coverage. The final selected model is presented for each cancer type.

**Supplementary Table 5.** Clinical characteristics of patients from the two clinical trials in mCRPC stratified by high and low sML.

**Supplementary Table 6.** Selected multivariate Cox proportional hazard models for mCRPC clinical trial cohorts and OCCAMS esophageal adenocarcinoma cohort. Models for immunotherapy clinical trials included age, binary sML (high vs low), PD-L1 density, TMB, sMLxTMB interaction, purity, ploidy, and coverage as candidate predictors, and age, sex, binary sML (high vs low), TMB, sMLxTMB interaction, N stage, M stage, smoking history, and BMI were used as candidate predictors for OS in OCCAMS esophageal adenocarcinoma patients. Models selected by bi-directional stepwise variable selection.

---

**Figure S1:**
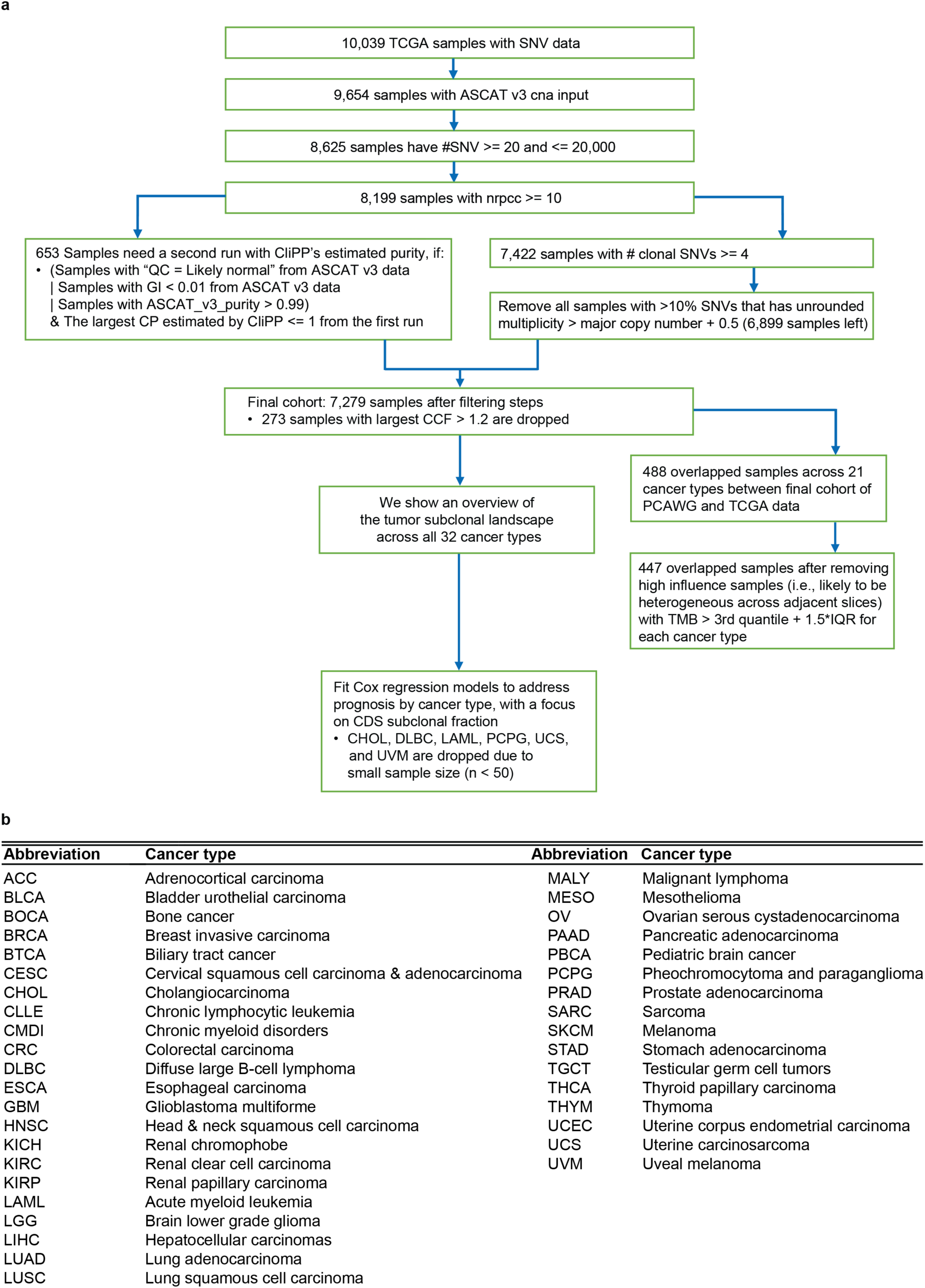
Overview of the TCGA data analysis. **(a)** CONSORT diagram for data processing in TCGA. **(b)** Acronyms used for cancer types analyzed in this study.

**Figure S2:**
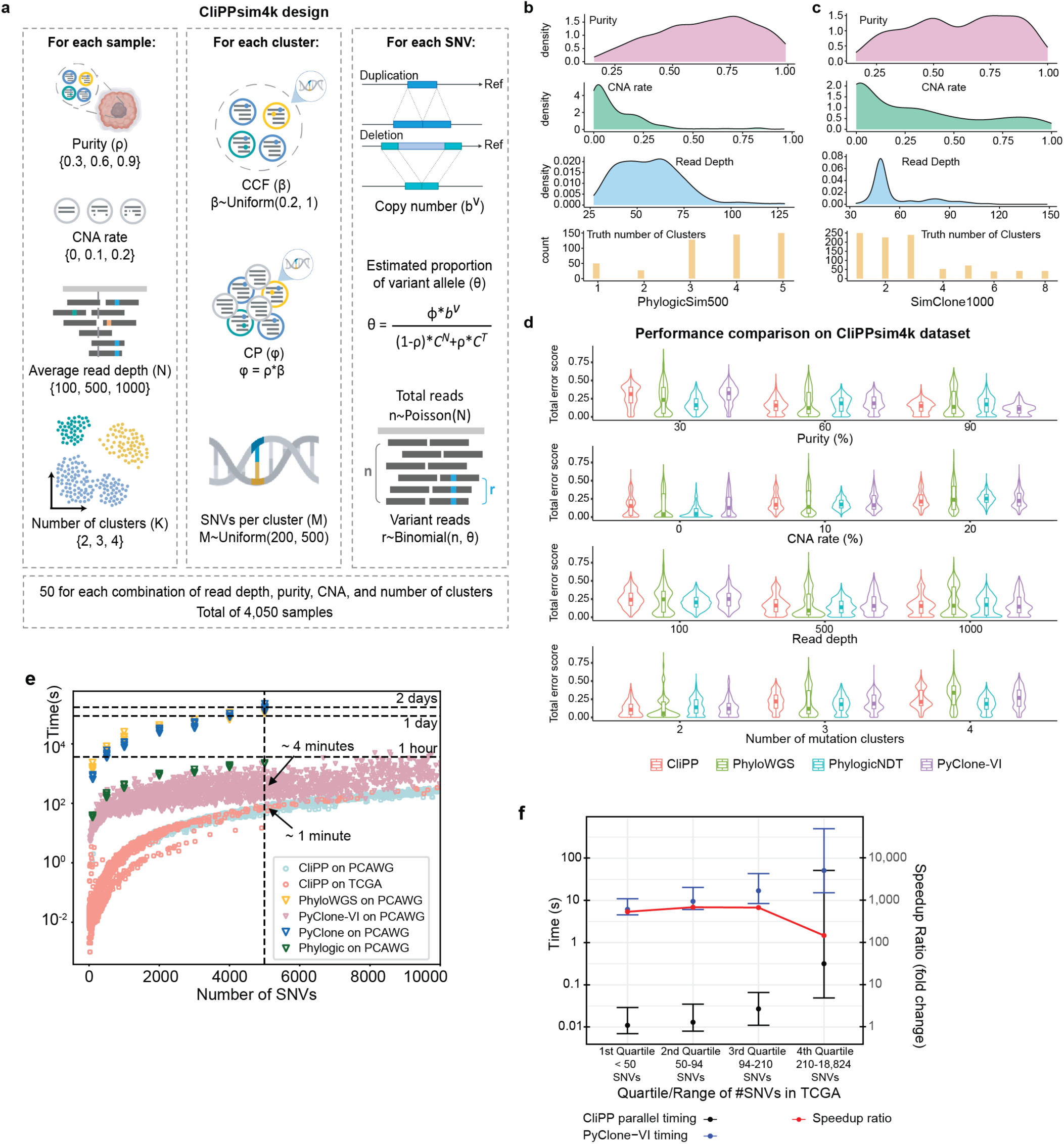
Performance benchmarking using both simulated and patient cohort datasets. **(a)** Overview of the simulation design for our CliPPSim4K dataset used to benchmark subclonal reconstruction performance. 50 samples are generated for each of 81 total combinations of sample-level parameters (purity, average read depth, CNA rate, and number of clusters), resulting in a simulated cohort of 4,050 samples. Within each sample, further parameters are simulated per cluster (CCF, CP, number of SNVs), and per SNV (copy number, proportion of the variant allele and total reads) to generate **Extended Data** Figure 2 **(cont’d):** variant and reference reads (see **Methods**). **(b-c)** Distributions of purity, CNA rate, read depth, and number of clusters across samples from PhylogicSim500 and SimClone1000. **(d)** Total error score (**Supplementary Information Section 1.1**) comparisons between CliPP, PhyloWGS, PhylogicNDT, and PyClone-VI on the CliPPSim4k dataset. **(e)** Computational times for subclonal reconstruction on 2,146 PCAWG WGS samples and 9,843 TCGA WES samples. The vertical dashed line corresponds to samples with 5,000 SNVs. Labels indicate average computing times for 5 samples with ∼5,000 SNVS: CliPP (1 minute) and PyClone-VI (4 minutes). Other methods’ timings: PhylogicNDT (37 minutes), PyClone (2,765 minutes), and PhyloWGS (2,450 minutes). **(f)** Time comparison between CliPP and Pyclone-VI in TCGA samples was categorized into four quartiles based on the number of SNVs. Dots correspond to the median time per quartile of samples for each method (see **Methods**). Error bars depict the 95% confidence intervals. The red line indicates the speedup ratio of CliPP relative to PyClone-VI.

**Figure S3:**
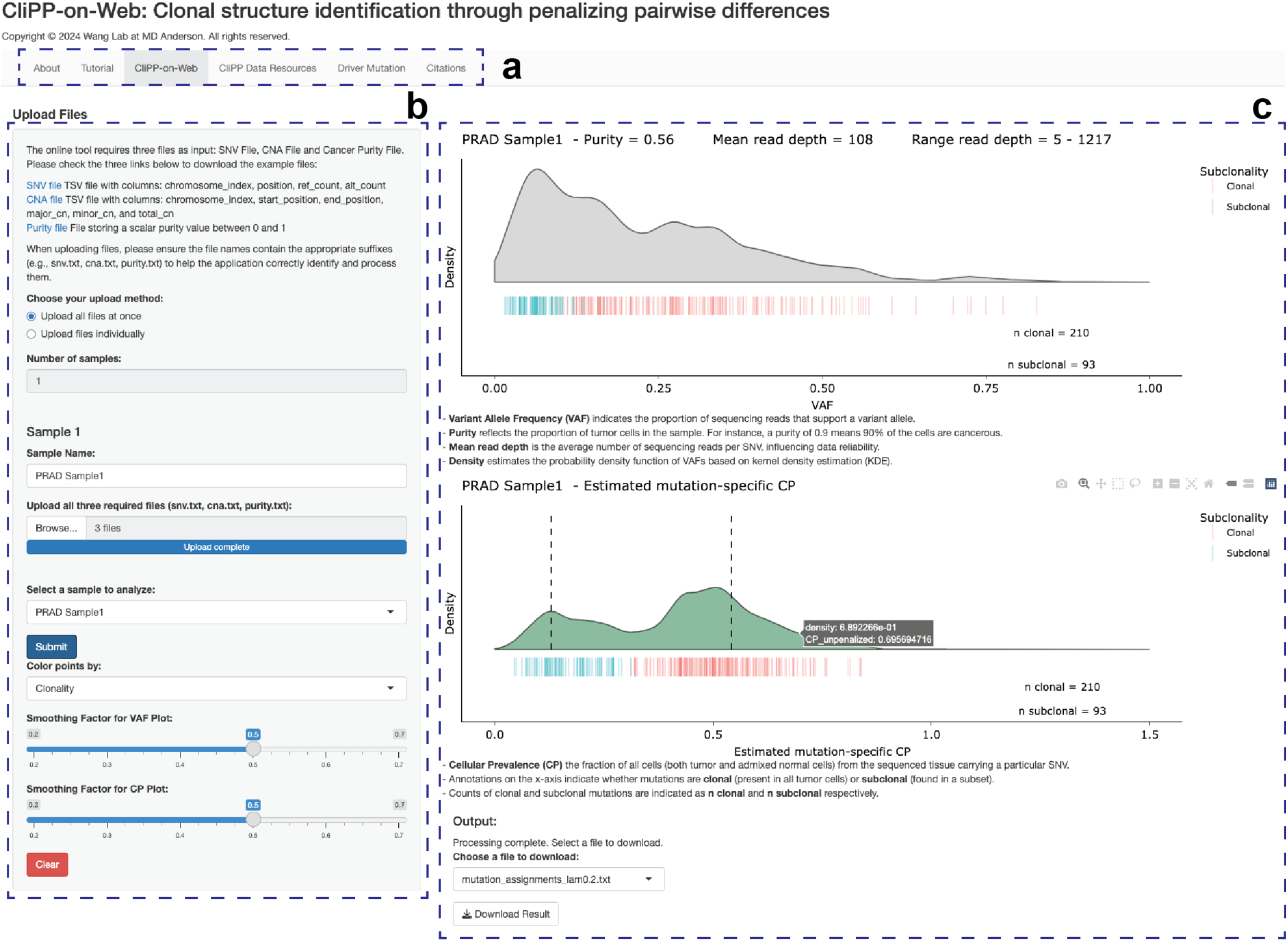
Graphical output of the CliPP-on-web Shiny app. **(a)** The navigation bar lets users upload and analyze their samples through “CliPP-on-Web”, query samples from TCGA or PCAWG via “CliPP Data Resources”, and view clonal and subclonal driver information for TCGA samples via “Driver Mutation Clonality”. **(b)** The control panel on the left allows users to interact with the web app, upload samples, and select which sample to analyze or view. **(c)** Visualization of the VAF and CP distributions per sample. After performing subclonal reconstruction on a given sample(s), graphs showing the VAF and CP distribution are generated, with lines in the “lawn” under the density curve representing individual SNVs, colored by clonality (red = clonal, blue = subclonal). On the CP density plot, dashed vertical lines indicate the estimated cluster CP by CliPP. Relevant information, including read depth, tumor purity, and number of mutations is printed on the graphs. An option to download the CliPP output in txt files is provided. The CliPP-on-web Shiny app is available at https://bioinformatics.mdanderson.org/apps/CliPP.

**Figure S4:**
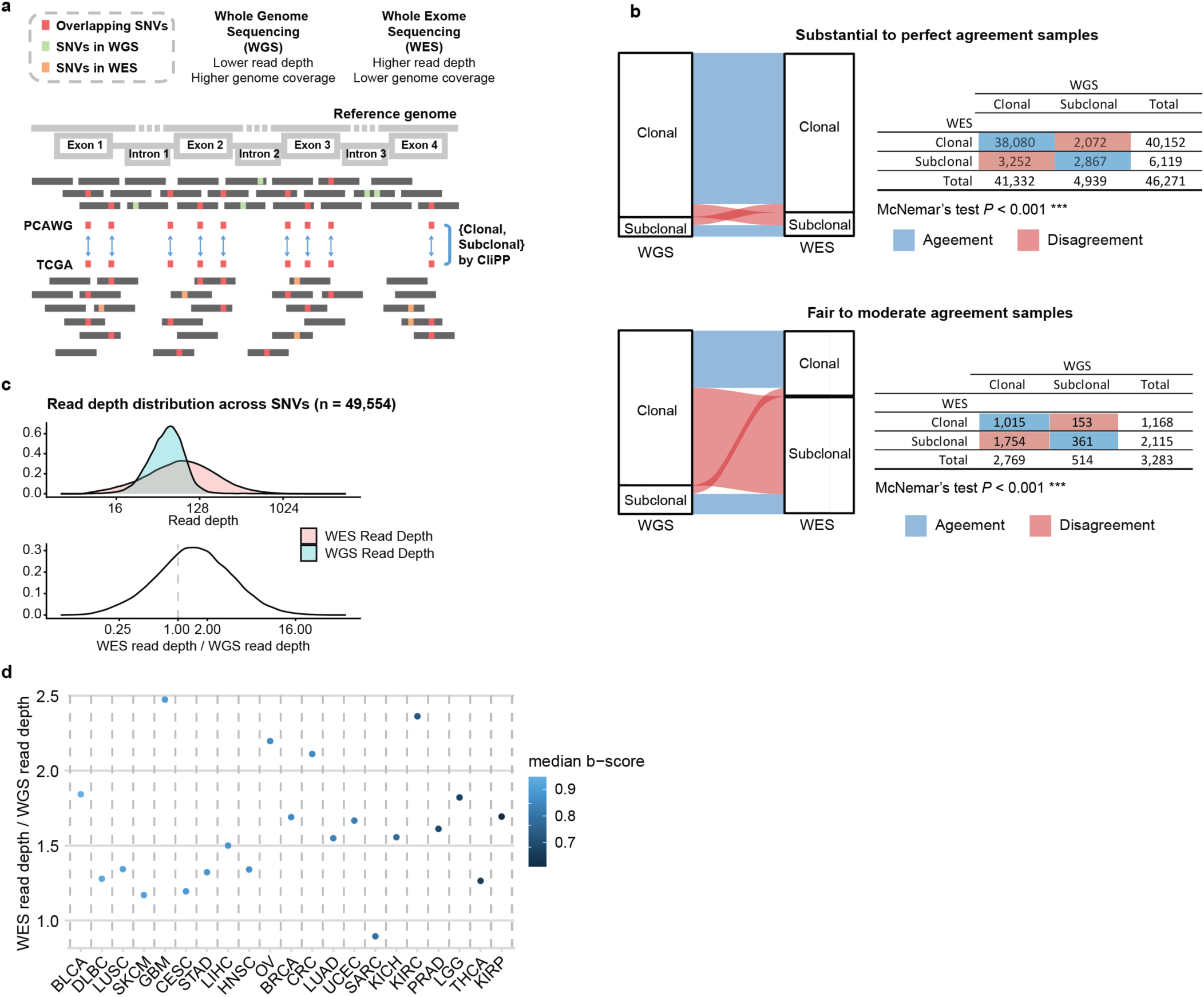
Benchmarking subclonal reconstruction pan-cancer using WES data. **(a)** Illustration of coverage and read depth differences between WGS and WES data. SNVs observed in samples sequenced by both methods are shown in red for cross-platform subclonal reconstruction comparison. **(b)** Alluvial plots comparing the clonality assignment per mutation from WGS to WES. Mutations in samples with moderate-to-high agreement samples (B ≥ 0.49) are shown on the left. Mutations in samples with low-to-moderate agreement (0.09 ≤ B < 0.49) are shown on the right. Each alluvial plot is paired with a two-way contingency table with detailed numbers, with *P*-values indicated for McNemar’s test for a biased shift between the red discordant squares. **(c)** Read depth distribution across SNVs for WES and WGS data (top) and the distribution of ratios of read depths between WES and WGS across SNVs (bottom). Vertical dotted line at 1 indicates equal read depth between WES and WGS. **(d)** Median read depth ratio of each overlapped sample from TCGA WES dataset and PCAWG WGS dataset by cancer type, ordered by the median B statistic (see **Methods**).

**Figure S5:**
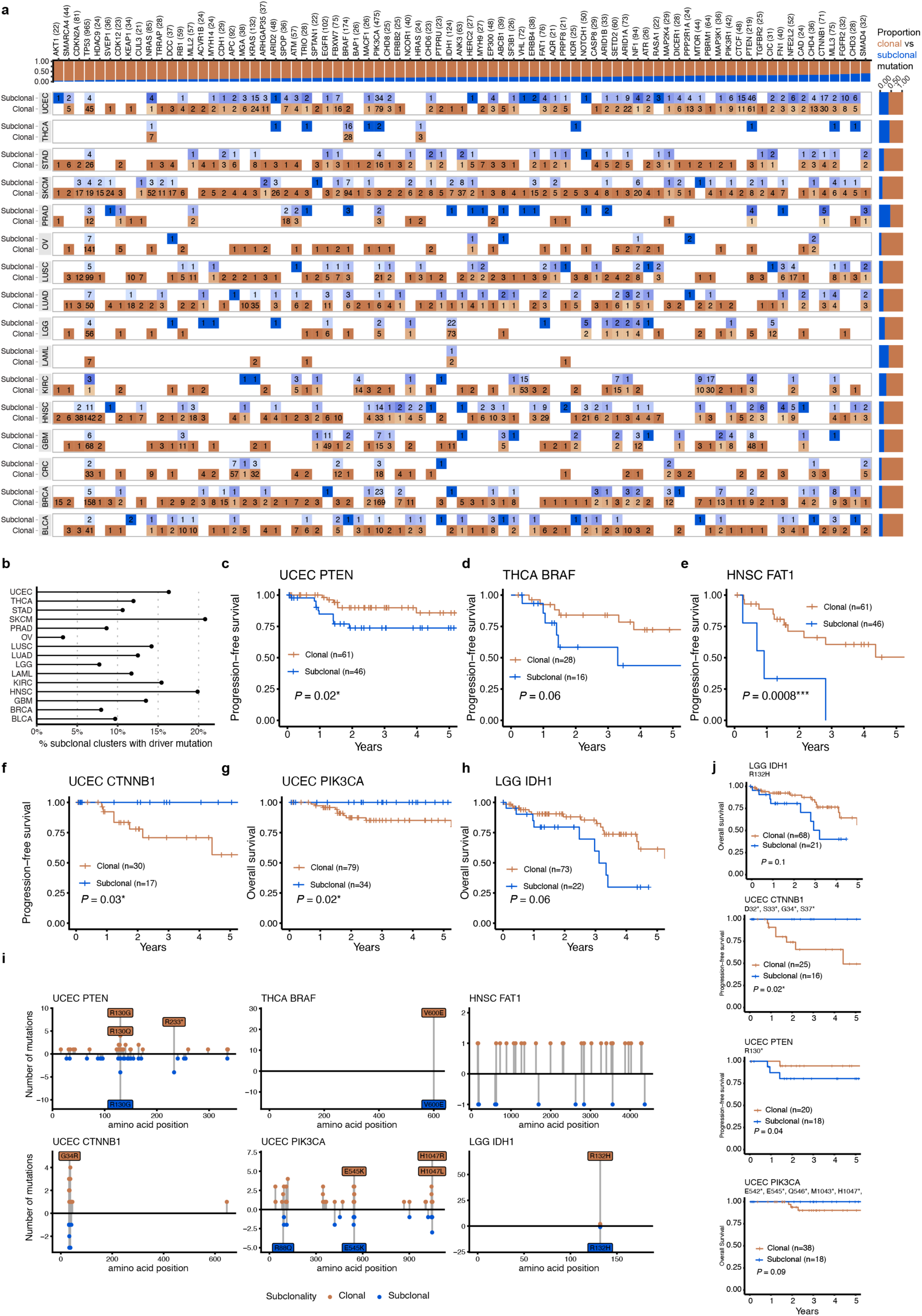
Subclonal assignment of driver mutations across most frequently mutated genes in TCGA. **(a)** Heatmap depicting the distribution of clonal (orange squares) and subclonal (blue squares) driver single nucleotide variants (SNVs) across top 78 genes in 16 cancer types (see **Methods**). Orange denotes clonal and blue denotes subclonal mutations. Numbers within squares represent SNV counts. A darker shade represents a higher proportion of clonal or subclonal mutations. Proportions of subclonal and clonal mutations are summarized by gene (top) and cancer type (right). **(b)** Percentage of subclonal mutation clusters containing a driver mutation, by cancer type. **(c-h)** Kaplan-Meier (KM) curves showing pairs of driver gene and cancer type with statistically significant survival differences between those with clonal (orange) versus subclonal (blue) driver mutations of the gene. Log-rank test *P*-values are shown. Significance levels are denoted as follows: \**P* < 0.05, \*\**P* < 0.01 and ***P < 0.001. **(i)** Lollipop plots showing mutation position distributions within each of the six significant cancer-driver gene pairs. Counts of clonal mutations in orange are indicated by the vertical height of the “lollipops”, while counts of subclonal mutations are shown in blue. Frequently recurrent mutations (n>4) are labeled. (**j**) KM curves showing progression-free survivals for patients with clonal and subclonal driver mutations restricting to the most recurrent sites for each cancer type – driver gene pair where applicable.

**Figure S6.**
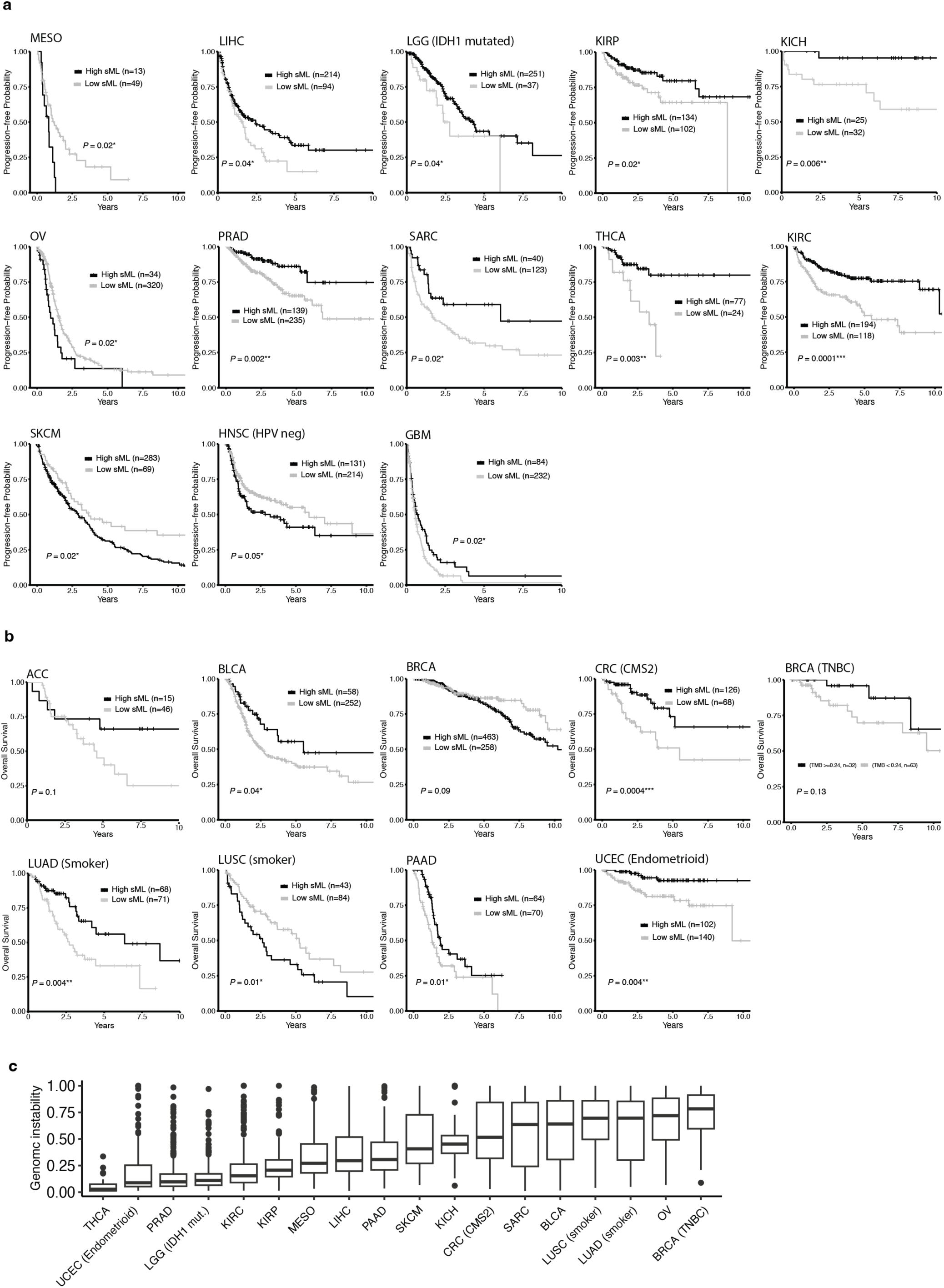
(**a**) KM curves of progression-free interval (PFI) for cancer types/subtypes shown in Figure 3c. Black lines denote high sML and grey lines denote low sML. *P*-values of log-rank tests between high- versus low-sML patient groups are shown. (**b**) KM curves of overall survival (OS) for cancer types/subtypes are shown in Figure 3c. Black lines denote high sML and grey lines denote low sML. *P*-values of log-rank tests between high- versus low-sML patient groups are shown. **(c)** Distributions of genome instability (GI) scores provided by ASCAT, a measure of the proportion of the genome displaying copy number aberrations, across cancer types featured in Figure 3c.

**Figure S7:**
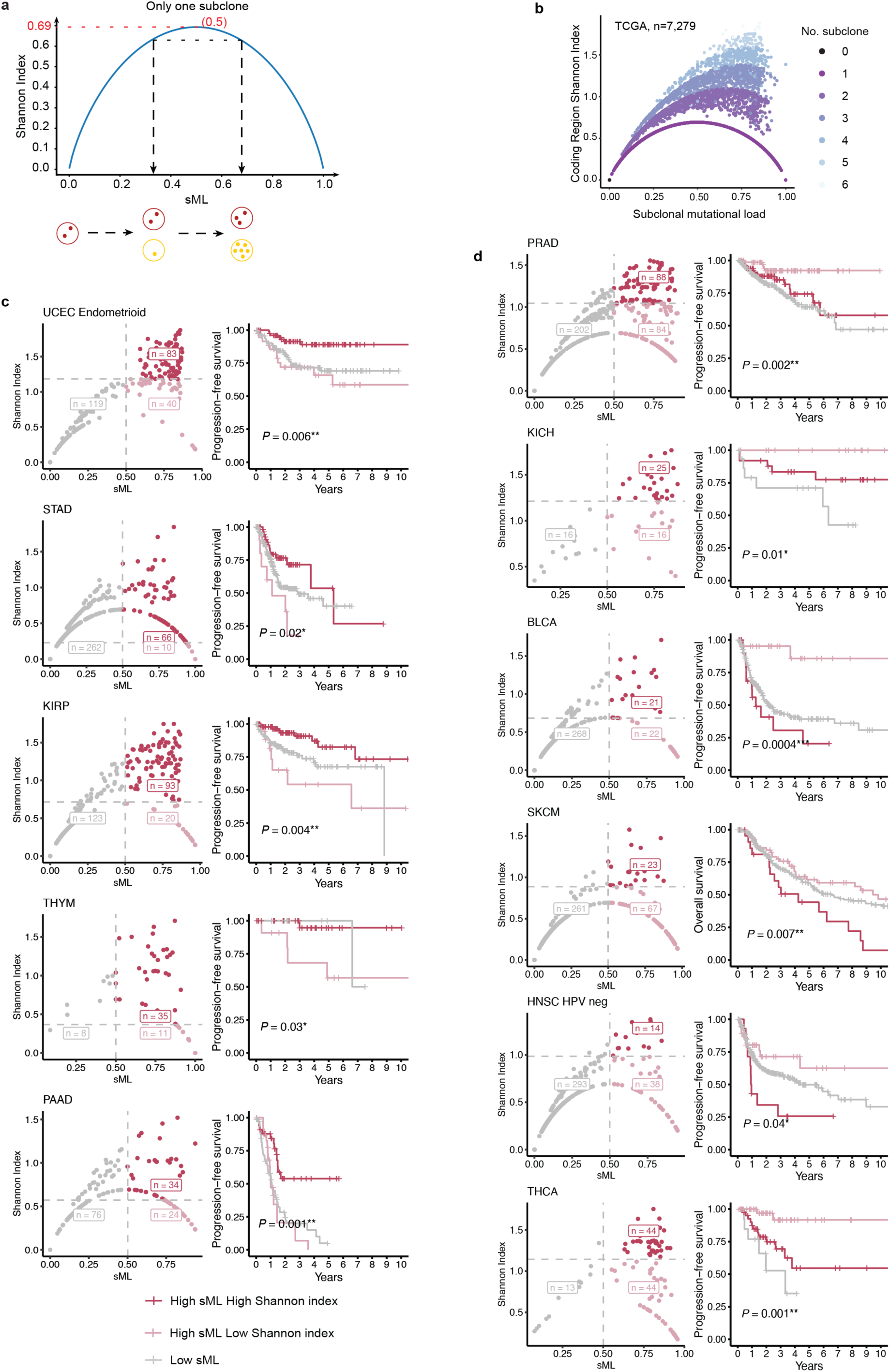
Subclonal diversity associated with subclonal mutation load. (**a**) Mathematical relationship between Shannon Index (SI) and sML using the simple case of one subclone as illustration. Two black dashed lines demonstrate that one subclone, despite sharing the same SI, can exhibit different sML values. The red dashed line denotes the highest SI score and the corresponding sML value of 0.5. **(b)** Scatter plot of SI versus sML in TCGA, where the color variations represent samples with different numbers of subclones. (**c-d**) Pairs of scatter plots displaying the relationship between SI,sML, and KM plots. Gray dots indicate low sML, while the red dots indicate high sML (sML > 0.5) with high SI, and the pink dots represent high sML with low SI. Additionally, for each cancer type, using the same cutoffs shown in the scatter plots and the same color scheme, KM curves are shown illustrating PFI or OS differences between the three groups. Log-rank test *P*-values are provided for comparing the patient groups. Cancer types are grouped as follows: (**c**) Cancer types where high sML and high SI present favorable outcomes (UCEC endometrioid, STAD, KIRP, THYM, PAAD) with statistical significance. (**d**) Cancer types where high sML and low SI present favorable outcomes (PRAD, KICH, BLCA, SKCM, HNSC (HPV neg), and THCA) with statistical significance.

**Figure S8:**
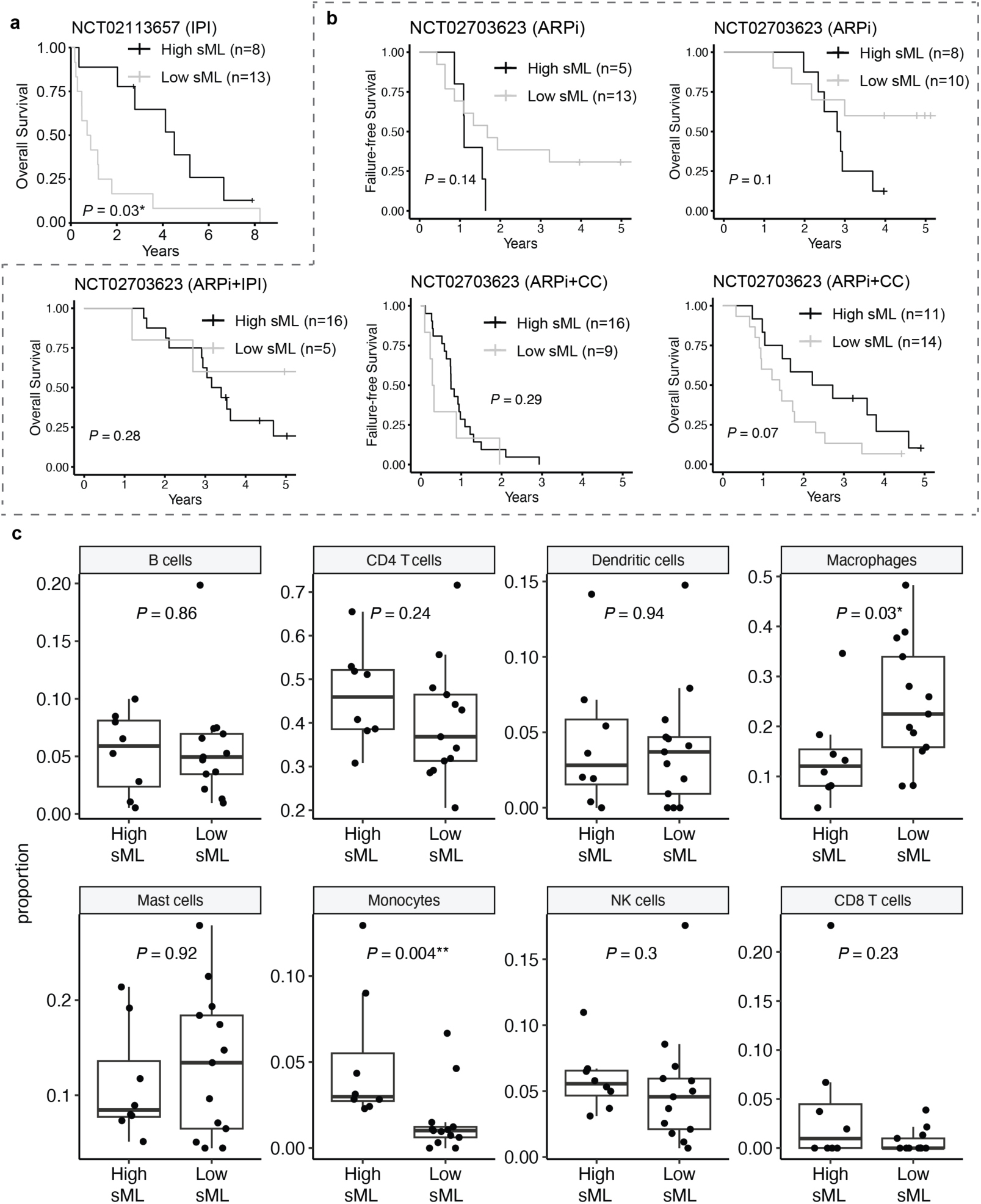
Clinical and immune associations of sML in mCRPC samples from two clinical trials. (**a**) K-M plot showing overall survival (OS) in patients from the IPI monotherapy trial stratified by sML, with log-rank test *P*-value shown. (**b**) K-M plots showing failure-free survival and overall survival (OS) in patients from all three treatment arms of the DynAMo trial stratified by sML, with log-rank test *P*-value shown. Patients were treated with androgen receptor pathway inhibitors (ARPi), ARPi with ipilimumab (IPI), or ARPi with carboplatin and cabazitaxel (CC). (**c**) Boxplots showing the distributions of immune cell types, estimated from CIBERSORTx (using LM22 as reference), between samples from IPI monotherapy trial with high and low sML (n=8 vs. 13). Major immune cell types with non-zero pooled proportion estimates are shown. *P*-value of the two-sided Wilcoxon rank sum test is shown. Significance levels are denoted as follows: \**P* < 0.05, \*\**P* < 0.01 and ***P < 0.001.

**Figure S9:**
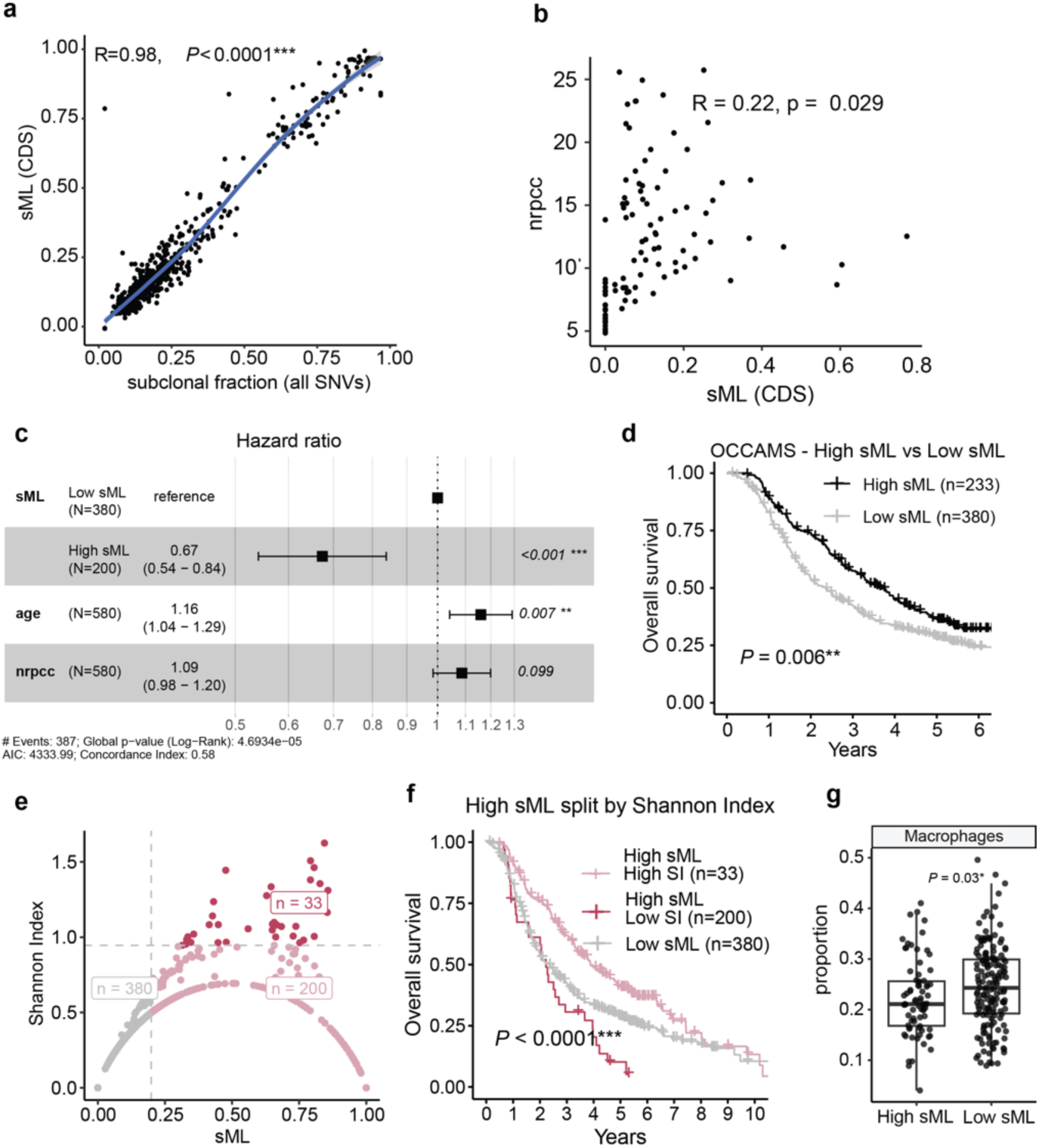
Analyses of the OCCAMS esophageal adenocarcinoma samples. **(a)** Scatter plot demonstrating a strong correlation between sML (of the coding region) and subclonal fraction (on all SNVs) in OCCAMS dataset, suggesting that either estimate can be used in downstream analyses. For consistency with the TCGA results, we use CDS sML for all subsequent analysis of the OCCAMS ESAD samples. (**b**) Scatter plot of nrpcc versus sML in OCCAMS, with Pearson correlation and *P*-value shown. **(c)** Forest plot showing hazard ratios from a Cox proportional hazard model of OCCAMS samples including of sML, nrpcc, and age. This suggests nrpcc does not explain the clinical association presented by sML. **(d)** KM plot showing overall survival difference between high sML and low sML for all 613 OCCAMS esophageal adenocarcinoma samples. **(e)** Scatter plot displaying the relationship between Shannon Index and sML. Gray dots indicate low sML, while the red dots indicate high sML with high SI, and the pink dots represent high sML with low SI, which has a small sample size of 33. (**f**) KM curves illustrating OS differences between low sML, high sML & low SI, and high sML & high SI. Log-rank test *P*-value is shown for comparing the patient groups. (**g**) Boxplot showing the distributions of macrophages estimated from CIBERSORTx (using LM22 as reference), between OCCAMS samples with available RNAseq with high and low sML (n=75 vs. 163). *P*-value of the two-sided Wilcoxon rank sum test is shown. Significance levels are denoted as follows: \**P* < 0.05, \*\**P* < 0.01 and ***P < 0.001.

## Notes

https://bioinformatics.mdanderson.org/apps/CliPP

### Summary of Updates

Title, additional data and modified conclusion accordingly.

