## Supplementary Information for "Subclonal mutation load predicts survival and response to immunotherapy in cancers with low to moderate tumor mutation burden"

### 1. Performance benchmarking using simulated datasets

As the true subclonal structure is difficult to obtain in real data, we assess the ability of CliPP to correctly reconstruct the subclonal organization of a tumor on three simulated datasets: PhylogicSim500[1], SimClone1000[1], and a newly generated dataset which we call CliPPSim4k.

#### 1.1 Metrics for benchmarking subclonal reconstruction accuracy

We evaluate how accurately each method recovers subclonal architecture using three metrics that measure bias in estimated number of clusters, fraction of clonal mutations (clonal fraction), and cellular prevalence (CP) across all variants, respectively.

- Measuring error in estimated number of clusters (rdNC): We calculate the relative difference in number of clusters $rdNC = \frac{|N_e - N_t|}{N_t}$, where $N_e$ is the estimated number of clusters and $N_t$ the true number of clusters.
- Measuring error in clonal fraction estimates (rdCF): We calculate the relative difference in clonal fraction $rdCF = \frac{|C_e - C_t|}{C_t}$, where $C_e$ and $C_t$ represent the CliPP estimated clonal fraction and the truth clonal fraction, respectively.
- Measuring error in CP estimates (RMSE): We further calculate the root mean squared error $RMSE = \sqrt{\frac{1}{N}\sum_{n=1}^{N}\left(\frac{\widehat{\phi_i} - \phi_i}{\rho}\right)^2}$, where $N$ is the total number of SNVs, $\widehat{\phi_i}$ is the estimated CP, $\phi_i$ is the true CP for SNV $i$, and $\rho$ is the true purity of the sample which was used to standardize the dynamic ranges of different samples.
- Measuring overall error: We also introduce the total error score = $\frac{rdNC + rdCF + RMSE}{3}$, to represent the overall performance.

In all these metrics, smaller values indicate better performance, and 0 indicates a correct reconstruction. To facilitate cross method comparison, we use min-max normalization to force the values to a [0,1] scale. The difference in normalized scores is calculated as $\frac{2(m_c - m_p)}{m_c + m_p}$, where $m_c$ and $m_p$ are scaled scores for CliPP and PhyloWGS respectively. A negative normalized difference score suggests better performance in CliPP.

#### 1.2 CliPPSim4k

We utilize our own simulation framework to include samples that are relatively silent in terms of copy number events and at a higher read coverage to comprehensively evaluate the subclonal reconstruction accuracy of CliPP. The simulation covers a range of values for key features that are known to influence accuracy, including tumor purity, percent genome with copy number alterations (CNA), sequencing read depth and the number of mutation clusters (**Extended Data Fig. 2a, Methods**). We call this dataset CliPPSim4k. We first set the following sample-level parameters: tumor purity $\rho$ as 0.3, 0.6, or 0.9, CNA rate as 0, 0.1, or 0.2, average read depth $N$ as 100, 500, or 1,000, and the number of clusters of SNVs (with unique CCF) $K$ , as 2, 3, or 4. The CCF for the $k$-th cluster of SNVs, denoted by $\beta_k$, is sampled from Uniform$(0.2, 1)$ with a minimum distance $d = 0.2$ between any two clusters. We then have CP $\phi_k = \rho\beta_k$ for the $k$-th cluster. The number of SNVs in each cluster is simulated from Uniform$(200, 500)$. To generate copy number status at each SNV, we take the following steps:

1. Initialize the status of mutated/reference allele of SNV at 1/1.
2. Sample status of SNV (1, 0 for with or without CNA, respectively) from Bernoulli$(p)$ with values of $p \in \{0,0.1,0.2\}$.
3. For each SNV with CNA, sample the number of copies of mutated allele and reference allele from a discrete uniform distribution over $\{0,1,2,3,4,5\}$

For each SNV $i$ , we sampled the total number of reads $n_i \sim$ Poisson$(N)$, where $N$ is the average read depth for the sample, taking values $N \in \{100,\ 500,\ 1{,}000\}$. We then sample the variant reads $r_i$ from Binomial$(n_i,\ \theta_i)$ , where $\theta_i$ is calculated from the purity, CP, and SNV-specific copy number using equation (1). We generated 50 samples for each combination of read depth, purity, CNA rate, and number of clusters. This gives us a total of $50 \times 81 = 4{,}050$ samples.

We compare the performance of CliPP with PhyloWGS[2] on the CliPPSim4k set of 4,050 simulated samples and find that overall CliPP performs slightly better than PhyloWGS, with a smaller total error score (**Fig. 2a**, median difference in total error score = -0.01, two-sided Wilcoxon test *P*-value <0.001). Within and across each feature combination (81 combinations in total), both methods are mostly comparable, with their accuracy more negatively impacted by high percent CNA and low purity than high cluster numbers (**Supplementary Figs. 1** and **2**, **Table S1**). Looking further into aspects of subclonal structure that are estimated, CliPP is slightly better at recovering the fraction of clonal mutations than PhyloWGS (median difference in rdCF is -0.04, two-sided Wilcoxon test *P*-value < 0.001), while performing comparably in recovering the number of clusters and the cellular prevalence.

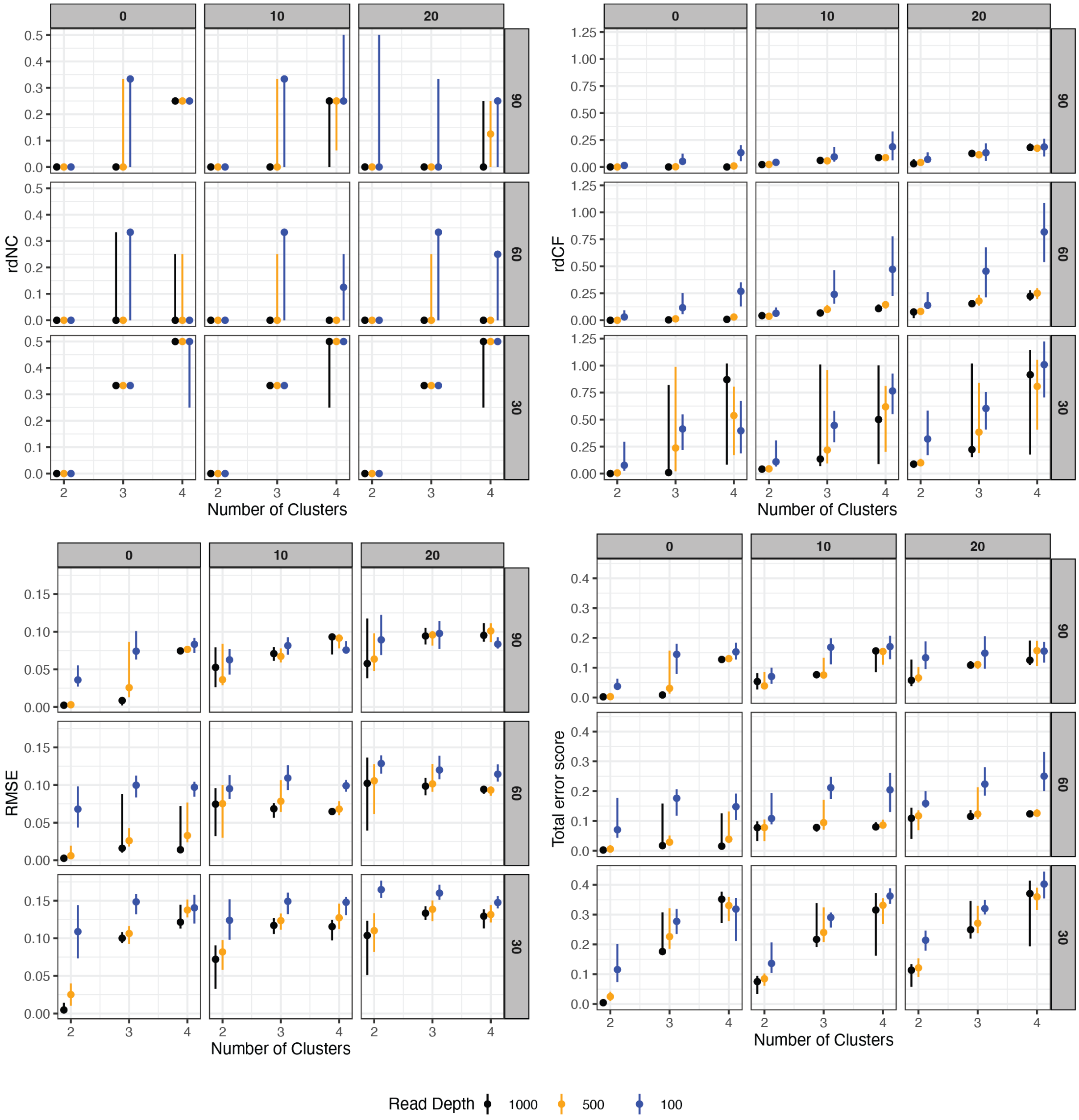

**Supplementary Fig. 1: Simulation results for CliPP in CliPPSim4k.** The "ball-stick" plots are utilized to display the performance of CliPP, where the ball represents the median value, and the upper and lower reaches of each stick denote the 25th and 75th percentiles, respectively. Smaller values in all metrics indicate better performance, with 0 suggesting an accurate reconstruction. (a) rdNC scores for all simulated scenarios. (b) rdCF scores for all simulated scenarios. (c) RMSE scores for all simulated scenarios. (d) Total error scores for all simulated scenarios. See **Table S1** for further information.

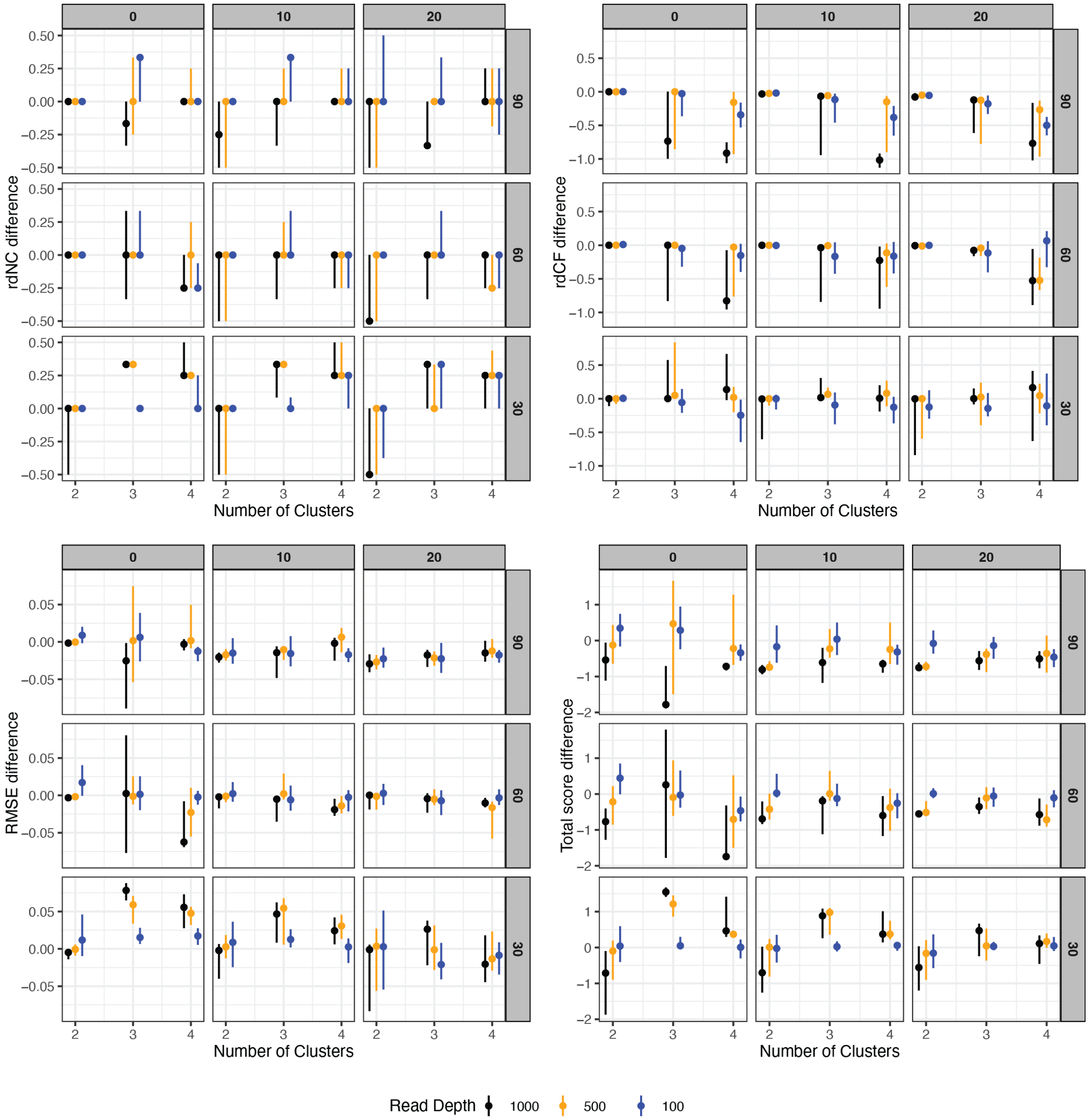

**Supplementary Fig. 2: Simulation results comparing CliPP with PhyloWGS in CliPPSim4k.** "Ball-stick" plots display the performance comparison between CliPP and PhyloWGS. A negative normalized difference score implies that CliPP outperforms PhyloWGS. (a) rdNC score differences between CliPP and PhyloWGS for all simulated scenarios. (b) rdCF score differences between CliPP and PhyloWGS for all simulated scenarios. (c) RMSE score differences between CliPP and PhyloWGS for all simulated scenarios (d) Normalized overall score differences between CliPP and PhyloWGS for all simulated scenarios.

| | | The means and standard deviations of rdNC for CliPP at K = 2 | | | The means and standard deviations of rdCF for CliPP at K = 2 | | | The means and standard deviations of RMSE for CliPP at K = 2 | | |
|---|---|---|---|---|---|---|---|---|---|---|
| Purity | CNA rate | Read Depth | | | Read Depth | | | Read Depth | | |
| | | 100 | 500 | 1000 | 100 | 500 | 1000 | 100 | 500 | 1000 |
| 0.3 | 0 | 0.01 (0.07) | 0 (0) | 0 (0) | 0.22 (0.29) | 0.02 (0.03) | 0.00 (0.01) | 0.11 (0.05) | 0.03 (0.02) | 0.01 (0.01) |
| | 0.1 | 0 (0) | 0 (0) | 0 (0) | 0.23 (0.24) | 0.06 (0.05) | 0.05 (0.04) | 0.12 (0.04) | 0.08 (0.03) | 0.06 (0.03) |
| | 0.2 | 0 (0) | 0 (0) | 0 (0) | 0.36 (0.23) | 0.12 (0.07) | 0.08 (0.04) | 0.16 (0.03) | 0.11 (0.04) | 0.09 (0.04) |
| 0.6 | 0 | 0.1 (0.20) | 0 (0) | 0 (0) | 0.11 (0.19) | 0.01 (0.01) | 0.00 (0.00) | 0.07 (0.04) | 0.01 (0.01) | 0.00 (0.00) |
| | 0.1 | 0.09 (0.19) | 0 (0) | 0 (0) | 0.16 (0.25) | 0.04 (0.02) | 0.04 (0.02) | 0.10 (0.03) | 0.07 (0.04) | 0.07 (0.03) |
| | 0.2 | 0.08 (0.19) | 0.01 (0.07) | 0 (0) | 0.25 (0.25) | 0.08 (0.04) | 0.07 (0.04) | 0.13 (0.02) | 0.10 (0.04) | 0.09 (0.05) |
| 0.9 | 0 | 0.07 (0.18) | 0 (0) | 0 (0) | 0.03 (0.08) | 0.01 (0.02) | 0.00 (0.00) | 0.04 (0.02) | 0.01 (0.01) | 0.00 (0.00) |
| | 0.1 | 0.09 (0.24) | 0 (0) | 0 (0) | 0.08 (0.13) | 0.03 (0.02) | 0.02 (0.02) | 0.06 (0.03) | 0.05 (0.03) | 0.06 (0.03) |
| | 0.2 | 0.19 (0.30) | 0.01 (0.07) | 0.03 (0.12) | 0.11 (0.12) | 0.04 (0.03) | 0.04 (0.04) | 0.10 (0.04) | 0.07 (0.03) | 0.08 (0.05) |
| | | The means and standard deviations of rdNC for CliPP at K = 3 | | | The means and standard deviations of rdCF for CliPP at K = 3 | | | The means and standard deviations of RMSE for CliPP at K = 3 | | |
| Purity | CNA rate | Read Depth | | | Read Depth | | | Read Depth | | |
| | | 100 | 500 | 1000 | 100 | 500 | 1000 | 100 | 500 | 1000 |
| 0.3 | 0 | 0.31 (0.09) | 0.33 (0.05) | 0.31 (0.08) | 0.43 (0.31) | 0.52 (0.50) | 0.32 (0.50) | 0.15 (0.02) | 0.10 (0.02) | 0.10 (0.02) |
| | 0.1 | 0.3 (0.10) | 0.33 (0.05) | 0.33 (0) | 0.47 (0.28) | 0.51 (0.47) | 0.50 (0.51) | 0.15 (0.02) | 0.12 (0.02) | 0.12 (0.01) |

| | | | | | | | | | | |
|---|---|---|---|---|---|---|---|---|---|---|
| | 0.2 | 0.32 (0.07) | 0.32 (0.07) | 0.33 (0.05) | 0.60 (0.30) | 0.54 (0.43) | 0.53 (0.45) | 0.16 (0.03) | 0.14 (0.02) | 0.13 (0.02) |
| 0.6 | 0 | 0.21 (0.18) | 0.08 (0.14) | 0.13 (0.16) | 0.19 (0.20) | 0.02 (0.02) | 0.01 (0.01) | 0.10 (0.02) | 0.04 (0.03) | 0.04 (0.04) |
| | 0.1 | 0.23 (0.16) | 0.09 (0.15) | 0.03 (0.10) | 0.34 (0.26) | 0.11 (0.06) | 0.08 (0.04) | 0.11 (0.02) | 0.08 (0.02) | 0.07 (0.02) |
| | 0.2 | 0.18 (0.17) | 0.09 (0.15) | 0.05 (0.12) | 0.47 (0.31) | 0.18 (0.08) | 0.16 (0.05) | 0.12 (0.02) | 0.11 (0.02) | 0.10 (0.02) |
| 0.9 | 0 | 0.23 (0.23) | 0.11 (0.16) | 0.05 (0.12) | 0.09 (0.09) | 0.01 (0.02) | 0.00 (0.01) | 0.08 (0.02) | 0.04 (0.04) | 0.02 (0.03) |
| | 0.1 | 0.28 (0.25) | 0.1 (0.15) | 0.05 (0.12) | 0.14 (0.12) | 0.05 (0.02) | 0.06 (0.02) | 0.09 (0.02) | 0.07 (0.02) | 0.07 (0.01) |
| | 0.2 | 0.15 (0.20) | 0.03 (0.10) | 0.01 (0.07) | 0.20 (0.22) | 0.11 (0.04) | 0.12 (0.03) | 0.10 (0.02) | 0.09 (0.02) | 0.09 (0.01) |
| | | The means and standard deviations of rdNC for CliPP at K = 4 | | | The means and standard deviations of rdCF for CliPP at K = 4 | | | The means and standard deviations of RMSE for CliPP at K = 4 | | |
| Purity | CNA rate | Read Depth | | | Read Depth | | | Read Depth | | |
| | | 100 | 500 | 1000 | 100 | 500 | 1000 | 100 | 500 | 1000 |
| 0.3 | 0 | 0.41 (0.12) | 0.47 (0.09) | 0.48 (0.07) | 0.45 (0.32) | 0.52 (0.36) | 0.68 (0.49) | 0.14 (0.02) | 0.13 (0.02) | 0.13 (0.02) |
| | 0.1 | 0.46 (0.10) | 0.45 (0.10) | 0.42 (0.12) | 0.75 (0.37) | 0.53 (0.33) | 0.56 (0.49) | 0.14 (0.02) | 0.13 (0.02) | 0.12 (0.02) |
| | 0.2 | 0.49 (0.06) | 0.46 (0.09) | 0.42 (0.12) | 0.98 (0.37) | 0.74 (0.39) | 0.75 (0.52) | 0.15 (0.01) | 0.13 (0.02) | 0.13 (0.02) |
| 0.6 | 0 | 0.06 (0.11) | 0.09 (0.14) | 0.08 (0.13) | 0.30 (0.22) | 0.03 (0.03) | 0.01 (0.01) | 0.09 (0.02) | 0.05 (0.03) | 0.03 (0.03) |
| | 0.1 | 0.13 (0.14) | 0.04 (0.09) | 0.03 (0.08) | 0.51 (0.33) | 0.15 (0.07) | 0.12 (0.06) | 0.10 (0.01) | 0.07 (0.02) | 0.07 (0.01) |
| | 0.2 | 0.15 (0.15) | 0.03 (0.08) | 0.01 (0.04) | 0.82 (0.39) | 0.25 (0.07) | 0.23 (0.07) | 0.12 (0.02) | 0.09 (0.01) | 0.09 (0.01) |
| 0.9 | 0 | 0.25 (0.16) | 0.23 (0.08) | 0.23 (0.08) | 0.16 (0.14) | 0.02 (0.02) | 0.00 (0.00) | 0.08 (0.02) | 0.07 (0.02) | 0.07 (0.02) |

| | | | | | | | | | | |
|---|---|---|---|---|---|---|---|---|---|---|
| | 0.1 | 0.27 (0.19) | 0.19 (0.11) | 0.18 (0.11) | 0.20 (0.15) | 0.08 (0.03) | 0.09 (0.03) | 0.08 (0.02) | 0.09 (0.02) | 0.09 (0.02) |
| | 0.2 | 0.21 (0.19) | 0.13 (0.13) | 0.11 (0.13) | 0.22 (0.18) | 0.18 (0.05) | 0.18 (0.05) | 0.08 (0.02) | 0.10 (0.01) | 0.10 (0.01) |

**Table S1. CliPP performance benchmarking results in CliPPSim4k**. Means and standard deviations of rdNC, rdCF, and RMSE for CliPP across all simulated parameters in the CliPPSim4k dataset.

#### 1.3 PhylogicSim500 and SimClone1000 data

In Dentro et al.[1], the PhylogicSim500 was the legacy simulation set with which various subclonal reconstruction methods including PhyloWGS were trained. For independent testing, SimClone1000 was further generated in the study. The PhylogicSim500 dataset contains 500 samples, where the copy number profiles were sampled from the PCAWG WGS data and the other parameters were independently sampled from fixed distributions[1]. In this dataset, the tumor purity ranges from 0.16 to 0.99, the CNA rate (defined as the proportion of SNVs with copy number change across all SNVs) ranges from 0 to 0.96, the read depth ranges from 27 to 129, and the true number of mutation clusters range from 1 to 5 (**Extended Data Fig. 2b**). The SimClone1000 dataset, which was a validation dataset for PCAWG generated by SimClone[1] and contains 965 samples. In this dataset, the tumor purity ranges from 0.16 to 1, the CNA rate ranges from 0 to 1, the read depth ranges from 34 to 149, and the true number of clusters ranges from 1 to 8 (**Extended Data Fig. 2c**). We obtained PhyloWGS results on 469 samples in PhylogicSim500 and 911 samples in SimClone1000. We ran CliPP on the same samples to compare the performances. We find that overall both methods are comparable in both datasets (**Supplementary Fig. 3**). CliPP has a slightly higher total error score, median absolute difference in total error score = 0.01 (two-sided Wilcoxon test *P*-value < 0.001) for PhylogicSim500 and 0.002 (two-sided Wilcoxon test *P*-value < 0.001) for SimClone1000.

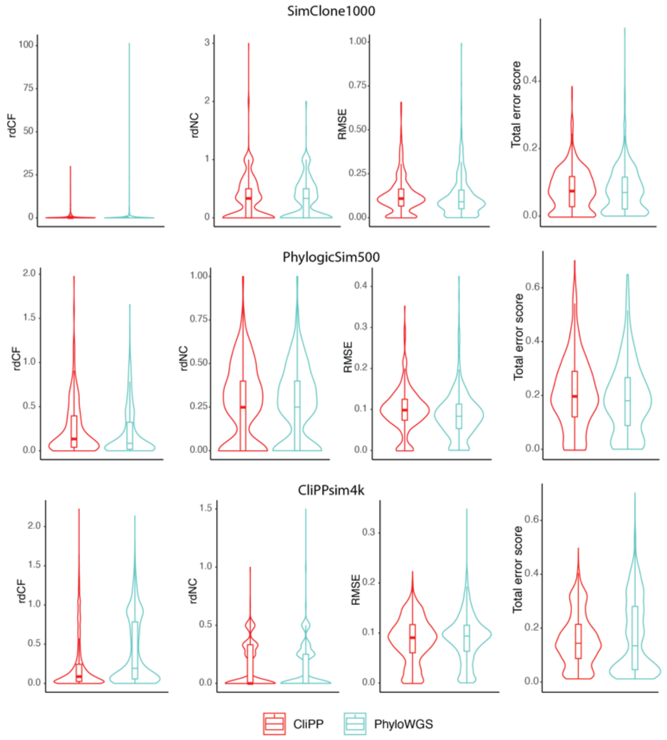

**Supplementary Fig. 3: Simulation results comparing CliPP with PhyloWGS across 3 simulated cohorts.** Performance benchmark comparisons between CliPP and PhyloWGS on PhylogicSim500, SimClone1000, and CliPPsim4k datasets. For PhylogicSim500, the median values for (CliPP, PhyloWGS) are: (0.25, 0.25) for rdNC, (0.13, 0.084) for rdCF, (0.099, 0.084) for RMSE. For SimClone1000, the median values are: (0.33, 0.33) for rdNC, (0.15, 0.15) for rdCF, (0.11, 0.091) for RMSE. For CliPPSim4k, the median values are: (0, 0) for rdNC, (0.088, 0.192) for rdCF, (0.091, 0.094) for RMSE.

#### 1.4 Benchmarking accuracy in PCAWG samples

We compared clustering results between CliPP and PyClone-VI using PCAWG samples with ≥10 the number of reads per tumor chromosomal copy (nrpcc). Concordance was evaluated using three metrics: Bangdiwala's B statistic[3,4] for subclonality classification (clonal versus subclonal), and Fowlkes-Mallows Index and purity for cluster assignment agreement. All metrics showed strong concordance (median >0.95) between the two methods (**Supplementary Fig. 4**). In addition to calculating the concordance correlation coefficients as shown in **Fig. 2a**, we further visualized the comparison of CliPP with 5 methods and performed linear regression, demonstrating the good concordance of CliPP with other high-ranking methods in terms of subclonal fraction (**Supplementary Fig. 5**).

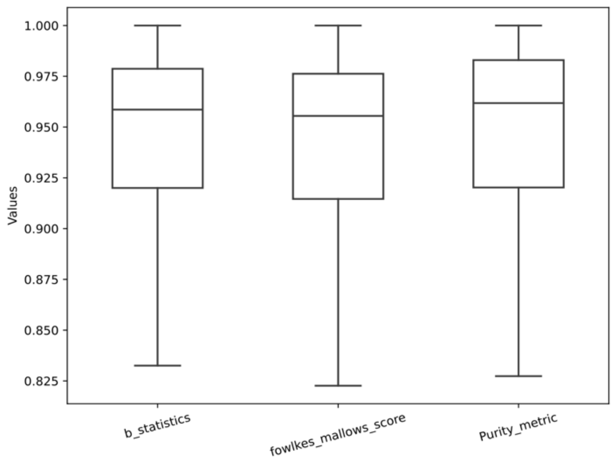

**Supplementary Fig. 4:** Boxplots illustrating the distributions of multiple concordance scores derived from pairwise comparisons of SNV cluster assignments between CliPP and PyClone-VI on the PCAWG data (n=1,985).

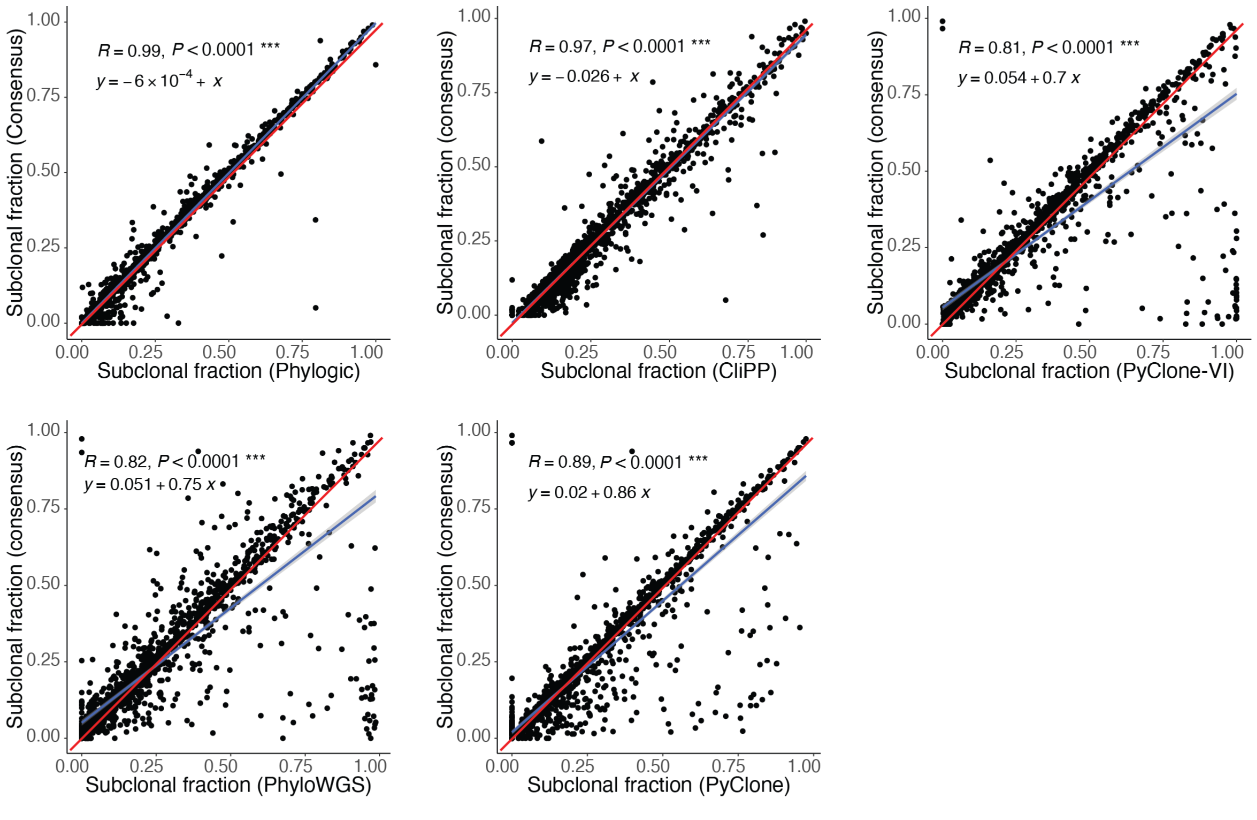

**Supplementary Fig. 5: Estimated subclonal fraction comparison to consensus subclonal fraction for PCAWG samples for 5 methods.** Scatter plots of subclonal fraction estimations from individual method versus consensus calls in PCAWG. Pearson correlation and *P*-values are shown, along with the equation for a fitted linear regression, which is plotted as a blue line, with the red line showing the diagonal.

### 2. Timing comparison among CliPP, PyClone-VI, and PhylogicNDT in real patient cohorts

CliPP consistently delivers superior efficiency across diverse datasets, significantly outperforming PyClone-VI[4], and PhylogicNDT[5]. Whether analyzing TCGA WES, OCCAMS WGS, or TCGA WGS samples, CliPP completes tasks in a fraction of the time required by other methods, enabling faster analysis, iterative refinements, and improved resource management for large-scale cancer genomics studies.

#### 2.1 TCGA WES cohort

To better compare the timing across CliPP, PyClone-VI[4], and PhylogicNDT[5], we obtained quartiles of the distribution of mutation counts in our TCGA cohort. Within each of the quartiles: <50, 50-94, 94-210, and 210-18,824, we compute the speed up ratio based on the recorded timing by CliPP and PyClone-VI. To project an average speed-up of over PhylogicNDT for the TCGA cohort, we took the time ratio of PhylogicNDT over PyClone-VI (1.9x) at SNV number=100 (**Extended Data Fig. 2e**), and multiplied it with the median ratio of 514x for PyClone-VI over CliPP.

#### 2.2 OCCAMS WGS dataset

Although not impossible, the entire process of subclonal reconstruction remains to be tedious and can be improved significantly. As an example, we evaluated the timing of analyzing WGS data from 613 esophageal samples that were released by OCCAMS consortium[6,7], as described further below.

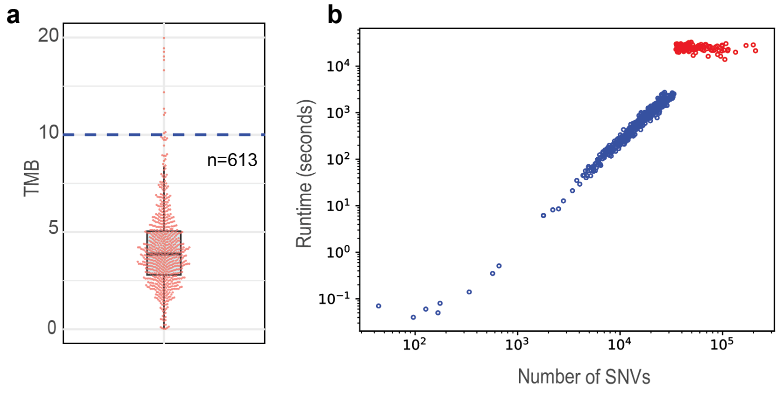

**Supplementary Fig. 6: Distributions of TMB and computational times for subclonal reconstruction using CliPP in OCCAMS.** (a) Boxplot showing the distribution of TMB for all OCCAMS samples. (b) Runtime for OCCAMS samples. Red and blue dots represent CliPP runs with and without a grid-based subsampling approach, respectively. The runtime was measured using the same configuration as the PCAWG analysis, running CliPP on an Intel® Xeon® Gold 6132 CPU @ 2.60GHz with 28 cores.

The majority of samples in this cohort exhibit low to moderate TMB (**Supplementary Fig. 6a)**. As shown in **Table S2**, CliPP delivered results quickly, processing a whole genome sequencing (WGS) sample between a few minutes and 20 minutes (**Supplementary Fig. 6b**) and completed the analysis for 400 samples presenting < 35,000 SNVs in about 2 hours. This efficiency significantly accelerates research efforts. For samples exceeding 35,000 SNVs, we employed subsampling with a size of 35,000 and repeated 10 times, hence the total runtime was about 10 times longer as represented by the red dots in **Supplementary Fig. 6b**.

| **Software** | **Timing** |
|---|---|
| CliPP Shiny app | Seconds per sample for clustering and visualization |
| CliPP | 2 hours for 400 samples |
| PhyloWGS | Approximately 1200 hours for 400 samples (by projection) |

**Table S2.** Timing of subclonal reconstruction in OCCAMS on Intel(R) Xeon(R) Gold 6132 CPU @ 2.60GHz.

#### 2.3 An example speed evaluation using 100 WGS samples from TCGA

While speed may not always be the primary concern in cancer genomics research, CliPP's efficiency offers substantial practical benefits in certain contexts. To demonstrate this, we evaluated the timing of our recent analysis of WGS data from 100 Papillary Thyroid Cancer (PTC) samples in TCGA, as detailed below:

**Alignment and variant calling:** With our recently published accelerated mutation calling approach, achieved by a consensus of Strelka2[8] and MuSE 2[9] and fully automated through a Snakemake pipeline[10], it took us a total of **345.92 hours (2 weeks)** to obtain reliable mutation calls from 100 raw BAM files (on Intel(R) Xeon(R) Gold 6132 CPU @ 2.60GHz). In a benchmark study, we estimated a 45x speed up of this pipeline with a higher accuracy for WGS data, as compared to running MuTect2[11]. Currently, there are common concerns for the speed of mutation calling being overwhelmingly slow as many research labs run MuTect2. However, we consider the issues of cost-efficient alignment and variant calling solvable as several good solutions have been provided recently by the sequencing community (e.g., bwa-mem2, bwa-meme-lisa, the DRAGEN server made by Illumina Inc, Nvidia’s Parabricks gpu-accelerated mutect2) as well as from our group.

**Subclonal reconstruction using CliPP:** As shown in **Table S3**, CliPP generates results within seconds for a sample and 30 minutes for all 100 samples, providing significant advantages for expediting research. Furthermore, subclonal reconstruction is never finalized after one single run. There are various nuanced investigations of input and output data that would lead to a rerun with modified configurations. Servers can stall or stop big jobs unexpectedly. We are seeing ~10 reruns for an experienced bioinformatician who can pre-empt most common mistakes. This is affordable by CliPP but not so much by other methods: e.g. 125 days for finalizing subclonal reconstruction by PhyloWGS. Hence the contribution of CliPP (reduce to 30min) to minimize the overhead time expenditure of a cancer genomic study becomes significant, as it allows for a substantial amount of time to be used for manual data curation which is still much needed in future studies.

| **Software** | **Timing** |
|---|---|
| CliPP Shiny app | Seconds per sample for clustering and visualization |
| CliPP | 30 minutes for 100 samples |
| PhyloWGS | Approximately 300 hours for 100 samples (by projection) |

**Table S3.** Timing of subclonal reconstruction in 100 WGS PTC samples on Intel(R) Xeon(R) Gold 6132 CPU @ 2.60GHz.

### 3. Benchmarking subclonal reconstruction using an intersection of WGS and WES data

Whole exome sequencing surveys only the coding part of the cancer genome but alternatively provides a significantly higher read depth as compared to WGS data at the same cost. Using tumor samples that had both WES and WGS data generated from the same tissue sample as part of the TCGA and PCAWG consortia, we compared CliPP subclonal reconstruction results to provide pan-cancer WES benchmark statistics as a new resource and to assess the utility of performing large-scale subclonal reconstruction using WES data.

Before comparing results at the SNV level, we removed a subset of samples with significantly elevated TMB, as these hypermutator samples would otherwise have an oversized influence on our agreement measures, hence confounding the benchmarking study. Therefore, we removed these outlier samples from the dataset, i.e. the 3$^{rd}$ quartile+1.5x interquartile range (IQR) of all samples for each cancer type. The cohort comprised 488 samples with 124,725 SNVs before filtering, and 447 samples and a total of 49,554 SNVs after filtering.

Across a total of 49,554 mutations from 447 samples that were called in both WES and WGS data, the median read depth in WES is 1.4 fold higher than in WGS (WES median = 82, WGS median = 57, two-sided Wilcoxon test *P*-value < 0.001, **Table S4**). We evaluated agreement in the CliPP-based clonal/subclonal assignment of the same mutations from the WES and WGS data using Bangdiwala's B statistic[3]. For interpretation of the degree of agreement, we used thresholds defined by Muñoz and Bangdiwala[11], which follow the benchmarking scale introduced by Landis and Koch[12]. Using these predefined categories, we observed no samples to have poor agreement, with most samples displaying high agreement between WGS and WES subclonal reconstruction results: 85% of samples having “substantial” to “perfect” agreement, and the median B-statistic for all samples (median = 0.85) lies in the “almost perfect” range. The remaining 68 samples (15%) show fair or moderate agreement ($0.49>B\geq0.09$). When ranking cancer types by their median B-statistic, we find a moderate but not entirely consistent trend in which tumors with higher TMB present higher WES/WGS concordance (Pearson correlation = 0.5). Across all samples, there is a consistent trend in more clusters being called in WES and more mutations changing identity from being clonal in WGS to subclonal in WES, as compared to the opposite direction (McNemar’s test *P*-values < 0.001, **Extended Data Fig. 4b).** Mutations shifting from WGS-clonal to WES-subclonal are expected when sequenced at a higher read coverage, hence less likely to be clustering errors. These mutations dominate the disagreement in the fair and moderate agreement samples (1,754 out of 1,907, 92%).

**Table S4. Read coverage evaluation at shared mutations between WES and WGS data (also see Extended Data Fig. 4d).** Consistent with our expectation, the sequencing coverage in TCGA datasets was substantially higher, averaging a 1.5-fold increase. Notably, certain cancer types, including GBM, KICH, and KIRC, exhibited more pronounced disparities (>2.5x). Surprisingly, SARC samples constituted an anomaly with a median sequencing depth that was reduced relative to PCAWG WGS samples. Note the read coverages were evaluated at the called mutation sites, as compared to the whole genome or exome.

| Cancer type | Median read depth (WES) | Median read depth ratio (WES/WGS) |
|---|---|---|
| BLCA | 73 | 1.84 |
| BRCA | 87 | 1.69 |
| CESC | 63 | 1.19 |
| CRC | 100 | 2.11 |
| DLBC | 95.5 | 1.28 |
| GBM | 102 | 2.47 |
| HNSC | 82 | 1.34 |
| KICH | 88 | 1.57 |
| KIRC | 120 | 2.36 |
| KIRP | 94.5 | 1.69 |
| LGG | 77 | 1.82 |
| LIHC | 91 | 1.50 |
| LUAD | 80 | 1.55 |
| LUSC | 90 | 1.34 |
| OV | 133 | 2.20 |
| PRAD | 90 | 1.61 |
| SARC | 47 | 0.89 |
| SKCM | 63 | 1.17 |
| STAD | 81 | 1.32 |
| THCA | 79 | 1.19 |
| UCEC | 81 | 1.67 |

In addition to higher read depth leading to identification of additional subclonal mutations hence lowering agreement in some samples, we found several technical factors independent of the subclonal reconstruction procedure were additional contributors to disagreement between WGS and WES. These factors are further described below and in **Supplementary Fig. 7**.

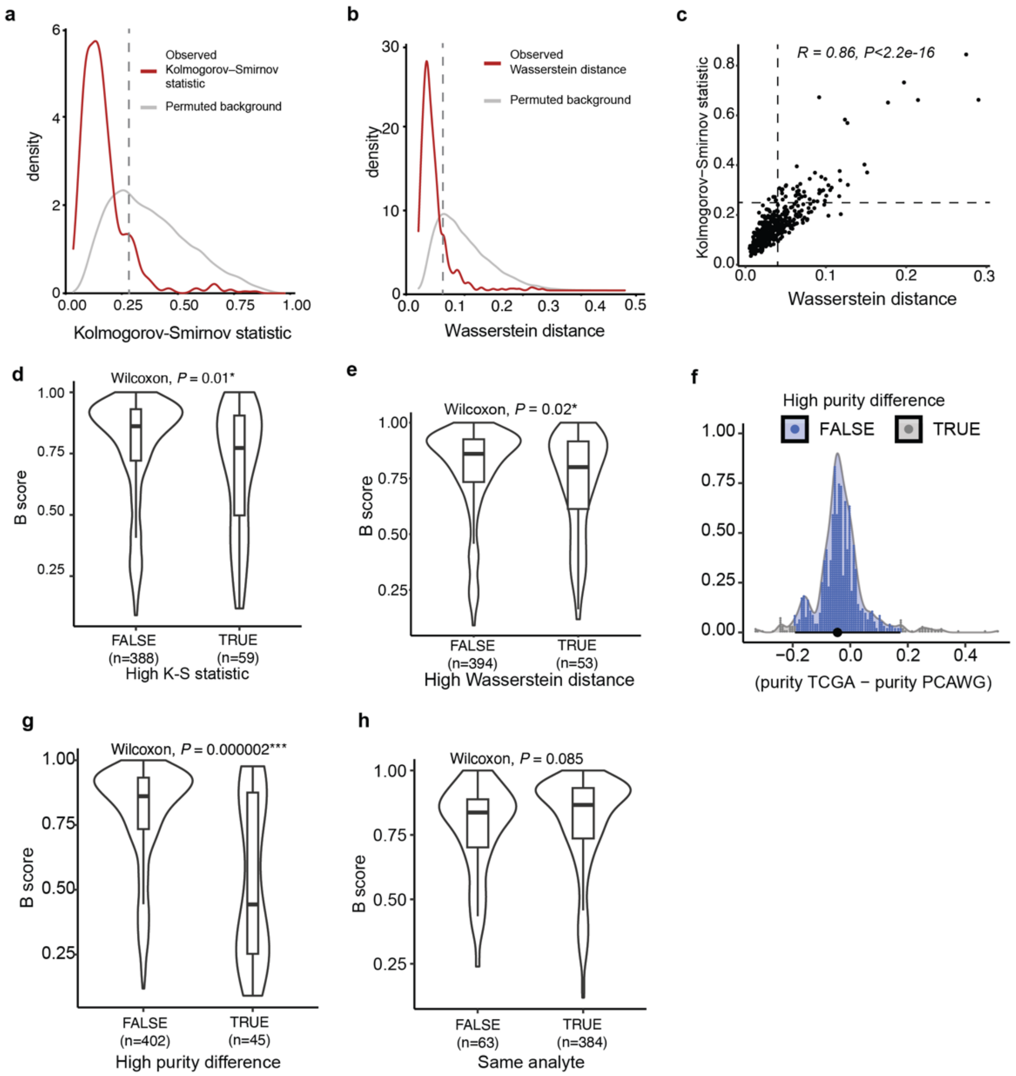

**Supplementary Fig. 7: Technical factors contributing to disagreement between TCGA and PCAWG samples. (a)** Observed Kolmogorov-Smirnov (K-S) statistic distribution between VAF distributions of WGS and WES pairs (red), with a permuted background distance distribution (grey) computed by comparing random WGS and WES samples (see **Methods**). A smaller K-S statistic indicates good consistency in the VAF distributions. Cutoff for “high K-S statistic” is shown as a vertical dotted line. **(b)** Observed Wasserstein distance distribution between VAF distributions of WGS and WES pairs (red), with a permuted background distance distribution (grey) computed by comparing random WGS and WES samples. A smaller Wasserstein distance indicates good consistency in the VAF distributions. Cutoff for “high Wasserstein distance” is shown as a vertical dotted line. **(c)** Scatter plot of K-S statistic values and Wasserstein distances between VAF distributions for all 447 samples. **(d)** Violin and boxplots showing the difference in B score between samples with high K-S statistic, with the Wilcoxon rank-sum test *P*-value shown. **(e)** Violin and boxplots showing the difference in B score between samples with high Wasserstein distance, with the Wilcoxon rank-sum test *P*-value shown. **(f)** Distribution of differences in sample purity estimates from TCGA and PCAWG. Dots represent individual sample comparisons, with the top and bottom 5% of samples indicated by gray color. **(g)** Violin and boxplots showing the difference in B score between samples with extreme differences (top and bottom 5% in panel (**f**)) in purity estimates, with the Wilcoxon rank-sum test *P*-value shown. **(h)** Violin and boxplots showing the difference in B score between samples with identical sample preparations compared to samples that derived from separate analytes, with the Wilcoxon rank-sum test *P*-value shown.

When the raw input data are vastly different, then even the identical subclonal reconstruction procedure will lead to different results. We assessed the similarity of the raw input data by calculating the Wasserstein distance[13,14] and Kolmogorov-Smirnov (K-S) test statistic between the WGS and WES variant allele frequency (VAF) distributions. When the observed Wasserstein distance or K-S test statistic were higher than the median background distance, as calculated by permutation, samples tended to have lower agreement (around 10% of all samples, **Supplementary Figs 7a-e**).

Different CNA profiles account for many of the low agreement samples. We used consensus copy number profiles for PCAWG samples, and ASCAT copy number profiles from SNP6 array for TCGA. While these results generally show high concordance, there are some samples where the copy number profiles are different, hence CliPP produced a different subclonal structure for these samples. In these overlapping TCGA/PCAWG samples, we can identify samples with drastically different copy number profiles by examining the difference in purity estimates between WGS and WES data. When the purity estimates are significantly different, the CNA profile solution differs for the samples as well. When we compare B scores between samples with high purity differences (top and bottom 5%, **Supplementary Fig. 7f**) to the rest of the samples, we observe much lower agreement in samples with large purity differences **(Supplementary Fig. 7g),** highlighting the importance of quality copy number profiles in subclonal reconstruction.

Additionally, by comparing sample barcodes we confirmed that 98% of the matched WGS and WES samples indeed came from the same sample portion. However, among these, some (n=63, 14%) came from separate analytes, with the WGS sample being from a standard DNA extraction, while the WES sample analyte field in the barcode indicates they were obtained via a whole genome amplification produced using Repli-G (Qiagen) DNA. We find that these samples show lower agreement than those with the same barcode, i.e., same analyte **(Supplementary Fig. 7h).**

Lastly, we found no difference in agreement based on nrpcc.

As more studies are developed to investigate the clinical implications of subclonality and intratumoral heterogeneity, it is imperative for researchers to employ data-driven approaches in their study design. Two major findings from our benchmarking WES/WGS based subclonal reconstruction include 1) WES-based subclonal reconstruction is reliable and may identify more subclonal coding mutations in cancers with low TMB; and 2) when designing WGS experiments for cancers with low TMB, increasing read depth is an important consideration, as more subclonal mutations are likely to be identified.

### 4. Subclonal landscape of driver mutations

Driver mutations are genetic alterations that contribute to (or “drive”) the development and progression of cancer[38]. Much effort has been made to identify and catalogue these mutations from the background “passenger” mutations by identifying signals of positive selection in these mutations[39]. Due to their role in tumor initiation and the fitness advantage they confer, most driver mutations are clonal. However, these same driver mutations can occur within subclones and confer a selective advantage later in tumor evolution[12]. Using a robustly annotated set of 8,586 driver mutations in 574 genes across 2,469 samples from 16 cancer types[40], we find that 1,757 (20.5%) driver mutations were subclonal (**Extended Data Fig. 5a**). Overall, driver mutations were reported in 12% of subclonal mutation clusters in TCGA, in line with the 11% observed in PCAWG[12] **(Extended Data Fig. 5b**).

Studies of specific single cancer types have found mixed evidence for survival being associated with clonal and subclonal drivers[41,42], yet this question has not been addressed across cancer types in a comprehensive manner. Out of all driver gene/cancer-type combinations with 9 or more samples bearing subclonal mutations (a total of 16 pairs, **Supplementary Table 1**), we find 4 combinations (*PTEN, CTNNB1*, and *PIK3CA* in UCEC, *FAT1* in HNSC) to present significant association with prognosis along with *BRAF* in THCA and *IDH1* in LGG which are marginally significant (**Extended Data Fig. 5c-h**). When limited to the most frequently occurring mutation sites, such as *BRAF* V600E, *PTEN* mutations to R130, the observed trends with PFS or OS remain significant for *CTNNB1* and *PTEN* in UCEC, with other showing marginal signficance (**Extended Data Figs. 5i-j**). These findings, our pan-cancer large-scale subclonal annotation using CliPP, have provided new insights into the potential impact of the timing of driver mutations on patient outcomes.

Sequence similarities often lead to ambiguous alignment, i.e., reads mapping to regions with high similarity to other genomic regions will be randomly aligned across them. This can skew the VAF of a mutation in such a region, leading to false subclonal assignment. As described in **Methods**, we compared the distribution of mapping quality scores between clonal and subclonal mutations using two-sided Wilcoxon rank sum tests and found evidence for significantly lower mapping quality scores in only 2 genes (*MLL3* and *NCOR1*, **Supplementary Fig. 8**).

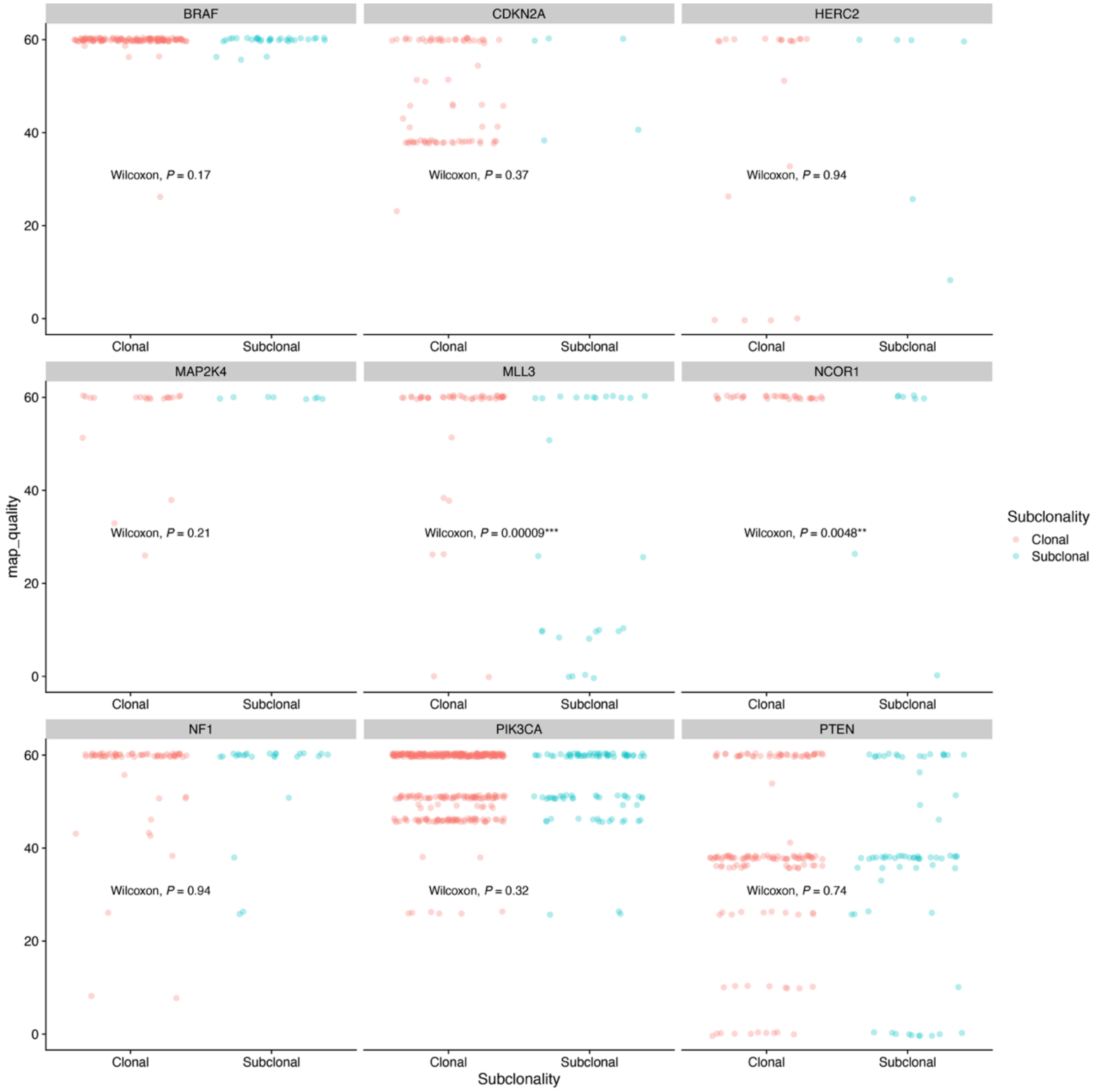

**Supplementary Fig. 8. Potential false subclonal identification caused by poor read mapping.** Scatterplots of the mapping quality scores for all mutations in 9 driver genes that have non-perfect mapped reads. Two-sided Wilcoxon rank tests compare the distributions of mapping quality scores between clonal (red) and subclonal (blue) mutations.

### 5. Evaluation of confounders in sML association analysis in TCGA

In this section, we aim to underscore the robustness of sML as a biomarker for studying tumor survival outcomes, by showing its independence from potential biological and technical confounders.

#### 5.1 sML versus multifaceted patient variables

We observed no significant associations between sML and age, sex, TMB, or clinical stage (**Supplementary Figs. 9-12**), suggesting a unique role of sML as a clinically relevant feature.

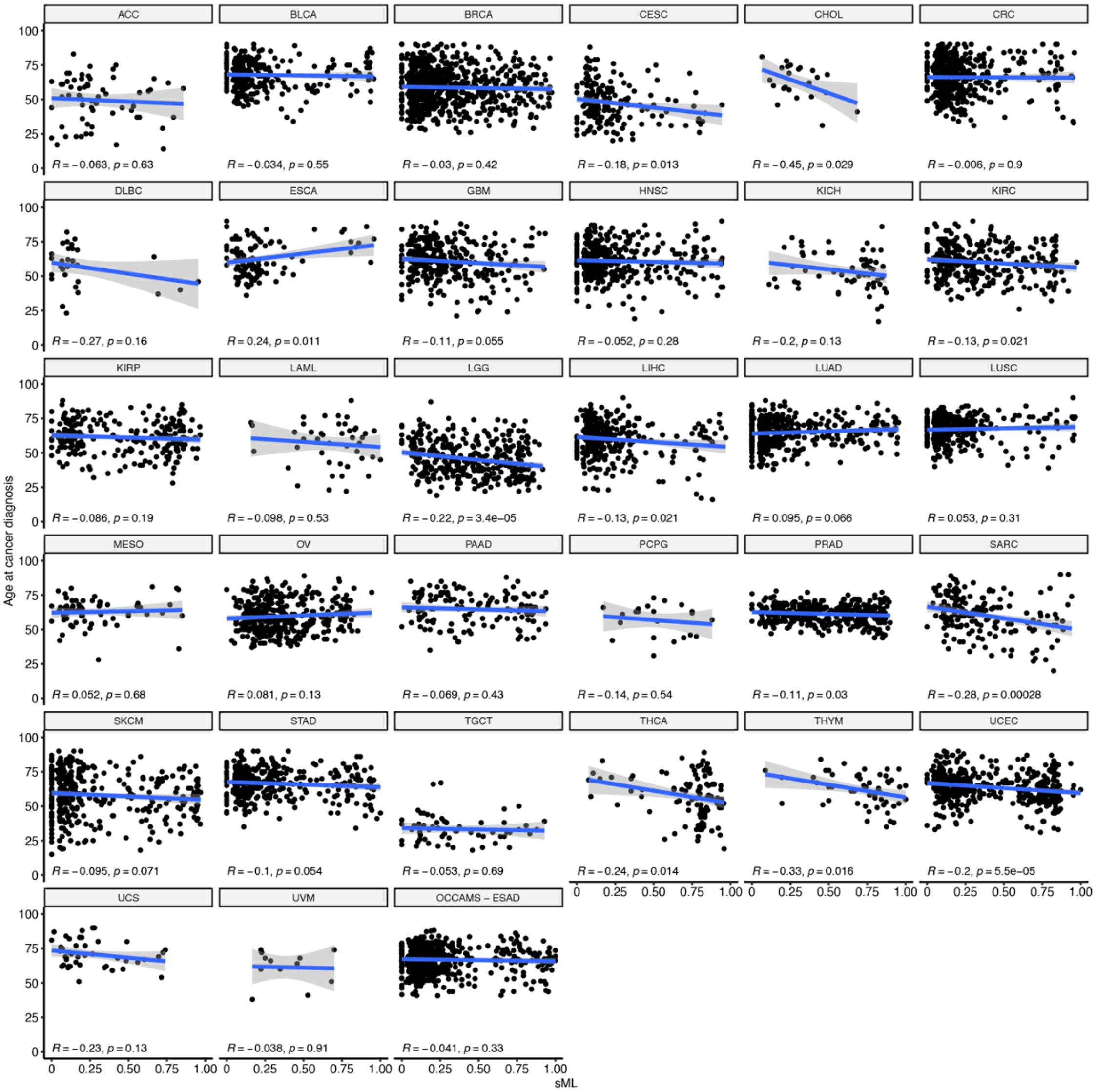

**Supplementary Fig. 9. Subclonal mutational load (sML) is not associated with age of diagnosis.** Scatter plots of CliPP estimated sML vs. patient age at cancer diagnosis from each of the 32 cancer types from TCGA, plus OCCAMS esophageal adenocarcinoma samples. The Pearson correlation coefficient R’s are shown for each cancer type.

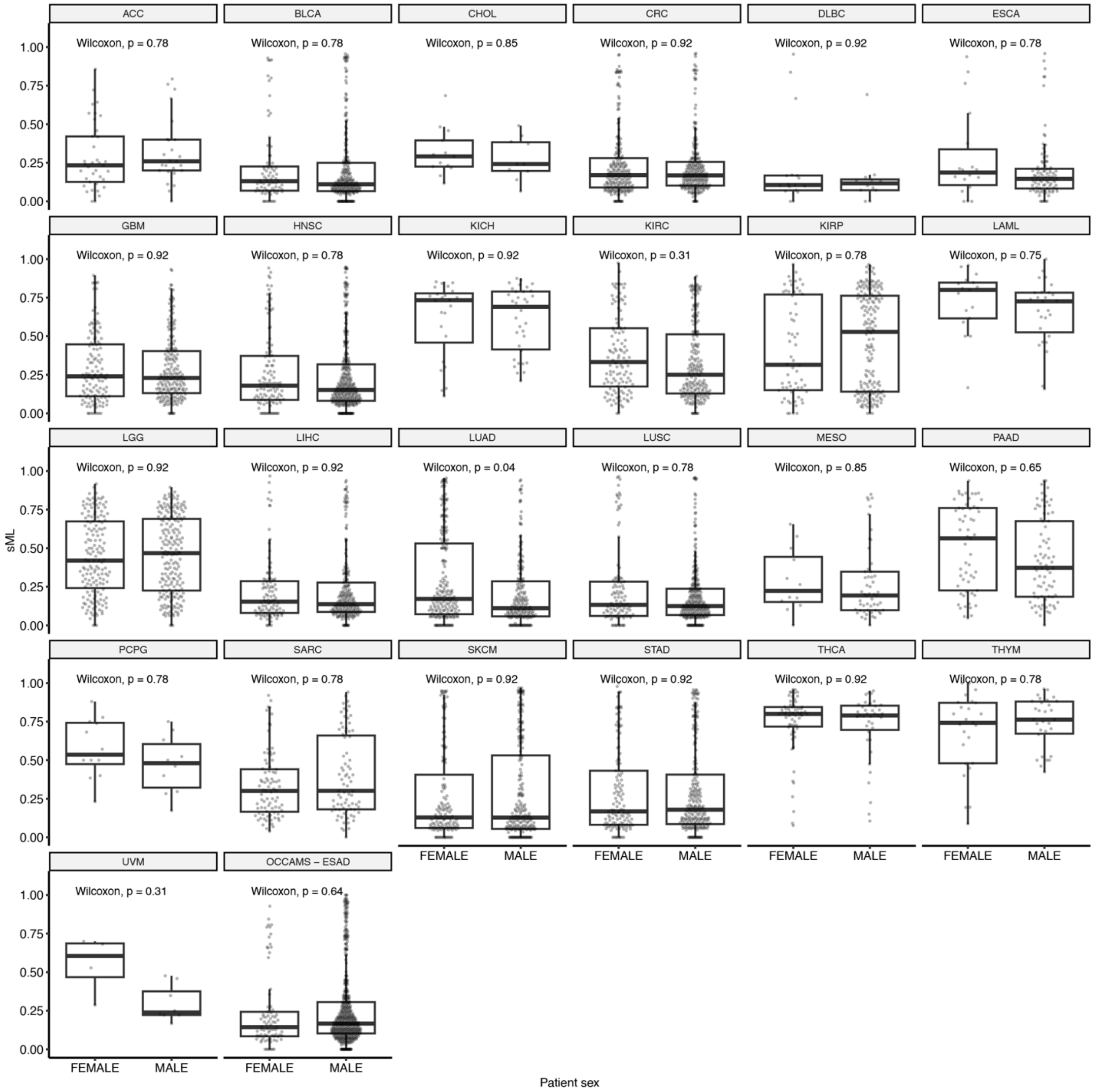

**Supplementary Fig. 10. Subclonal mutational load (sML) is not associated with sex.** Distribution of CliPP estimated sML for female and male patients across 25 cancer types, excluding sex-specific cancers, plus OCCAMS esophageal adenocarcinoma samples. Only one (LUAD) of the BH-adjusted *P*-values of two-sided Wilcoxon rank-sum tests reached significance at a confidence level of 0.05.

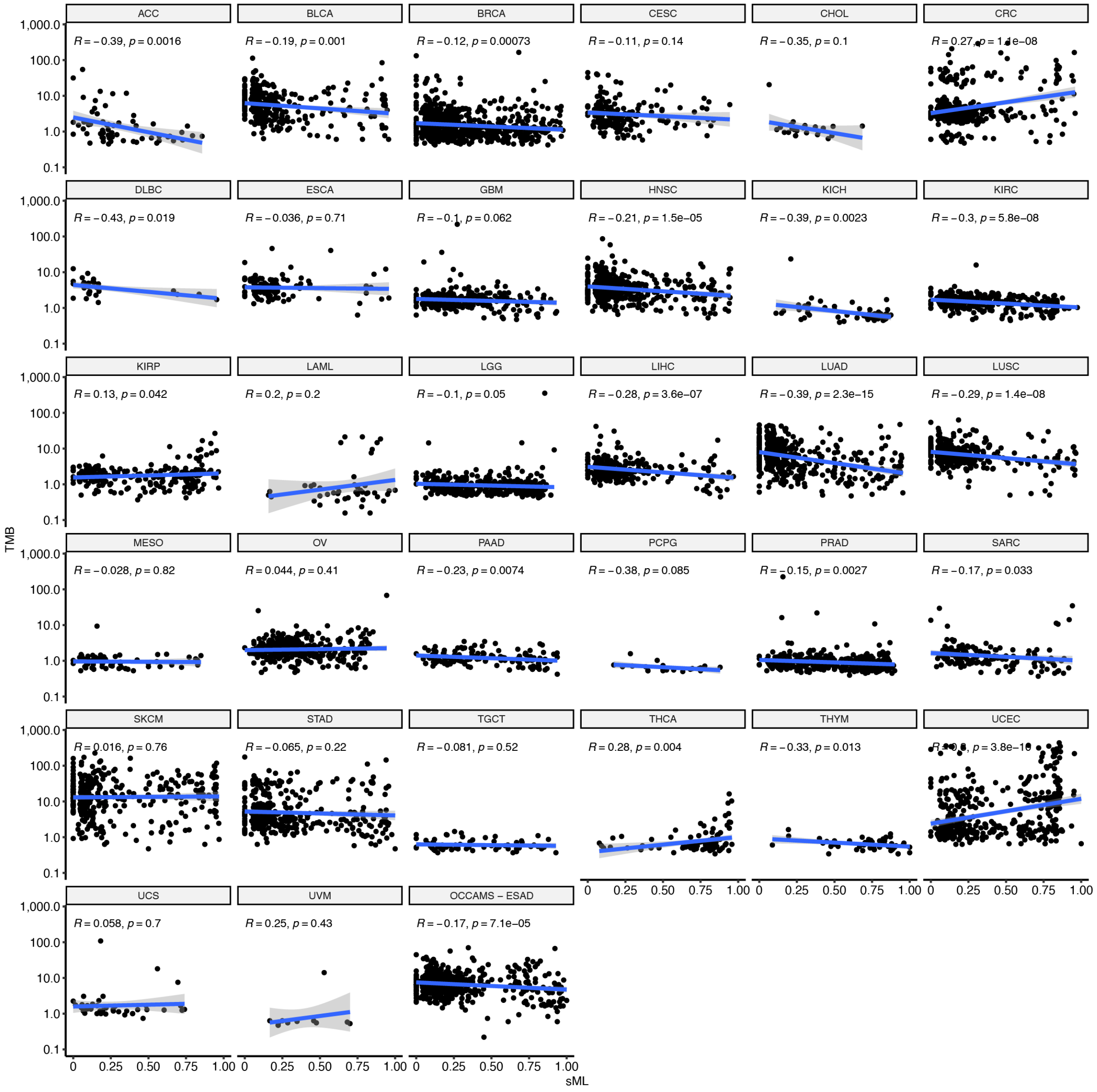

**Supplementary Fig. 11. Subclonal mutational load (sML) is not associated with TMB.** Scatter plots of CliPP estimated sML vs. TMB from each of the 32 cancer types, plus OCCAMS ESAD samples. The Pearson correlation coefficient Rs are shown for each cancer type.

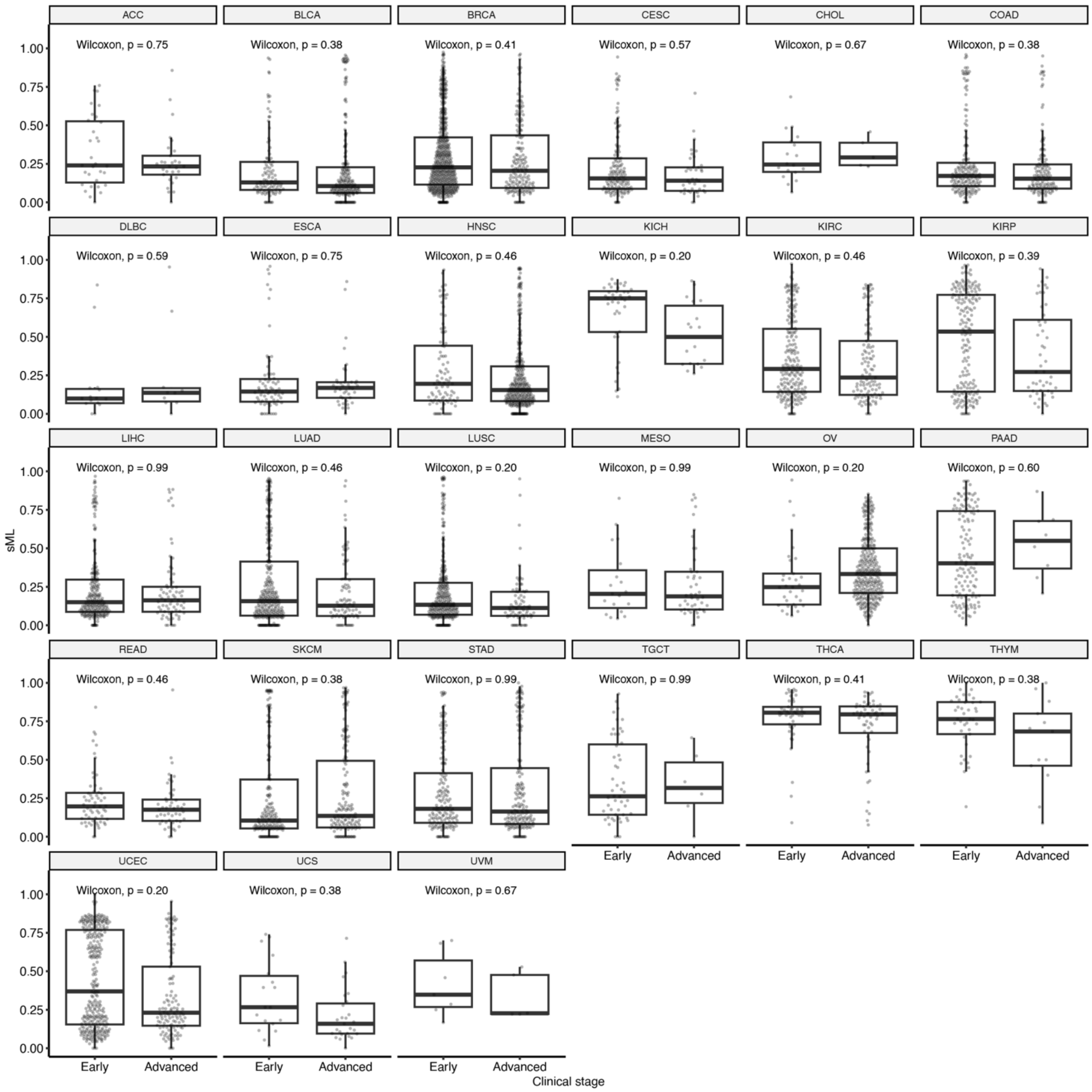

**Supplementary Fig. 12. Subclonal mutational load (sML) is not associated with clinical/pathological stage.** Scatter plots of CliPP estimated sML and the clinical stage which was dichotomized into early and advanced groups. Patients across 27 cancer types have clinical stage information. None of the *P*-values of two-sided Wilcoxon rank-sum tests reached significance at a confidence level of 0.05.

We further summarized the weak and mostly negative correlations between sML and TMB across cancer types in **Table S5**. This weak and slightly negative correlation mitigates concerns regarding collinearity and suggests that sML captures a distinct aspect of the tumor genomic landscape and that while TMB quantifies the number of mutations, sML still reflects a different dimension of tumoral changes, providing unique insights into the tumor biology. To account for potential technical biases caused by tumor purity, ploidy, and sequencing coverage, we also included the number of reads per tumor chromosomal copy (nrpcc) into our analysis. Note that nrpcc represents the sequencing coverage per haploid genome, and as nrpcc increases, the signal of true CCF peaks becomes clearer relative to read-sampling noise, and clones are easier to be distinguished. In **Table S5**, we show the Pearson correlation between nrpcc and sML within each cancer type, which reveals mostly weak to moderate associations (max Pearson correlation = 0.57 in TGCT). This gives more credibility to sML being a biological feature as compared to a technological feature. Lastly, we verified that higher sML is not caused by a larger tumor biopsy sample, which could in theory include more subclonal populations within the tumor. We compared the tumor portion weight with sML and found no significant correlations in any cancer type (**Table S5**).

**Table S5. Correlations of sML with TMB, nrpcc, and tumor weight in TCGA.**

| Cancer type | sML_TMB_Pearson_Correlation | sML_nrpcc_Pearson_Correlation | sML_Tumor_Portion_Weight_Pearson_Correlation |
|---|---|---|---|
| ACC | -0.27 | 0.15 | -0.06 |
| BLCA | -0.05 | 0.49 | 0.00 |
| BRCA | -0.03 | 0.32 | 0.02 |
| CESC | -0.12 | 0.28 | 0.08 |
| CHOL | -0.36 | 0.07 | -0.09 |
| CRC | 0.25 | 0.34 | 0.01 |
| DLBC | -0.39 | 0.32 | -0.04 |
| ESCA | 0.08 | 0.50 | 0.02 |
| GBM | -0.02 | 0.23 | 0.14 |
| HNSC | -0.10 | 0.34 | 0.03 |
| KICH | -0.27 | 0.31 | -0.04 |
| KIRC | -0.18 | 0.11 | -0.02 |
| KIRP | 0.22 | 0.25 | -0.02 |
| LAML | 0.20 | 0.10 | NA |
| LGG | 0.08 | 0.26 | 0.01 |
| LIHC | -0.10 | 0.29 | -0.03 |
| LUAD | -0.25 | 0.55 | -0.04 |
| LUSC | -0.14 | 0.47 | -0.01 |
| MESO | -0.05 | 0.22 | 0.14 |

| | | | |
|---|---|---|---|
| OV | 0.14 | 0.23 | 0.07 |
| PAAD | -0.17 | 0.51 | -0.09 |
| PCPG | -0.37 | 0.07 | 0.16 |
| PRAD | -0.07 | 0.29 | 0.10 |
| SARC | 0.07 | 0.24 | -0.02 |
| SKCM | 0.04 | 0.02 | 0.03 |
| STAD | 0.05 | 0.52 | -0.03 |
| TGCT | -0.09 | 0.57 | 0.18 |
| THCA | 0.24 | 0.06 | 0.00 |
| THYM | -0.35 | 0.42 | -0.09 |
| UCEC | 0.20 | 0.35 | 0.06 |
| UCS | -0.01 | 0.20 | -0.18 |
| UVM | 0.25 | 0.33 | -0.07 |

#### 5.2 Unique contribution of sML to stratifying patient survival

We divided samples within cancer types into high and low sML groups using rpart (see **methods**), and we used survival outcomes following the recommendations from the TCGA Pan-Cancer Clinical Data Resource (TCGA-CDR)[12]. Survival outcomes used and sML cutoff for dichotomizing sML are summarized in **(Table S6).**

**Table S6. Summary of clinical outcomes and cutoffs used to dichotomize sML as a biomarker.**

| Cancer type | Survival type | Cutoff |
|---|---|---|
| ACC | OS | 0.420 |
| BLCA | OS | 0.333 |
| BRCA | OS | 0.153 |
| BRCA_TNBC | OS | 0.238 |
| CRC_CMS2 | OS | 0.142 |
| LUAD_smoker | OS | 0.163 |
| LUSC_smoker | OS | 0.194 |
| PAAD | OS | 0.438 |
| UCEC_Endometrioid | OS | 0.645 |
| GBM | PFI | 0.407 |
| HNSC_HPVnegative | PFI | 0.197 |
| KICH | PFI | 0.737 |
| KIRC | PFI | 0.193 |
| KIRP | PFI | 0.314 |

| | | |
|---|---|---|
| LGG_IDH1 | PFI | 0.187 |
| LIHC | PFI | 0.095 |
| MESO | PFI | 0.413 |
| OV | PFI | 0.726 |
| PRAD | PFI | 0.610 |
| SARC | PFI | 0.513 |
| SKCM | PFI | 0.052 |
| THCA | PFI | 0.695 |
| ESAD (OCCAMS) | OS | 0.199 |
| mCRPC (NCT02113657) | OS, rcPFS | 0.500 |
| mCRPC (NCT02703623) | Failure-free survival (FFS) | 0.293 |

We further evaluated the unique contribution of sML to patient survival stratification. For all Cox proportional hazard models, we performed stepwise variable selection (see **methods**), with candidate variables including tumor sample purity, ploidy, and coverage. These technical factors were rarely selected in a powered and balanced Cox regression **(Supplementary Tables 3-4, 6).**

Adjusting for nrpcc also accounts for potential bias in sample generation, where more aggressive tumors may have been sequenced at higher depth. Therefore, we reconstructed the sML Cox regression model to include nrpcc as an optional variable. Upon performing stepwise variable selection, we observed that nrpcc does not retain significance when sML is present in the model (as demonstrated in **Supplementary Fig. 13** for KIRC and LUAD-smokers**).** This outcome corroborates the robustness of sML as an independent variable in modeling survival outcomes, not overshadowed by adjusted sequencing depth as measured by nrpcc.

We further show KM plots illustrating that sML's stratification of patient survival cannot be captured by TMB (as demonstrated in **Supplementary Fig. 14** for HPV-negative HNSC and LUAD smokers). Among LUAD-smokers, we are not able to stratify patient survival by either 1) ordering the samples by TMB and splitting to get a similar number in each group as the sML split, or 2) using rpart to iteratively search for a significant threshold to stratify patients using TMB. Similarly, for the HPV-negative HSNC dataset, ordering the samples according to TMB and splitting into groups the same size as the sML groups is not significant, and the rpart identified threshold is too extreme (only 3 samples in low TMB group). These results suggest that while weakly correlated, sML measures a unique feature of tumors that is not captured by TMB.

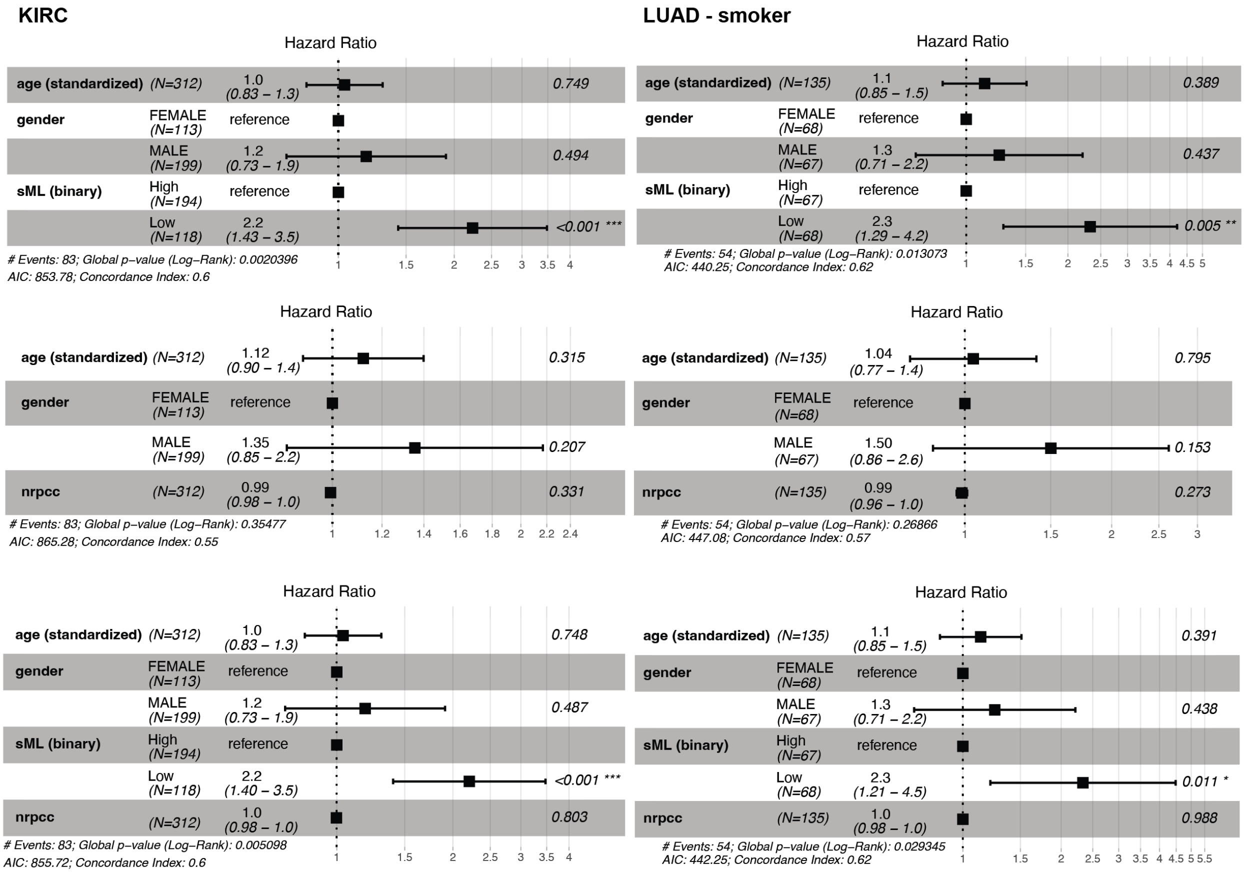

**Supplementary Fig. 13. Sequencing coverage measured by nrpcc cannot replace sML in association with clinical outcomes.** Adjusting by age and sex, we construct Cox regression models using either sML (binary) alone, nrpcc alone, or using both sML (binary) and nrpcc for KIRC and LUAD (smoker), as two examples.

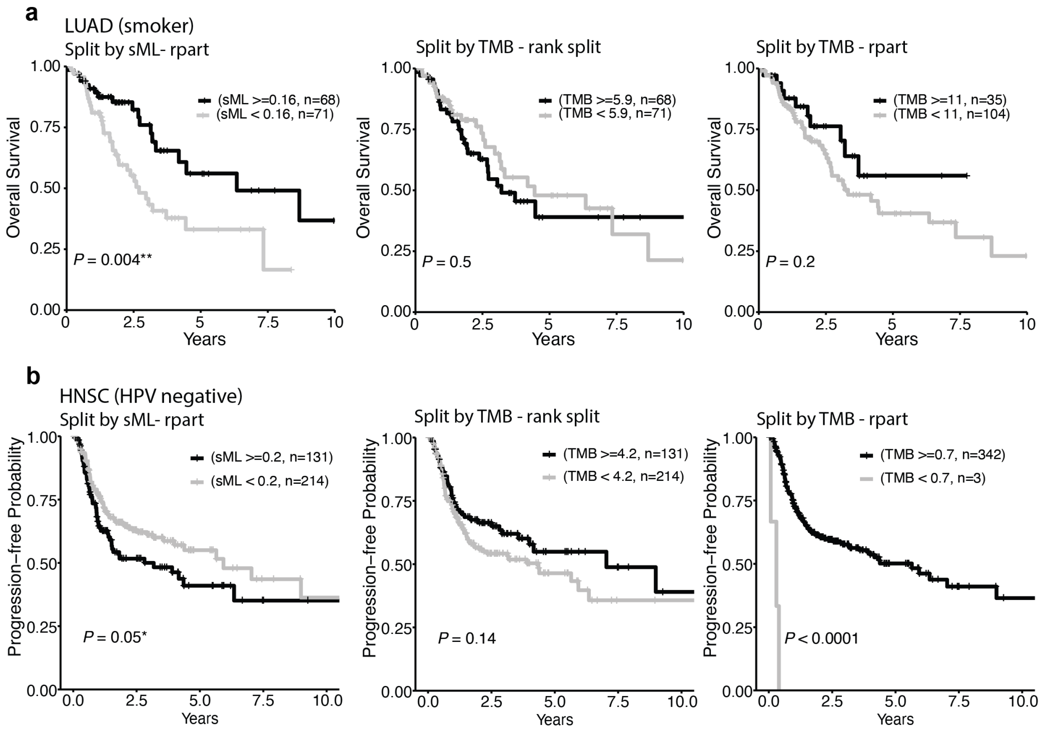

**Supplementary Fig. 14. TMB cannot replace sML in association with clinical outcomes.** (**a-b**) Kaplan-Meier curves of overall survival for LUAD (smoker) samples and progression-free survival for HNSC (HPV negative) samples, using either TMB or sML to stratify patients. Left panel shows results using sML. Middle and right panels show what happens when TMB is used to replace sML, using the sML-defined split or rpart split. Black lines indicate patients with high sML or TMB, and grey lines indicate patients with low sML or TMB. Log-rank test *P*-values between high- and low sML/TMB groups are shown.

#### 6. Shannon Index (SI) as an additional measure of intratumor heterogeneity

Subclonal diversity has previously been quantified through the number of clusters[13] or through entropy scores such as the Shannon Index (SI) , a population diversity metric that takes into account both cluster richness and cluster abundance. It can be calculated as $-\sum_{i=1}^{s} p_i \ln p_i$, where $s$ denotes the number of clusters present and $p_i$ denotes the proportion of mutations within a specific cluster to the total number of mutations in the tumor sample.

When there are 2 mutation clusters, i.e. one clonal and one subclonal, SI is a function of sML, as $-(1 - sML) \times \log(1 - sML) - sML \times \log(sML)$. As such, one sML value maps to one SI value, but one SI value can map back to two sML values (**Extended Data Fig. 7a**). Therefore, for cancer types that mostly present 2 mutation clusters as well as low sML, which tend to be high TMB (**Fig. 2d**), SI does not provide additional information to sML. In cancer types with more than 2 mutation clusters and reaching high sML, which tend to be low TMB (**Fig. 2d**, **Extended Data Fig. 7b**), SI presents further divergence at the same sML level. In these cancers, we hypothesize that using SI and sML together will further delineate different patterns of cancer evolution in association with clinical outcomes. Since the high collinearity between these two measures prevents us from finding optimal paired cutoffs using recursive partitioning, we only search for an optimal cutoff In SI recursively within patients with sML>0.5. We find that the addition of SI partitions high sML into further distinct survival groups for 11 cancer types, i.e., 5 cancers where high sML and low SI present the least favorable outcomes including uterine (endometrioid), stomach, renal papillary cell, thymoma, and pancreatic (log-rank test *P*-values < 0.05, **Extended Data Fig. 7c**), and 6 cancer types where high sML and low SI presents the most favorable outcomes, including prostate, kidney chromophobe, bladder, melanoma, HPV-negative head and neck, and thyroid cancers (log-rank test *P*-values < 0.05, **Extended Data Fig. 7d**). Among these cancer types, we find that TMB does not differ significantly between the high-sML high-SI group and the high-sML low SI group in more than 50% of them (and in inconsistent directions in the rest), suggesting that the observed survival differences are not merely due to differing TMB between these groups (**Supplementary Fig. 15**). Hence, we conclude that sML makes important and unique contributions to our understanding of cancer evolution, and in distinct manners for high-TMB cancers (it supersedes SI), and for low TMB cancers (it complements SI).

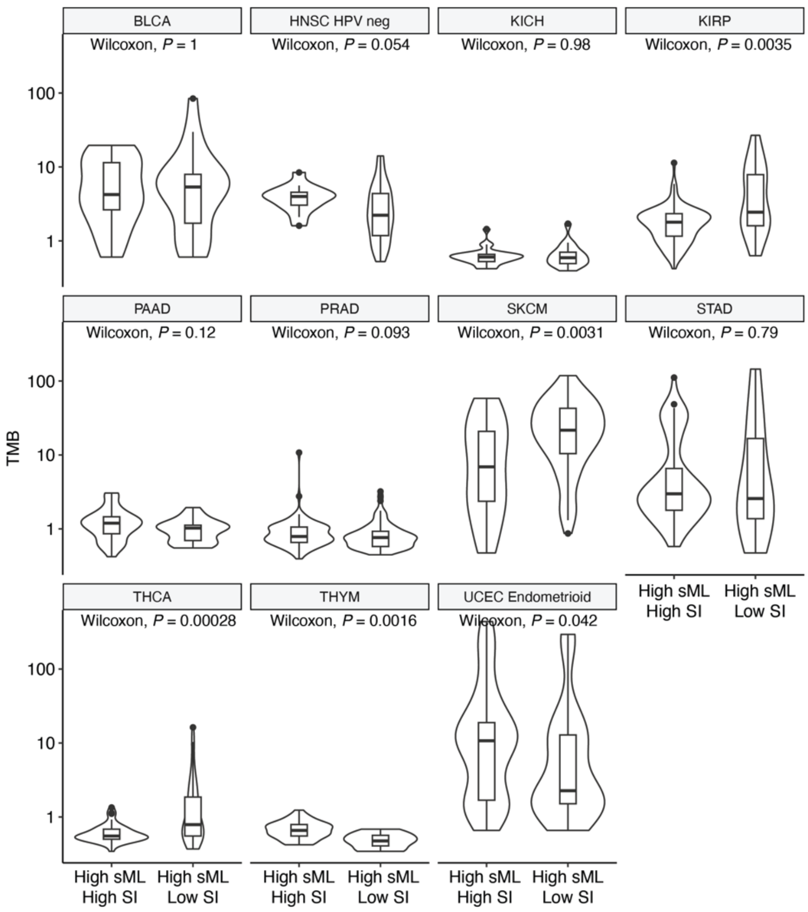

**Supplementary Fig. 15. Distributions of TMB between High-sML High SI and High-sML Low SI patient groups.** Box and violin plots comparing TMB between the high SI and low SI groups among high sML samples for the 11 cancer types with significant survival differences. Two-sided Wilcoxon rank sum test *P*-values comparing the distributions are shown.

### 7. Evaluation of confounders in sML association analysis in other clinical cohorts

#### 7.1 mCRPC clinical trial cohorts

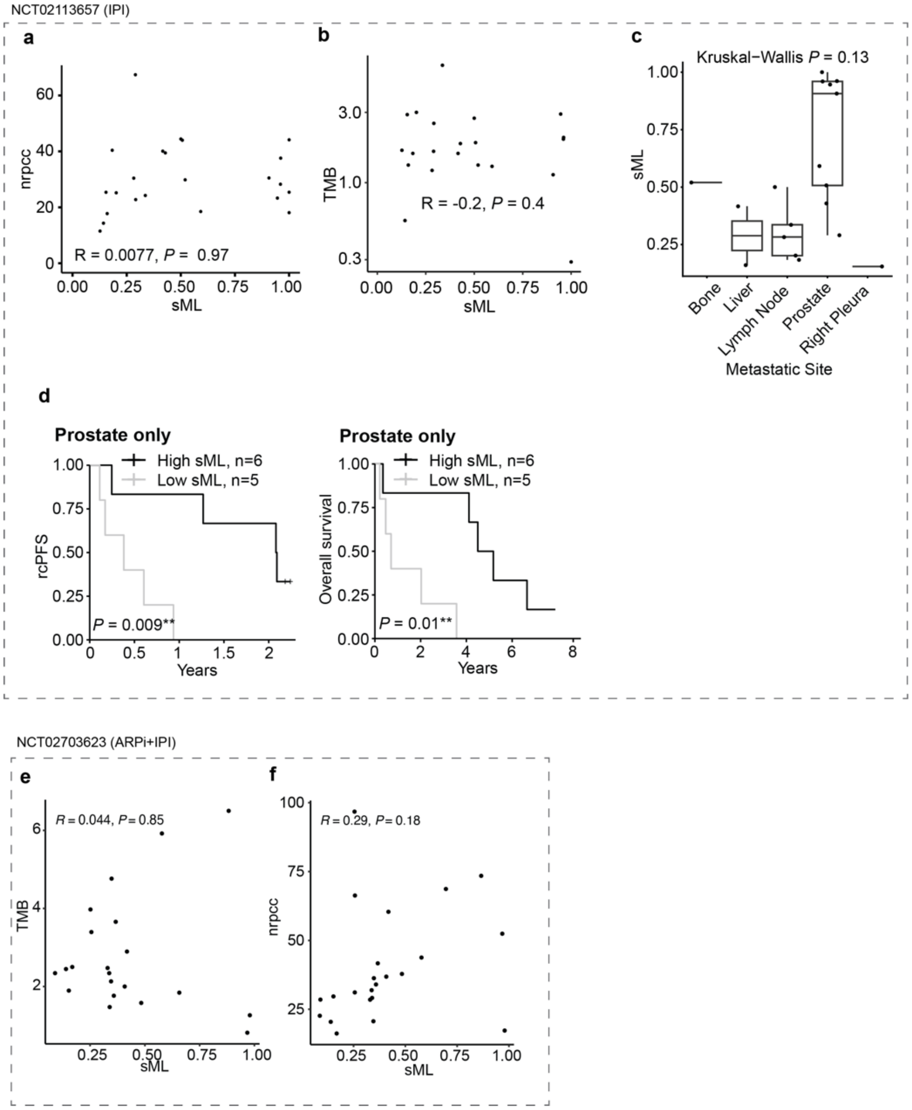

**Supplementary Fig. 16. Evaluation of confounders of sML analysis in mCRPC clinical trial cohorts, the ipilimumab (IPI) (NCT02113657) and the DynAMo (NCT02703623).** (**a**) Scatter plot with Pearson correlation coefficient and *P*-value between sML and nrpcc in trial NCT02113657. (**b**) Scatter plot with Pearson correlation coefficient and *P*-value between sML and TMB in trial NCT02113657. (**c**) Boxplots showing distribution of sML values by metastatic site among the 21 samples from NCT02113657, with Kruskal-Wallis test *P*-value shown. (**d**) Kaplan-Meier (KM) plot showing radiographic/clinical progression-free survival (rcPFS, left) and overall survival (OS, right) in patients stratified by sML, only comparing samples derived from prostate tissue, with log-rank test *P*-value shown. (**e**) Scatter plot with Pearson correlation coefficient and

*P*-value between sML and TMB in trial NCT02703623. (**f**) Scatter plot with Pearson correlation coefficient and *P*-value between sML and nrpcc in trial NCT02703623.

#### 7.2 OCCAMS esophageal adenocarcinoma samples

The WGS samples from the OCCAMS cohort provide the opportunity to compare sML to mutational signatures and survival outcomes. Calling of these signatures was done previously[7,15], and was possible in this cohort due to the much higher number of mutations captured over the whole genome. We find that mutation signatures do not differ between high and low sML **(Supplementary Fig. 17a)**. Additionally, we used recursive partitioning to identify the optimal cutoff between high and low signature proportion for each signature, following the same criteria used for sML (see **methods**). Of the 6 signatures tested[7,15] (SBS1, SBS2, SBS3, SBS17A, SBS17B, SBS18), only SBS17A shows a significant association with survival **(Supplementary Fig. 17b)**. However, this association does not persist after adjusting for covariates in a Cox proportional hazard model **(Supplementary Fig. 17c)**. For each signature, we fit a Cox model with age, gender, M stage, N stage, TMB, sML and the signature. For all models, we found the effect of sML was not impacted by the inclusion of the signature.

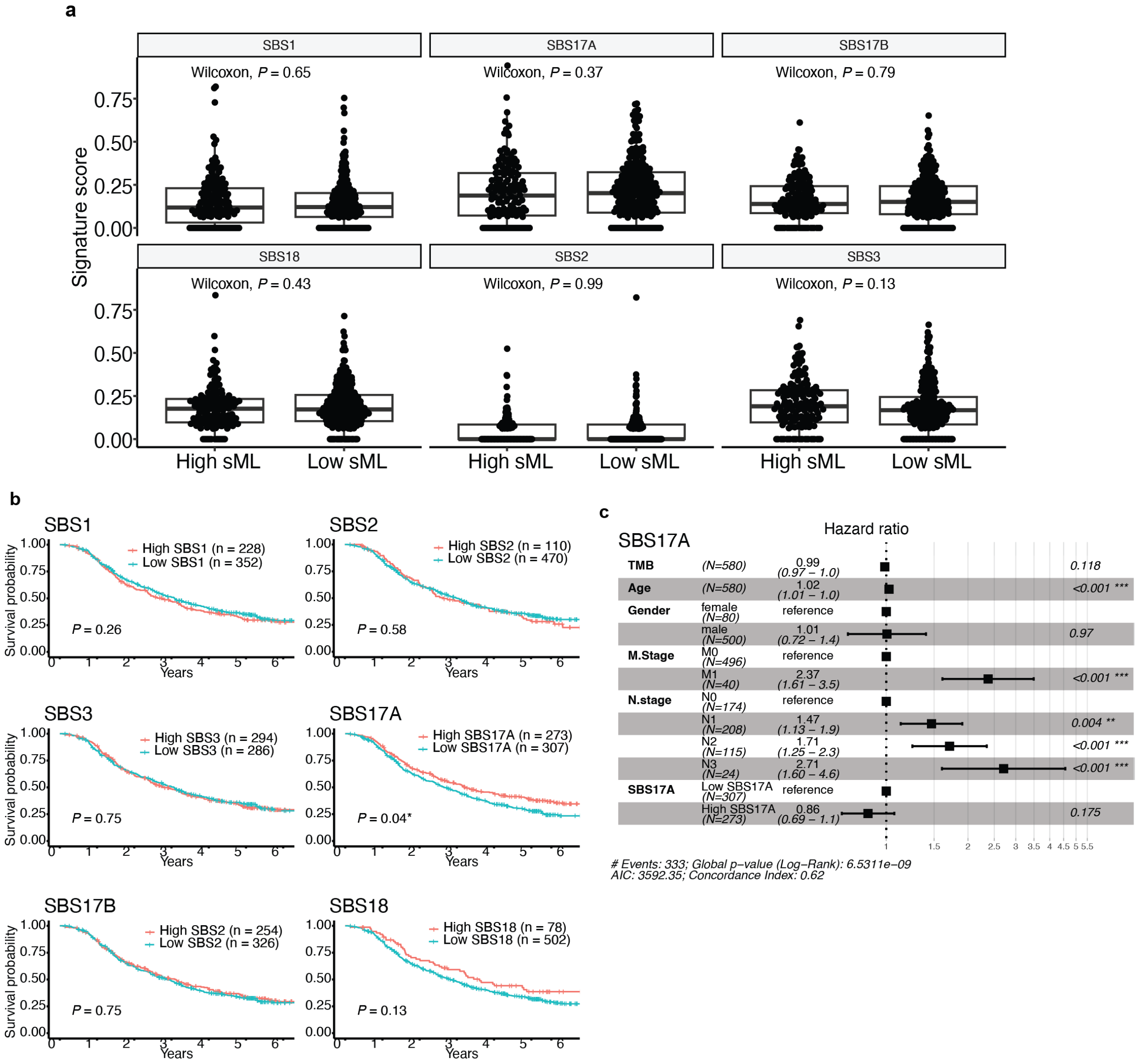

**Supplementary Fig. 17: Analysis of the OCCAMS WGS esophageal adenocarcinoma cohort.** **(a)** Boxplots with individual points overlayed showing no significant difference in signature scores for the previously reported esophageal cancer mutational signatures, with Wilcoxon rank-sum *P*-values shown. **(b)** KM plot showing overall survival between samples with high and low signature proportion for 6 signatures (SBS1, SBS2, SBS3, SBS17A, SBS17B, SBS18), with log-rank test *P*-values shown. **(c)** Forest plot showing hazard ratios and confidence intervals for Cox proportional hazard model with high vs. low SBS17A signature, adjusted for TMB, age, gender, M stage, and N stage.

### 8. Technical considerations for CliPP

*Notations:*

| Parameter | Definition |
|---|---|
| $\boldsymbol{\rho}$ | Tumor purity, takes value between 0 and 1 |
| $\boldsymbol{\beta}$ | Cancer cell fraction (CCF), takes value between 0 and 1 |
| $\boldsymbol{\phi}$ | Cellular prevalence (CP), is equal to $\boldsymbol{\rho\beta}$, takes value between 0 and 1 |
| $\boldsymbol{r_i}$ | Number of reads observed with variant alleles covering SNV $i$ |
| $\boldsymbol{n_i}$ | Total number of reads covering SNV $i$ |
| $\boldsymbol{c_i^T}$ | Total copy number for tumor cell covering SNV $i$ |
| $\boldsymbol{c_i^N}$ | Total copy number for normal cell covering SNV $i$ |
| $\boldsymbol{m_i^T}$ | Copy number of the major allele covering SNV $i$ |
| $\boldsymbol{b_i^V}$ | SNV-specific copy number covering SNV $i$, also known as multiplicity. |
| $\boldsymbol{\lambda}$ | Tuning parameter that controls the degree of the penalization |
| $\boldsymbol{S}$ | Total number of SNVs |
| $\boldsymbol{D}$ | Average read depth for the sample |
| $\boldsymbol{\theta}$ | Expected proportion of the variant allele |

#### 8.1 Computational structure using ADMM

The alternating direction method of multipliers (ADMM) is a well-established algorithm that addresses constrained optimization challenges by decomposing them into simpler subproblems.

To solve the optimization problem in Eq.(8) in the manuscript, we introduce auxiliary variables $\eta_{ij} = \omega_i - \omega_j$ for $i < j$ , thereby reformulating the problem as an equivalent constrained optimization problem:

$$S(\boldsymbol{\omega}, \boldsymbol{\eta}; \lambda) = -\hat{\ell}(\boldsymbol{\omega}) + \sum_{1 \le i < j \le S} p_\lambda(|\eta_{ij}|), \text{ subject to } \omega_i - \omega_j - \eta_{ij} = 0. \tag{S.1}$$

Using the augmented Lagrangian method, we further translate (S.1) into an unconstrained optimization problem and obtain the estimation of parameters by minimizing:

$$L(\boldsymbol{\omega}, \boldsymbol{\eta}, \boldsymbol{\tau}; \lambda) = S(\boldsymbol{\omega}, \boldsymbol{\eta}; \lambda) + \frac{\alpha}{2} \sum_{i<j} (\omega_i - \omega_j - \eta_{ij})^2 - \sum_{i<j} \tau_{ij}(\omega_i - \omega_j - \eta_{ij}), \tag{S.2}$$

where the dual variables $\boldsymbol{\tau} = \{\tau_{ij},\ i < j\}$ are Lagrange multipliers, and $\alpha$ is a penalty parameter and set to be 0.8 in our implementation. The estimates of $\boldsymbol{\omega}, \boldsymbol{\eta}$, and $\boldsymbol{\tau}$ are then computed iteratively by ADMM. The bottleneck in updating $\boldsymbol{\omega}$ at each ADMM iteration lies in the complex function $-\hat{\ell}(\boldsymbol{\omega})$. To address this challenge, we first use an estimated variance for the normal distribution and then employ a three-part piece-wise linear function to approximate each $g(\omega_i)$, leading to a quadratic approximation to $-\hat{\ell}(\boldsymbol{\omega})$ (see **Sections 8.2 and 8.3 for the full derivation**). It is noteworthy that when $\gamma > \frac{1}{\alpha} + 1$, the objective function $L(\boldsymbol{\omega}, \boldsymbol{\eta}, \boldsymbol{\tau};\ \lambda)$ based on the SCAD penalty is convex with respect to $\eta_{ij}$. Furthermore, the minimizer of $L(\boldsymbol{\omega}, \boldsymbol{\eta}, \boldsymbol{\tau};\ \lambda)$ with respect to $\eta_{ij}$ is unique and has a closed-form expression for given $(\boldsymbol{\omega},\ \boldsymbol{\tau})$. Therefore, updates of $\boldsymbol{\omega}, \boldsymbol{\eta}$, and $\boldsymbol{\tau}$ at one ADMM iteration proceed as follows. Given $\boldsymbol{\omega}^{(k)}$, $\boldsymbol{\tau}^{(k)}$ and $\boldsymbol{\eta}^{(k)}$ obtained from the $k$-th iteration, the update $\boldsymbol{\omega}^{(k+1)}$ is given by:

$$\boldsymbol{\omega}^{(k+1)} = \left(\boldsymbol{B}^T\boldsymbol{B} + \alpha\boldsymbol{\Delta}^T\boldsymbol{\Delta}\right)^{-1}[\alpha\boldsymbol{\Delta}^{\mathrm{T}}\left(\boldsymbol{\eta}^{(k)} - \boldsymbol{\tau}^{(k)}\right) - \boldsymbol{B}^T\boldsymbol{A}], \tag{S.3}$$

where $\boldsymbol{A}$ is a vector associated with $\boldsymbol{\omega}^{(k)}$, $\boldsymbol{B}$ is a matrix associated with $\boldsymbol{\omega}^{(k)}$ (see **Section 7.3 for the full derivation**), and $\boldsymbol{\Delta} = \{(e_i - e_j), i < j\}$ with $e_i$ defined as an $S \times 1$ vector whose $i$-th element is 1 and others are 0. For given $(\boldsymbol{\omega}^{(k+1)},\ \boldsymbol{\tau}^{(k)})$, $\eta_{ij}$ is updated by solving:

$$\hat{\eta}_{ij} = \operatorname{argmin}_{\eta_{ij}} \frac{\alpha}{2}\left(\delta_{ij} - \eta_{ij}\right)^2 + p_\lambda\left(|\eta_{ij}|\right),$$

where $\delta_{ij} = \omega_i^{(k+1)} - \omega_j^{(k+1)} - \alpha^{-1}\tau_{ij}^{(k)}$. This is a one-dimensional SCAD-penalized linear regression problem which has a closed-from solution. Thus, update of $\eta_{ij}$ at the $(k+1)$-th iteration for SCAD penalty with $\gamma > 1/\alpha + 1$ is given by Ma and Huang[16]

$$\hat{\eta}_{ij} = \begin{cases} ST\left(\delta_{ij}, \lambda/\alpha\right), & \text{if } \left|\delta_{ij}\right| \le \lambda + \lambda/\alpha, \\ \frac{ST(\delta_{ij}, \gamma\lambda/((\gamma-1)\alpha))}{1 - 1/((\gamma-1)\alpha)}, & \text{if } \lambda + \lambda/\alpha < \left|\delta_{ij}\right| \le \gamma\lambda, \\ \delta_{ij}, & \text{if } \left|\delta_{ij}\right| > \gamma\lambda \end{cases} \tag{S.4}$$

where $ST(x, t)$ is the soft thresholding operator defined as $\operatorname{sign}(x)(|x| - t)$ if $|x| \ge t$ and 0 otherwise. At last, we update $\boldsymbol{\tau}$ by:

$$\boldsymbol{\tau}^{(k+1)} = \boldsymbol{\tau}^{(k)} - \alpha\left(\boldsymbol{\Delta}\boldsymbol{\omega}^{(k+1)} - \boldsymbol{\eta}^{(k+1)}\right). \tag{S.5}$$

Consequently, the ADMM algorithm to solve the proposed model can be summarized as follows: (i) Initialize the estimates. Normally, a random initialization works well. We initialize $\omega_i^{(0)} = \log\frac{r_i/n_i}{1 - r_i/n_i}$, $\eta_{ij}^{(0)} = \omega_i^{(0)} - \omega_j^{(0)}$, and $\boldsymbol{\tau}^{(0)} = \mathbf{1}_S$, where $1_S$ is defined as a vector of 1's with length $S$. (ii) At the $(k+1)$-th iteration, compute $\left(\boldsymbol{\omega}^{(k+1)},\ \boldsymbol{\eta}^{(k+1)},\ \boldsymbol{\tau}^{(k+1)}\right)$ as described above. (iii) Terminate the algorithm if a stopping criterion (e.g., $\max\left(\eta_{ij}^{(k+1)} - \eta_{ij}^{(k)}\right) < 0.01$) is met . Otherwise, go back to step (ii) and repeat.

#### 8.2 Brief induction of the linear approximation

To addressing the challenge in updating $\boldsymbol{\omega}$, we adopt a three-part piece-wise linear function to approximate $g(\omega_i)$ in the negative log-likelihood in Eq.(8) in the Method section. Shown on the right is an example of such an approximation. Specifically, we assume the linear approximation takes the form of $g(\omega_i) \approx u_i + v_i\omega_i$. Recall in the manuscript, we already have:

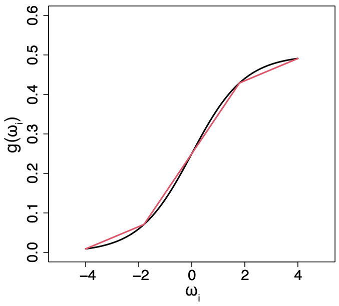

$$\theta_i = g(\omega_i) = \frac{b_i^V e^{\omega_i}}{(1+e^{\omega_i})A}$$

where $A = 2 - 2\rho + c_i^V\rho$ is a constant. Let's set

$$f(x) = \frac{b_i^V e^x}{(1+e^x)A}$$

$$g(x) = \begin{cases} a_1x + b_1, & \text{if } x < x_1 \\ a_2x + b_2, & \text{if } x \geq x_2 \\ a_3x + b_3, & \text{if } x_1 \leq x \leq x_2 \end{cases}$$

where $g(x)$ is used to approximate $f(x)$. We define the approximation error $D$ as follows:

$$D = \sup_{x\in R} |f(x) - g(x)|.$$

Then our goal is to find a pair of separation points $(x_1, x_2)$ for constructing $g(x)$ that achieves the smallest $D$. To that end, we employ a grid search strategy and limit the range of $x$ to $[-M, M]$ to save computational cost. In practice, we set $\mathrm{M} = 4$ which can roughly recover the full dynamic range of $f(x)$. The grid spans from $-M$ to $M$ with a step size of 0.1, and we record the approximation error $D$ at each pair of separation points $(x_1, x_2)$. Finally, the approximation function is designed using the pair of $(x_1, x_2)$ associated with the smallest $D$.

#### 8.3 Computational details

To estimate $\boldsymbol{\omega}, \boldsymbol{\eta}, \boldsymbol{\tau}$, we designed an ADMM-based algorithm for minimizing the objective function in Eq.(S.2). Given $(\boldsymbol{\eta}, \boldsymbol{\tau})$, an update of $\boldsymbol{\omega}$ can be derived by setting $\partial L(\boldsymbol{\omega}, \boldsymbol{\eta}, \boldsymbol{\tau}\,;\,\lambda)/\partial\boldsymbol{\omega} = 0$. However, it is difficult to directly compute the derivative. Here, we utilize two strategies. First, instead of using the theoretic variance, we use an estimated variance based on the estimated $\boldsymbol{\omega}$ from the previous step. Second, we approximate $g(\omega_i)$ by a linear function as discussed above in Section 7.2. That is, at step $k+1$, we rewrite the objective function as follows:

$$L(\boldsymbol{\omega}, \boldsymbol{\eta}, \boldsymbol{\tau}; \lambda, \boldsymbol{\omega}^{(k)}) = S(\boldsymbol{\omega}, \boldsymbol{\eta}; \lambda, \boldsymbol{\omega}^{(k)}) + \frac{\alpha}{2}\sum_{i<j}(\omega_i - \omega_j - \eta_{ij})^2 - \sum_{i<j}\tau_{ij}(\omega_i - \omega_j - \eta_{ij})$$

$$= \sum_{\mathrm{i}=1}^{S}\{-\log\frac{1}{\sqrt{2\pi n_i g(\omega_i)(1-g(\omega_i))}} + \frac{(n_i g(\omega_i)-\mathrm{r_i})^2}{2n_i g(\omega_i)(1-g(\omega_i))}\} + \sum_{i\le i<j\le S} p_\lambda(|\eta_{ij}|)$$

$$+\frac{\alpha}{2}\sum_{i<j}(\omega_i-\omega_j-\eta_{ij})^2 - \sum_{i<j}\tau_{ij}(\omega_i-\omega_j-\eta_{ij})$$

$$\approx \frac{1}{2}\sum_{i=1}^{S}\frac{n_i\left(g(\omega_i)-\frac{r_i}{n_i}\right)^2}{g\left(\omega_i^{(k)}\right)\left(1-g\left(\omega_i^{(k)}\right)\right)} + \sum_{i\le i<j\le S} p_\lambda(|\eta_{ij}|)$$

$$+\frac{\alpha}{2}\sum_{i<j}(\omega_i-\omega_j-\eta_{ij})^2 - \sum_{i<j}\tau_{ij}(\omega_i-\omega_j-\eta_{ij}) + C_0$$

$$= \frac{1}{2}\sum_{i=1}^{S}\frac{n_i\left(u_i+v_i\omega_i-\frac{r_i}{n_i}\right)^2}{g\left(\omega_i^{(k)}\right)\left(1-g\left(\omega_i^{(k)}\right)\right)} + \sum_{i\le i<j\le S} p_\lambda(|\eta_{ij}|)$$

$$+\frac{\alpha}{2}\sum_{i<j}(\omega_i-\omega_j-\eta_{ij})^2 - \sum_{i<j}\tau_{ij}(\omega_i-\omega_j-\eta_{ij}) + C_0$$

$$= \frac{1}{2}\sum_{i=1}^{S}(A_i+B_i\omega_i)^2 + \frac{\alpha}{2}\sum_{i<j}\left((e_i-e_j)^T\boldsymbol{\omega}-\eta_{ij}-\alpha^{-1}\boldsymbol{\tau}\right)^2 + C$$

$$= \frac{1}{2}\left|\left|\boldsymbol{A}+\boldsymbol{B\omega}\right|\right|^2 + \frac{\alpha_k}{2}\left|\left|\Delta\boldsymbol{\omega}-\boldsymbol{\eta}-\alpha_k^{-1}\boldsymbol{\tau}\right|\right| + C$$

where $A_i = \frac{\sqrt{n_i}(u_i-r_i/n_i)}{\sqrt{g(\omega_i^{(k)})(1-g(\omega_i^{(k)}))}}$, and $B_i = \frac{\sqrt{n_i}v_i}{\sqrt{g(\omega_i^{(k)})(1-g(\omega_i^{(k)}))}}$, $C$ is a function independent of $\boldsymbol{\omega}$, $e_i$ is $S\times 1$ vector whose $i^{th}$ element is 1 and others are 0, and $\Delta = \left\{(e_i-e_j\ ), i<j\right\}^T$, A is a vector given by $\boldsymbol{A} = \{A_1,\ldots,A_S\}^T$, and $\boldsymbol{B}$ is a matrix defined as

$$\boldsymbol{B} = \begin{bmatrix} B_1 & 0 & \cdots & 0 \\ 0 & B_2 & \cdots & 0 \\ \vdots & \vdots & \ddots & \vdots \\ 0 & \cdots & 0 & B_N \end{bmatrix}$$

Therefore,

$$\boldsymbol{\omega}^{(k+1)} = \left(\boldsymbol{B}^T\boldsymbol{B}+\alpha\boldsymbol{\Delta}^T\boldsymbol{\Delta}\right)^{-1}\left[\alpha\boldsymbol{\Delta}^{\mathrm{T}}\left(\boldsymbol{\eta}^{(k)}-\boldsymbol{\tau}^{(k)}\right)-\boldsymbol{B}^T\boldsymbol{A}\right].$$

The direct inversion of matrix $\boldsymbol{B}^T\boldsymbol{B}\ +\alpha\Delta^T\Delta$ may not be feasible when $S$ is large, but fortunately we do not need to perform a $N\times N$ matrix inversion, as Miller[17] suggested. Notice that $\Delta^{\boldsymbol{T}}\Delta = N\boldsymbol{I}_S - \mathbf{1}_S\mathbf{1}_{\boldsymbol{S}}^T$, thus,

$$\boldsymbol{B}^T\boldsymbol{B}+\alpha\Delta^T\Delta = (\boldsymbol{B}^T\boldsymbol{B}+\boldsymbol{S}\alpha\mathbf{1}_S) - \alpha\mathbf{1}_S\mathbf{1}_S^T.$$

Now let $\boldsymbol{M} = \boldsymbol{B}^T\boldsymbol{B} + \boldsymbol{S}\alpha\mathbf{1}_S$, and $\boldsymbol{R} = -\alpha\mathbf{1}_S\mathbf{1}_S^{\boldsymbol{T}}$. $\boldsymbol{M}$ is easy to invert since it is a diagonal matrix with non-zero diagonal elements, and $rank(\boldsymbol{R}) = 1$. Then we have:

$$\left(\boldsymbol{B}^T\boldsymbol{B} + \alpha\boldsymbol{\Delta}^{\boldsymbol{T}}\boldsymbol{\Delta}\right)^{-1} = (\boldsymbol{M} + \boldsymbol{R})^{-1} = \boldsymbol{M}^{-1} - \frac{1}{1+g}\boldsymbol{M}^{-1}\boldsymbol{R}\boldsymbol{M}^{-1}$$

where $g = trace(\boldsymbol{R}\boldsymbol{M}^{-1})$. Here we have avoided computing the inverse of an $S \times S$ matrix, instead, we only need to compute a series of matrix multiplications. The updates of $\boldsymbol{\eta}$ and $\boldsymbol{\tau}$ are achieved by Eq.(S.4) and Eq.(S.5), respectively, as discussed in **Section 8.1**.

#### 8.4 Post-processing steps

*Post-processing of the clustering output.* While the $\phi_i'$s provide a natural definition of tumor subclonal architecture, manual post-processing are necessary in practice[1,18]. Commonly seen with penalized likelihood-based approach, for example, are spurious clusters containing a negligible number of SNVs ($< 5$ SNVs or $< 1\%$ of the total SNVs. Therefore in CliPP, we implement several post-filtering steps to alleviate the need for manual curation for the following scenarios: (1) The presence of superclusters, which are clusters with an estimated $CCF \gg 1$. This occurrence often correlates with errors in CNA estimates taken as input by CliPP. (2) The presence of insignificant resulting clones, including (a) when the sample has > 2 clusters and the current proportion of clonal mutation $\leq 0.15$; (b) small gaps between CP values of any two clusters, i.e., when CP values between any two neighboring clusters $< 0.1$; and (c) when the number of mutations within a subclone is less than $5\%$ of the total number of mutations in a given sample. CliPP responds to each of these scenarios by merging the affected cluster with its nearest neighboring cluster. The current choice of cutoffs in these filters was trained with a sensitivity analysis using the PCAWG WGS data. These steps can be further modified by users when applying CliPP to a new dataset.

*Down-sampling of SNVs in hypermutated tumor samples.* While samples with fewer than 50,000 SNVs can typically be processed using approximately 256 GB of memory, a lower threshold of 30,000 to 35,000 SNVs is recommended to optimize memory efficiency. For hyper-mutated samples exceeding this range, we adopt a down-sampling strategy in which 30,000 to 35,000 SNVs are randomly sampled to ensure that the resulting matrices remain within feasible computational limits. To maximize SNV coverage, this random sampling is performed 10 times, and the CliPP clustering results are integrated using kernel smoothing.

#### 8.5 Advancements in CliPP1.3.3

The advancements of CliPP1.3.3 compared to the earlier developing version which was used in the PCAWG study in Dentro et al. 2021[1] are summarized as follows:

1. **Mathematical Corrections:**
    - Corrected the computation of multiplicity.
    - Revised the penalized likelihood formulation.
    - Improved hyperparameter selection strategy.
2. **Coding Improvements:**
    - Completely re-designed both the pre- and post-processing pipelines.
    - Addressed reported bugs that caused incorrect clustering results on some SNVs
3. **Performance Enhancements:**
    - Optimized the code to further speed up over 10x.
4. **User Experience Improvements:**
    - Easier installation via GitHub and Docker.
    - Encapsulated the entire pipeline into a single step with flexible parameter modifications.
5. **New GUI tool:**
    - Developed an R Shiny app for easy CliPP implementation without any need for programming.
